# Free water elimination tractometry for aging brains

**DOI:** 10.1101/2024.11.10.622861

**Authors:** Kelly Chang, Luke Burke, Nina LaPiana, Bradley Howlett, David Hunt, Margaret Dezelar, Jalal B. Andre, Patti Curl, James D. Ralston, Ariel Rokem, Christine L. MacDonald

## Abstract

Tractometry of diffusion-weighted magnetic resonance imaging (dMRI) non-invasively quantifies tissue properties of brain connections. It is widely used in aging studies but could be less reliable in aging brains due to increased white matter free water. We demonstrate that computational free water elimination (FWE) increases reliability and accuracy of tractometry in a large (n = 339) cohort of older adults (66 – 103 y.o.). We found substantial (up to ∼37%) improvements in reliability in a split-half comparison at every stage of the pipeline: estimation of voxel-level fiber orientation distribution functions, delineation of major pathway trajectories, and assessment of tissue properties along the pathways. FWE also improves inferences from tractometry, producing more accurate cross-validated predictions of clinician Fazekas scores. By sub-sampling a multi-b-value dataset, we demonstrated that these findings generalize to both single-b-value data, which is important for many datasets where only one b-value may be available. Overall, the results highlight the importance of accounting for free water in tractometry studies, especially in aging brains. We provide opensource software for free-water elimination that can be applied to a wide range of clinical and research datasets (https://github.com/nrdg/fwe).

## 1 Introduction

Tractometry based on diffusion MRI (dMRI) measurements provides accurate and reliable assessments of white matter tissue properties along the length of major white matter pathways, which can be used to make inferences about brain connections (Kruper et al., 2021; Takemura et al., 2024). Tractometry processing pipelines include several steps: modeling of individual-voxel fiber orientation distribution functions (fODF), computational tractography (Jeurissen et al., 2019), delineation of major white matter pathways (Yeatman et al., 2012), modeling of individual voxel tissue properties, and the projection of these tissue properties onto the length of each of the tracts. These so-called “tract profiles” are then used as the input to statistical analysis and inference.

Tractometry is particularly useful in assessing brain aging as brain connectivity is profoundly affected by aging, and the cognitive effects of brain disconnection in aging may be mediated by changes to brain white matter microstructure (Cox et al., 2016). Areas of leukoaraiosis, apparent as white matter hyperintensites (WMH) in fluid attenuated inversion recovery (FLAIR) imaging, may be particularly interesting to investigate with tractometry in aging subjects, because they are thought to be indicative of ischemic brain injury and related to its impact on cognition (Wardlaw et al., 2015). However, these are also regions in which the directional specificity of the diffusion signal along the trajectory of white matter fiber bundles is potentially confounded by the effects of excess cerebrospinal fluid (CSF) that may have infiltrated the white matter (Chang et al., 2023).

Computational free water elimination (FWE) provides a potential solution, because it removes the confounding effect of free water on the diffusion signal (Hoy et al., 2014; Pasternak et al., 2009; Pierpaoli & Jones, 2004). Previous work has shown the benefits of this approach in improving tractography estimates of major white matter pathways (Hoy et al., 2015) and in tractography around regions of brain white matter edema (e.g., around tumors; Ould Ismail et al., 2019).

Here, we studied the impact of free water elimination on tractometry in multi-shell high angular resolution dMRI measurements in a large (n = 339) group of aging (ages 66 - 103 y.o.) participants from the Adult Changes in Thought (ACT) Study, a well-characterized, community-based, longitudinal, prospective cohort study (Kukull et al., 2002). FWE was conducted using a free water diffusion tensor imaging (FWDTI) model (Golub et al., 2021; Henriques et al., 2017; Hoy et al., 2014; Pasternak et al., 2009). A split-half approach showed improvements in reliability at every step of the tractometry pipeline. We found that these improvements in reliability also translated to improved accuracy in classification of these subjects into different white matter lesion categories (Fazekas et al., 1987). The improvements in inference from FWE tractometry translate to single-shell dMRI measurements, which are more common in clinical research settings, even while improvements in reliability are more modest. Taken together, these results demonstrate that FWE could provide a benefit in a wide range of studies that use dMRI to understand brain connections in aging individuals.

## 2 Methods

### 2.1 Participants

A total of 339 individuals underwent a research-grade MRI scan which was leveraged for this analysis. Participants were a part of the Adult Changes in Thought (ACT) study, a longitudinal cohort study embedded in an integrated healthcare delivery system. The overall goal of the ACT study is to understand factors that contribute to Alzheimer’s Disease and related dementia. The ACT study prospectively and retrospectively collects data on participants through biannual assessments, electronic medical research abstraction, and through ancillary activities such as research-grade MRI scans, optical coherence tomography scans, and collection of biofluids for genetic and biomarker analysis (Kukull et al., 2002). The institutional review board at the Kaiser Permanente Research Institute approved the study protocol.

Participants were excluded from the analysis if their average neighboring dMRI correlation (NDC; Yeh et al., 2019) – a data quality metric, which we have previously shown to be closely related to expert observers (Richie-Halford et al., 2022) – fell 2 standard deviations below the sample average. This resulted in 330 participants (ages 66 - 103, mean age = 79.52; 176 females) remaining in the sample.

### 2.2 MRI Acquisitions

Data were acquired at the University of Washington at the Diagnostic Imaging Sciences Center (DISC) on a 3T Philips Ingenia Elition MRI scanner with a 32-channel head coil. The current analysis leveraged the T1-weighted (T1w) and fluid attenuated inversion recovery (FLAIR) whole-brain structural images and two diffusion-weighted acquisitions of opposite phase encoding directions from each participant.

#### 2.2.1 T1w

3D MPRAGE T1w images were acquired at 1 mm^3^ isotropic resolution (TR = 6.5 ms, TE = 2.9 ms, flip angle = 9°, FOV = 256 × 256 mm, matrix size = 256 × 256, 211 sagittal slices).

#### 2.2.2 FLAIR

3D FLAIR images were acquired with an inversion recovery sequence at 1 mm^3^ isotropic (TR = 5000, TE = 291 ms, TI = 1800 ms, flip angle = 90°, FOV = 256 × 256 mm, matrix size = 256 × 256, 176 sagittal slices).

#### 2.2.3 Diffusion-weighted imaging

Diffusion-weighted images were acquired with a spin-echo echo-planar imaging sequence at an in-plane spatial resolution of 1.8128 × 1.8125 mm^2^ (TR = 3500 ms, TE = 89 ms, FOV = 232 × 232 mm, matrix size = 128 × 128, slice thickness = 2.20 mm, slice gap = 0.22 mm, 57 axial slices). The dMRI data were collected at 3 *b*-values, 500, 1000, 2500 s/mm^2^, with 32, 64, and 128 directions, respectively. Twenty-six non-diffusion-weighted (*b* = 0) measurements were interleaved with the diffusion-weighted measurements. Two diffusion-weighted images with opposite phase-encoding at all directions were acquired for each participant.

### 2.3 Preprocessing

#### 2.3.1 T1w

T1w images were preprocessed using the QSIprep-0.18.1 (Cieslak et al., 2021) anatomical pipeline. In brief, the T1w image was corrected for intensity non-uniformity (INU) using N4BiasFieldCorrection (ANTs-2.4.3; Tustison et al., 2010), and used as an anatomical reference throughout the workflow. The T1w image was reoriented into AC-PC alignment via a 6-DOF transform extracted from a full affine registration to the MNI152NLin2009cAsym template (Fonov et al., 2009). A nonlinear registration to the T1w image from AC-PC space was estimated via symmetric nonlinear registration (SyN) using antsRegistration (ANTs-2.4.3; Avants et al., 2008). Brain extraction was performed on the T1w image using SynthStrip (Hoopes et al., 2022), and automated segmentation was performed using SynthSeg (Billot, Greve, et al., 2023; Billot, Magdamo, et al., 2023) from FreeSurfer-7.3.1.

#### 2.3.2 FLAIR

First, FLAIR images were skull-stripped with mri_synthstrip (Hoopes et al., 2022) and corrected for intensity non-uniformity with the N4 algorithm (Tustison et al., 2010) implemented in ANTsPy. Images were then coregistered to the participant’s preprocessed T1w image from Section 2.3.1 through a rigid body registration. FLAIR images were standardized by participants’ white matter voxel intensities based on the individual white matter masks created in Section 2.3.1. Lastly, nonlinear spatial normalization to the MNI152NLin2009cAsym template was performed on the FLAIR images.

#### 2.3.3 Diffusion MRI

Diffusion MRI preprocessing was performed using the QSIprep-0.18.1 (Cieslak et al., 2021) diffusion pipeline: Diffusion images were processed with MP-PGCA denoising (MRtrix3’s dwidenoise, 5 voxel window; Veraart et al., 2016) and Gibbs ringing correction (MRtrix3’s mrdegibbs; Kellner et al., 2016). The mean intensity of the dMRI series was adjusted so all the mean intensity of *b* = 0 images matched across each dMRI scanning sequence. Diffusion images were grouped by their phase-encoded polarity and merged into a single file. FSL-6.0.5.1 eddy was used for head motion correction and Eddy current correction (*q*-space smoothing factor = 10, 5 iterations, 1000 voxels; Andersson & Sotiropoulos, 2016). A linear first- and second-level models were used to characterize Eddy current related spatial distortion. *q*-space coordinates were forcefully assigned to shells and field offset was attempted to be separated from participant movement. Shells were aligned post-eddy correction. Eddy’s outlier replacement was run by grouping data by slice, only including values from slices determined to contain at least 250 intracerebral voxels. Groups deviating by more than 4 standard deviations from the prediction had their data replaced with imputed values. Data was collected with reversed phase-encode blips, resulting in pairs of images with distortions going in opposite directions. Multiple dMRI series were acquired with opposite phase encoding directions, so *b* = 0 images were extracted from each to be used for diffeomorphic registration (Irfanoglu et al., 2015). The susceptibility-induced off-resonance field was estimated from these pairs using a method similar to that described in Andersson et al. (2003). The field maps were ultimately incorporated into the Eddy current and head motion correction interpolation. Final interpolation was performed using the jac method. dMRI time series were resampled, generating a preprocessed dMRI run with 1.8 × 1.8 × 1.8 *mm*^3^ isotropic voxels. Lastly, B1 field inhomogeneity correction was applied to the resampled images (MRtrix3’s dwibiascorrect with N4 algorithm; Tustison et al., 2010).

### 2.4 Fazekas Scores

Fazekas scores are a visual assessment score of WMH burden in the brain (Fazekas et al., 1987). Subscores range from 0 (absence of WMH) to 3 (abundant WMH) and are assessed for periventricular and deep white matter hyperintensity separately. A final Fazekas score is assessed as the total of the periventricular and deep Fazekas scores (range of 0 - 6). Fazekas scores were rated for each participant FLAIR image by one of two board-certified radiologists (authors J.B.A. and P.C.).

### 2.5 FLAIR White Matter Hyperintensity Segmentation and Categorization

We used a convolutional neural network (HyperMapper; Mojiri Forooshani et al., 2022) to automatically segment WMH voxels from FLAIR images. HyperMapper generated probabilistic voxel-wise maps of WMH from the spatially normalized T1w, T2w, and FLAIR images. Voxels with probabilities ≥ 0.5 were defined as WMH voxels. Separate WMH regions of interest (ROIs) were defined as contiguous voxels with scikit-image (van der Walt et al., 2014). The WMH ROIs were transformed from the MNI152NLin2009cAsym template space to the space of the individuals and used for the remaining analyses.

Next, we classified the WMH ROIs based on its location in the white matter. WMH ROIs were categorized as periventricular if the ROI was adjacent to the lateral ventricles – excluding voxels immediately adjacent (within 1 mm) to the ventricles to avoid partial volume effects – or as deep otherwise. The remaining white matter volume was categorized as normal-appearing white matter (NAWM).

### 2.6 Creating a Single-Shell Diffusion Dataset

We created a single-shell diffusion-weighted dataset by subsampling the preprocessed multi-shell diffusion-weighted measurements to only include non-diffusion (*b* = 0, 26 measurements) and *b*-value of 1000 s/mm^2^ (64 directions) images. We define the complete multi-shell dataset as the “full multi-shell” data. Correspondingly, we define the *b* = 1000 sub-sampled dataset as the “full single-shell” dataset.

### 2.7 Free water Elimination

Free water is modeled in each voxel in the white matter using a mixture model (Pierpaoli & Jones, 2004):

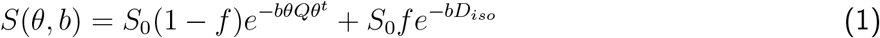

where *S*(*θ, b*) is the signal measured when a gradient is applied in direction *θ* (encoded as a unit vector) with diffusion-weighting *b, S*_0_ is the signal measured when no diffusion-weighting is applied, *Q* is a symmetric matrix representing a diffusion tensor, which has six free parameters, and *D*_*iso*_ = 3*mm*^2^*/s* is the diffusivity of free water at body temperature, and *f* is a free parameter for the free water fraction in the voxel. This is the free water DTI model (FWDTI).

FWDTI is well-posed for multi-shell acquisitions, and can be fit accurately using non-linear least squares optimization (Hoy et al., 2014). We used the implementation previously described in (Henriques et al., 2017), which is included in the open-source DIPY software library (Garyfallidis et al., 2014). In single-shell acquisitions, the model is ill-posed and a spatially-regularized gradient descent algorithm is used to fit the model (Pasternak et al., 2009). We used the software implementation from (Golub et al., 2021). After the FWDTI model is fit, the free water component can be eliminated:

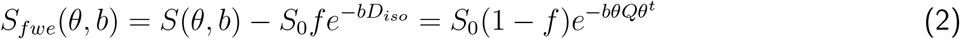

where FWE stands for “free water eliminated” and *S*_*fwe*_ is the signal with the free water component removed.

### 2.8 Within-Participant Split-Half Datasets

To examine the impact of free water elimination on reliability, we created within-participant split-half datasets from the full multi-shell and single-shell datasets. The dMRI images were randomly and evenly split into two at each *b*-value.

For the multi-shell dataset, this resulted in two dMRI images with 3 *b*-values, 500, 1000, 2500 s/mm^2^, with 16, 32, and 64 directions, respectively, and 13 non-diffusion-weighted (*b* = 0) measurements for each participant. We define this dataset as the “split-half multi-shell” dataset.

For the single-shell dataset, this resulted in two dMRI images with 32 directions at *b*-value of 1000 s/mm^2^ and 13 non-diffusion-weighted (*b* =0) measurements for each participant. We define this dataset as the”split-half single-shell” dataset. Subsequent analyses were performed on the full and split-half, multi-shell and single-shell datasets.

### 2.9 Tractometry

We used pyAFQ (Kruper et al., 2021), an open-source automated tractometry pipeline. The methods were previously described in detail in Kruper et al. (2021); briefly: We used methods from DIPY (Gary-fallidis et al., 2014) to perform constrained spherical deconvolution (CSD; Tournier et al., 2007) using all of the *b*-value shells in the dMRI acquisition, and we used the fODFs in every voxel as cues for probabilistic tractography, with 8 streamlines seeded in every voxel in the white matter. Every individual’s brain was aligned to the MNI152NLin2009cAsym template using non-linear registration, implemented in DIPY. For every one of 28 major white matter pathways (Table 1), inclusion and exclusion ROIs defined in the space of the template were back-transformed into the space of the individual and used to select streamlines that passed through inclusion and did not pass through exclusion ROIs. Each streamline was resampled to 100 points. Streamlines that deviated significantly (*>* 5 standard deviations) from the median trajectory were removed from each tract.

**Table 1:**
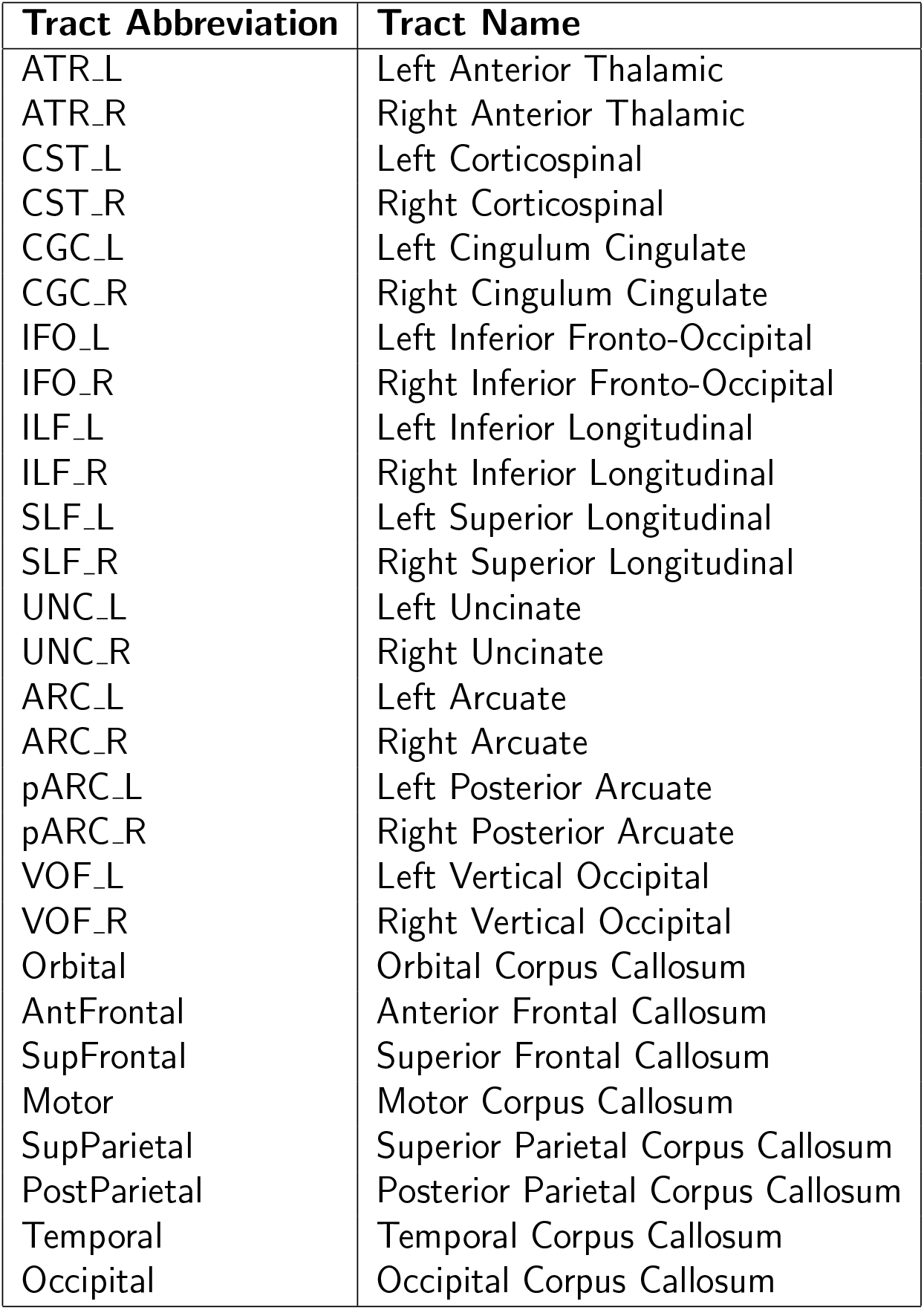
Tract names and their corresponding abbreviations. Subsections of the Corpus Callosum based on Dougherty et al. (2007).

| Tract Abbreviation | Tract Name |
| --- | --- |
| ATR_L | Left Anterior Thalamic |
| ATR_R | Right Anterior Thalamic |
| CST_L | Left Corticospinal |
| CST_R | Right Corticospinal |
| CGC_L | Left Cingulum Cingulate |
| CGC_R | Right Cingulum Cingulate |
| IFO_L | Left Inferior Fronto-Occipital |
| IFO_R | Right Inferior Fronto-Occipital |
| ILF_L | Left Inferior Longitudinal |
| ILF_R | Right Inferior Longitudinal |
| SLF_L | Left Superior Longitudinal |
| SLF_R | Right Superior Longitudinal |
| UNC_L | Left Uncinate |
| UNC_R | Right Uncinate |
| ARC_L | Left Arcuate |
| ARC_R | Right Arcuate |
| pARC_L | Left Posterior Arcuate |
| pARC_R | Right Posterior Arcuate |
| VOF_L | Left Vertical Occipital |
| VOF_R | Right Vertical Occipital |
| Orbital | Orbital Corpus Callosum |
| AntFrontal | Anterior Frontal Callosum |
| SupFrontal | Superior Frontal Callosum |
| Motor | Motor Corpus Callosum |
| SupParietal | Superior Parietal Corpus Callosum |
| PostParietal | Posterior Parietal Corpus Callosum |
| Temporal | Temporal Corpus Callosum |
| Occipital | Occipital Corpus Callosum |

To characterize microstructural tissue properties within each voxel of the white matter, we used the diffusional kurtosis imaging model (DKI; Jensen et al., 2005). DKI was fit on multi-shell datasets using the Diffusion Imaging in Python (DIPY) software library (Garyfallidis et al., 2014; Henriques et al., 2017, 2021). We derived fractional anisotropy (DKI-FA), mean diffusivity (DKI-MD), and mean diffusional kurtosis (DKI-MK). In addition, the parameters of the DKI model were used to fit the biophysically-motivated White Matter Tract Integrity (WMTI) model (Fieremans et al., 2011), from which we derived a metric of axonal water fraction (DKI-AWF). In single-shell data, we fit the diffusion tensor imaging (DTI) model (Basser et al., 1994) using DIPY, and we derived fractional anisotropy (DTI-FA) and mean diffusivity (DTI-MD) metrics.

Tract profiles of tissue properties were derived by weighting the contribution of each streamline at the voxel that corresponds with each node, inversely weighted by that node’s distance from the median node in that position.

### 2.10 Reliability Estimates

The tractometry pipeline is composed of several different steps. To assess the effects of FWE on reliability at each one of these steps, both the split-half multi-shell and single-shell datasets were submitted to the same steps, with or without the preceding FWE procedure, and reliability was assessed across the two halves at each step, in each one of these cases.

#### 2.10.1 Fiber Orientation Distribution Functions (fODF)

We used the Constrained Spherical Deconvolution (CSD) model (Tournier et al., 2007), implemented in DIPY, to estimate fODFs. To assess reliability, each fODF was discretized to 362 uniformly-distributed positions on the sphere. Pearson’s correlation coefficient was obtained by correlating the discretized fODF for each voxel across split-halves. FODF reliability was assessed in terms of explained variance, quantified as the squared Pearson’s correlation coefficient, denoted as *r*^2^. Percent increase in fODF reliability in FWE, relative to the original data, was calculated as:

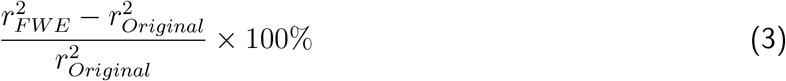

#### 2.10.2 Tract delineation

Tract density maps were created by counting the number of streamlines that pass through each voxel for each participant, tract, dataset, and tractography method. The Dice coefficient, weighted by the streamline visitation map (Cousineau et al., 2017) was computed across the tract density maps of each split-half.

#### 2.10.3 Tract profiles

The intraclass correlation coefficient (ICC) was calculated across split-halves for each tract, dMRI metric, dataset, and tractography method. We treat each split-half as an independent rater, and used the ICC(2,1) implementation in pingouin-0.5.4 (Vallat, 2018).

### 2.11 Inferences from tractometry

The tract profiles for each participant were used as the features to Fazekas scoresm. We used DKI-AWF, DKI-FA, DKI-MD, and DKI-MK for the full multi-shell dataset, and we use DTI-FA and DTI-MD for the full single-shell dataset. The tract profiles were z-score standardized and PCA transformed prior to model fitting (separately for the test and training data). Both models used to predict Fazekas scores were implemented in scikit-learn (version 1.4.2; Pedregosa et al., 2011).

#### 2.11.1 Fazekas Score Prediction

We implemented 5-fold logistic regression to predict Fazekas scores using participant tract profiles, stratified into folds to include Fazekas scores that are proportional to the sample as a whole. The logistic regression model was trained using the Scikit Learn LogisticRegressionCV (Pedregosa et al., 2011), which used L2 regularization and performed a 3-fold internal cross-validation to tune the regularization parameter. The logistic regression was configured for multiclass classification and adjusted imbalanced Fazekas scores frequencies by adjusting class weights inversely proportional to their frequencies in the dataset. The cross-validation process was repeated 20 times using RepeatedStratifiedKFold to minimize variability from unrepresentative splits due to the uneven Fazekas scores distribution.

## 3 Results

### 3.1 Free water elimination improves reliability of tractometry

Reliability was assessed across both split-half multi-shell and single-shell datasets to evaluate how FWE impacts reliability at each step in the pipeline.

#### 3.1.1 Fiber orientation distribution functions (fODFs)

FODF reliability was quantified as the explained variance (Pearson correlation squared) between the fODF in the two split-halves, discretized to 362 points on the sphere, calculated in each voxel. Because we hypothesized that regions of leukoaraiosis would be particularly susceptible to confounds because of CSF intrusion, we split our analysis to voxels in WMH and voxels in NAWM (Figure 1A shows one participant’s data). FWE fODF reliability was 14.07% (± 0.90% SEM; Wilcoxon signed rank test: z = 14.19, p ¡ 0.0001) higher in NAWM and 37.02% (± 2.13% SEM; Wilcoxon signed rank test: z = 13.15, p ¡ 0.0001) higher in WMH in multi-shell data (Figure 1B). Similarly, FWE fODF reliability was 1.57% (± 0.05% SEM; Wilcoxon signed rank test: z = 15.72, p ¡ 0.0001) higher in NAWM and 8.23% (± 0.26% SEM; Wilcoxon signed rank test: z = 15.44, p ¡ 0.0001) higher in WMH in the single-shell data (Figure 1C).

**Figure 1:**
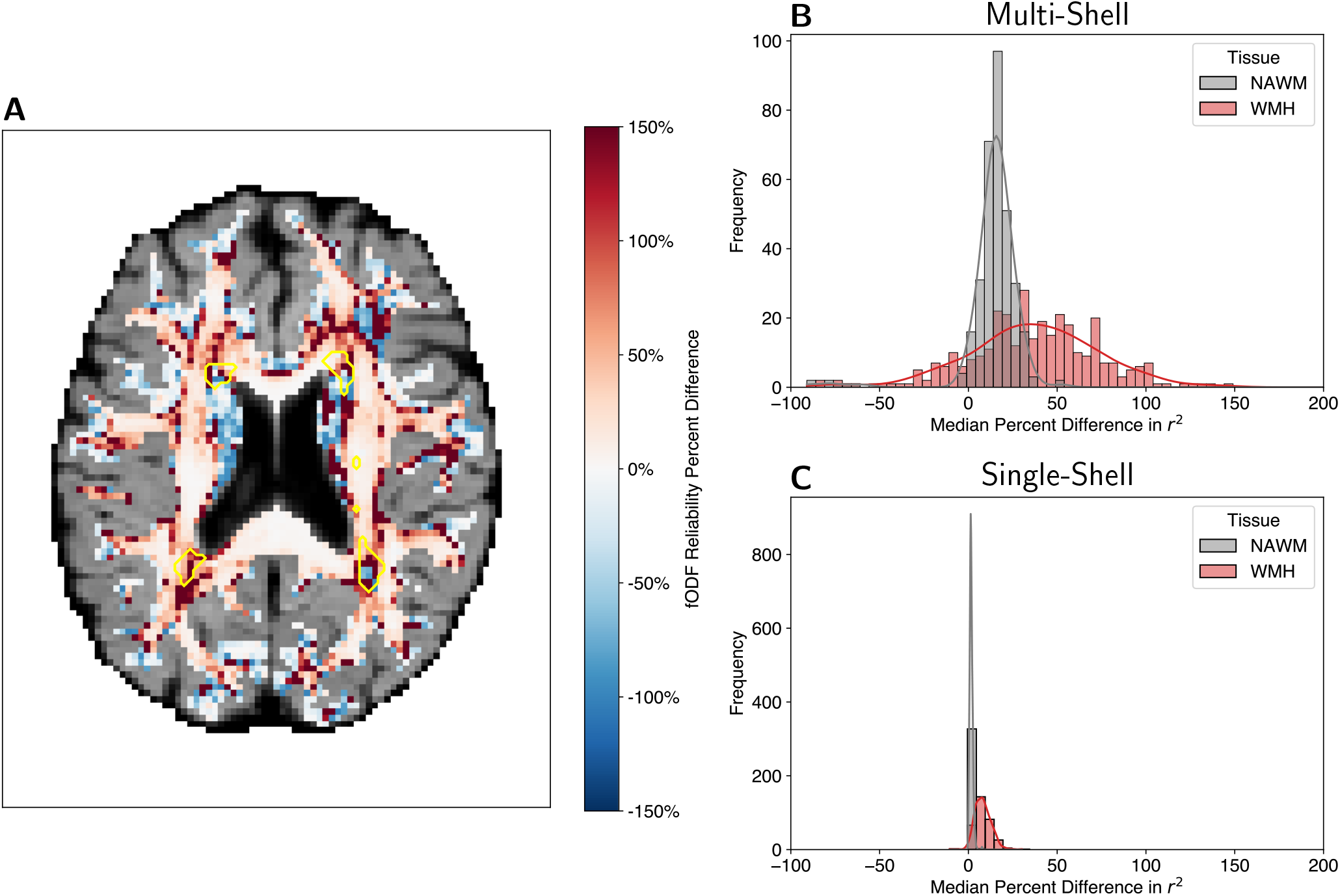
Fiber orientation distribution functions (fODFs) split-half reliability. **(A)** Percent difference (FWE - Original)/ Original in fODF reliability of an example participant. The yellow outlines represent regions of white matter hyperintensities. **(B)** Histogram of multi-shell split-half fODF reliability percent difference in normal appearing white matter (NAWM) and white matter hyperintensities (WMH). **(C)** Histogram of single-shell split-half fODF reliability percent difference in NAWM and WMH.

#### 3.1.2 Tract delineation

Tract delineation reliability was quantified as the difference in FWE from the original weighted Dice coefficient between tract density maps from each split-half of the data. On average, the weighted Dice coefficients across tracts were 0.50 (± 0.21 s.d.) for the multi-shell dataset with FWE processing, compared to an average 0.42 (± 0.25 s.d.) for the original tracts. For the single-shell dataset, FWE tracts had an average weighted Dice coefficient of 0.71 (± 0.20 s.d.), while the original tracts had an average of 0.67 (± 0.23 s.d.).

For the split-half multi-shell dataset, FWE tracts had larger weighted dice coefficients for all tracts except for the corticospinal tracts (CST L, CST R) and the cingulate section of the cingulum bundles (CGC L, CGC R) as compared to the original tracts (Figure 2). A smaller, yet still highly consistent increase in reliability with FWE was assessed in the split-half single-shell data.

**Figure 2:**
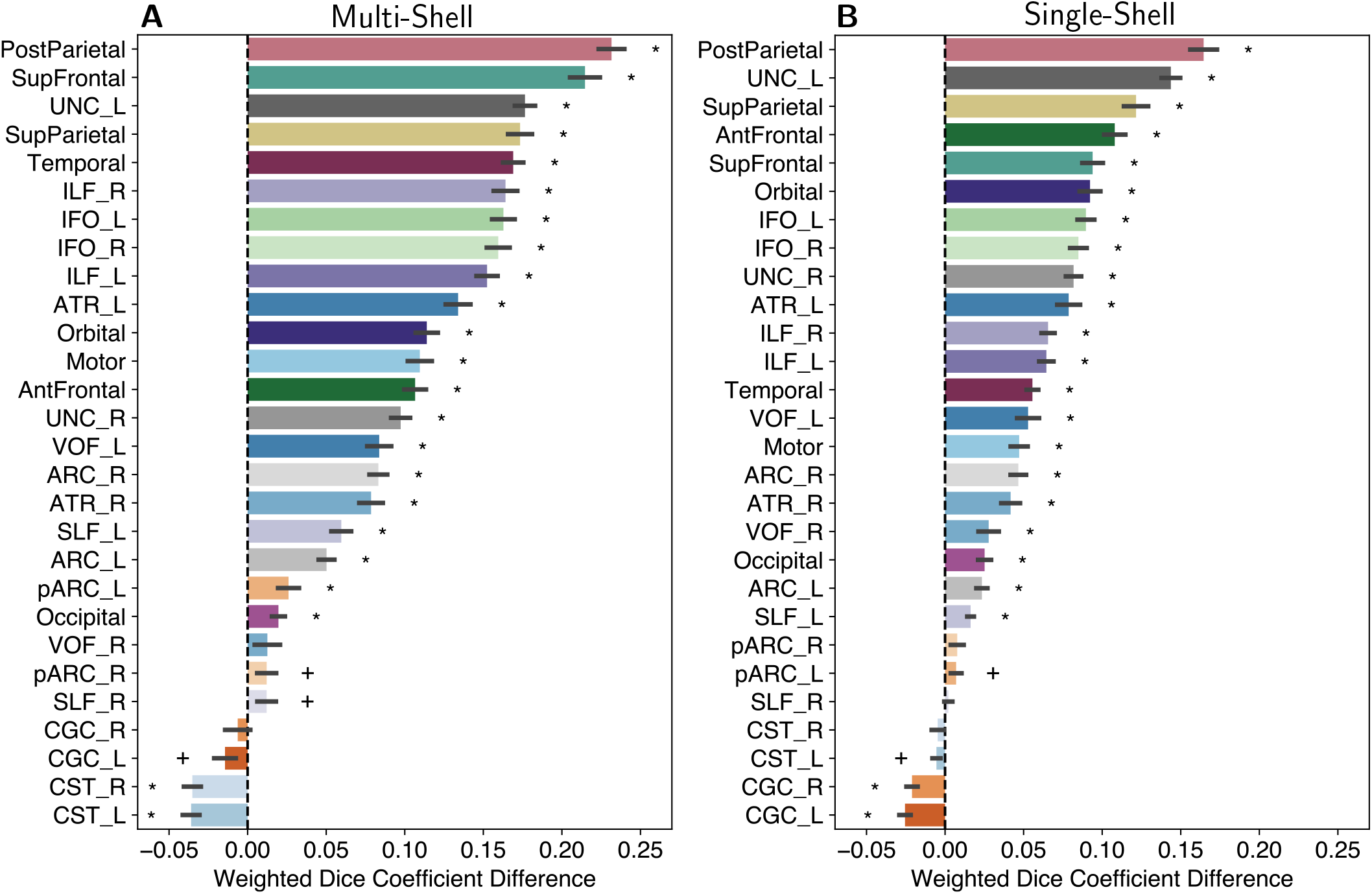
Tract weighted dice coefficient difference. Weighted dice coefficient differences shown for the split-half **(A)** multi-shell and **(B)** single-shell datasets. The difference was calculated as FEW - Original. Error bars represent ±1 SEM. Asterisks represent tracts with weighted Dice coefficient differences significantly different (Bonferonni-corrected) from 0. Crosses represent tracts with weighted Dice coefficient difference (without Bonferonni correction) from 0.

#### 3.1.3 Tract profiles

Tract profile reliability was quantified as the intraclass correlation coefficient (ICC) for each tract across split-halves. For the multi-shell dataset, profile ICC across tracts and dMRI metrics ranged from 0.42 to 0.90 for the FWE processed tracts and from 0.46 to 0.86 without FWE. For the single-shell dataset, profile ICC across tracts and metrics ranged from 0.84 to 0.96 with FWE and 0.78 to 0.94 without FWE. These ICC values are consistent with previously reported tract profile reliability estimates (Kruper et al., 2021).

The difference in tract profile ICC was used to compare tract profile reliability with or without FWE processing. For the multi-shell dataset, FWE tract profile ICCs for all DKI measures were greater or the same as the original tract profile ICC values (Figure 3 A, B, C, D). A similar, but substantially smaller effect was observed in the single-shell dataset (Figure 3 E, F).

**Figure 3:**
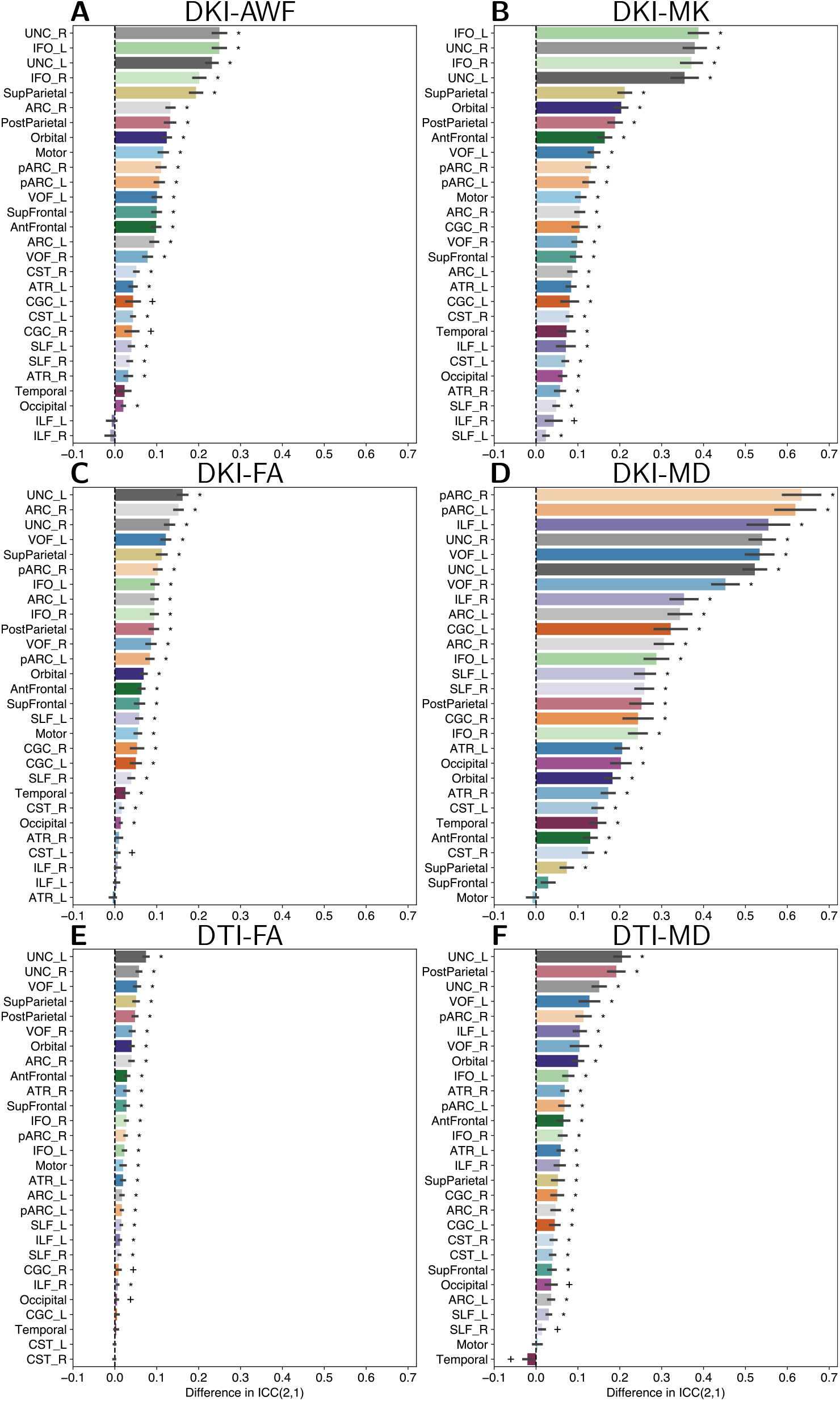
Differences in tract profile ICC(2,1). Tract profile ICC(2,1) differences are shown for split-half multi-shell **(A)** DKI-AWF, **(B)** DKI-MD, **(C)** DKI-FA, and **(D)** DKI-MD. Tract profile ICC(2,1) differences are shown split-half single-shell **(E)** DTI-FA and **(F)** DTI-MD. The difference was calculated as FWE - Original. Error bars represent ±1 SEM.

### 3.2 Free water elimination improves Fazekas score predictions

An indication that FWE results in improved inferences is apparent in a qualitative observation of tract profiles of the FA in the anterior thalamic radiation tract profiles, divided by the clinician WMH Fazekas scores. While tract profiles of different Fazekas score groups are hard to distinguish in the original data (Figure 4A, C), the profiles diverge in FWE data, both in multi-shell data (Figure 4B) and in single-shell data (Figure 4D). Figures showing qualitative observations in every tract and metric are provided in Supplemental Material.

**Figure 4:**
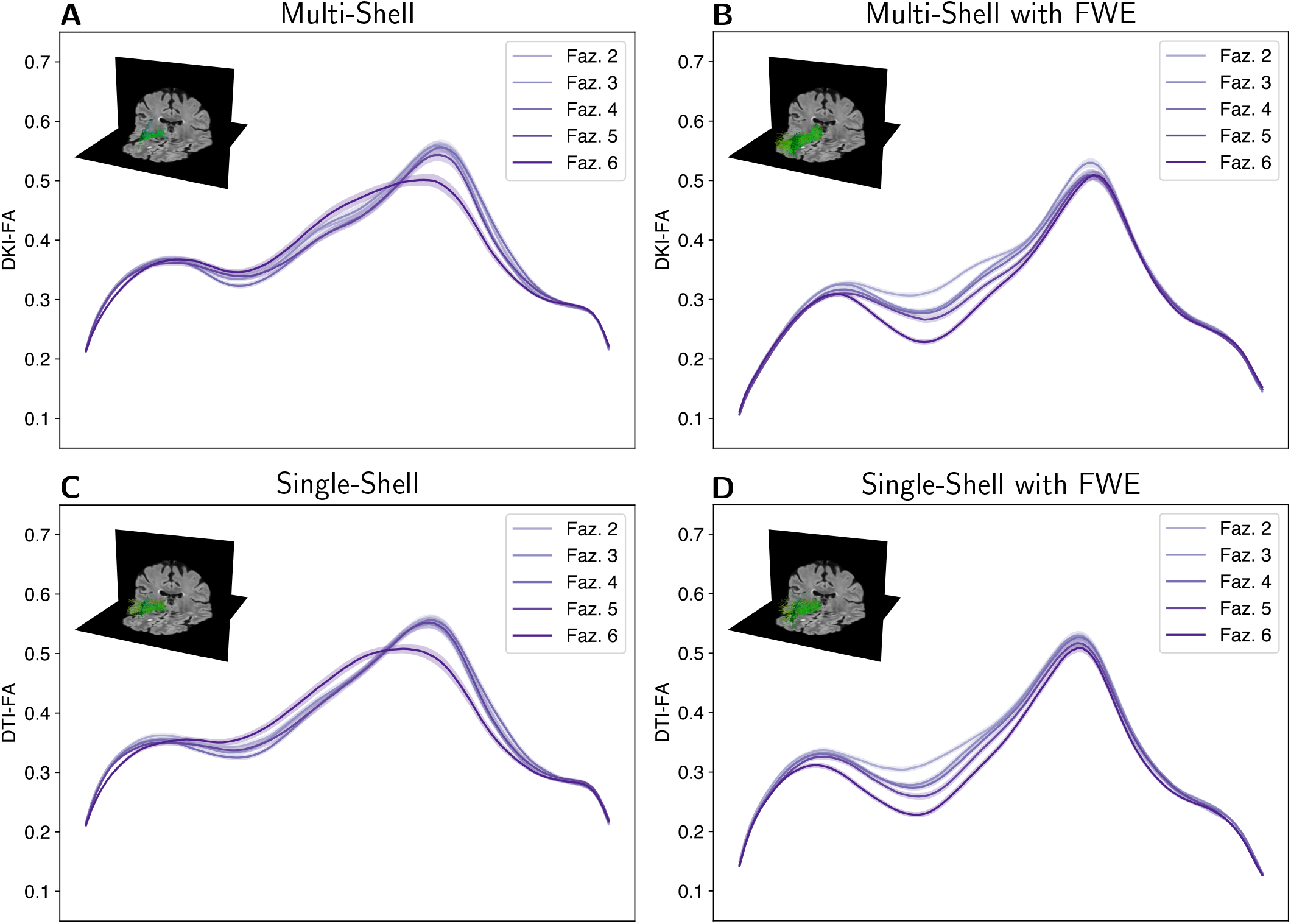
Right anterior thalamic radiation profiles by Fazekas scores. Tract profiles of multi-shell **(A)** DKI-FA and **(B)** DKI-FA with FWE are shown in the top row. The second row **(C)** single-shell DTI-FA and **(D)** single-shell DTI-FA with FWE. The x-axis represents positioning along the tract. The line colors correspond to Fazekas scores and the ribbon width represents ±1 SEM. Insets in each panel show the anterior thalamic radiations in the right hemisphere of the same participant (female, age = 75, Fazekas score = 5) with the FLAIR imaging as background

To quantify the qualitative observation from Figure 4, we used a machine learning approach to assess increase in the information that can be read out from the tract profiles regarding the extent and severity of leukoaraiosis. In this approach, a logistic regression model was fit to the tract profile data, classifying the individual-level Fazekas score. Predictions on held-out data were assessed in a 5-fold cross-validation procedure.

Fazekas score predictions improved for the multi-shell data by lowering the mean absolute error (MAE) from 0.96 to 0.93 points and increasing the variance explained from *R*^2^ = 0.14 to *R*^2^ = 0.20, an improvement of approximately 6%. A similar improved was shown for the FWE single-shell data, in which MAE decreased from 0.99 to 0.95 points and variance explained increased from *R*^2^ = 0.10 to *R*^2^ = 0.21, an improvement of approximately 11%

The classification results were compared using the area under the curve (AUC) from receiver operating characteristic (ROC) curves. We performed a series of ROC analyses to evaluate the model’s ability to discriminate between progressively higher Fazekas scores (Figure 5). The analysis began with a binary classification between the lowest Fazekas score and all higher scores. Specifically, we first examined the classification of Fazekas scores of 2 versus scores of 3, 4, 5, and 6. We then incrementally incorporated additional lower Fazekas scores in to the lower bin (e.g., {2, 3} vs. {4, 5, 6}). This stepwise approach continued until we compared all lower Fazekas scores {2, 3, 4, 5} against the highest score of 6.

**Figure 5:**
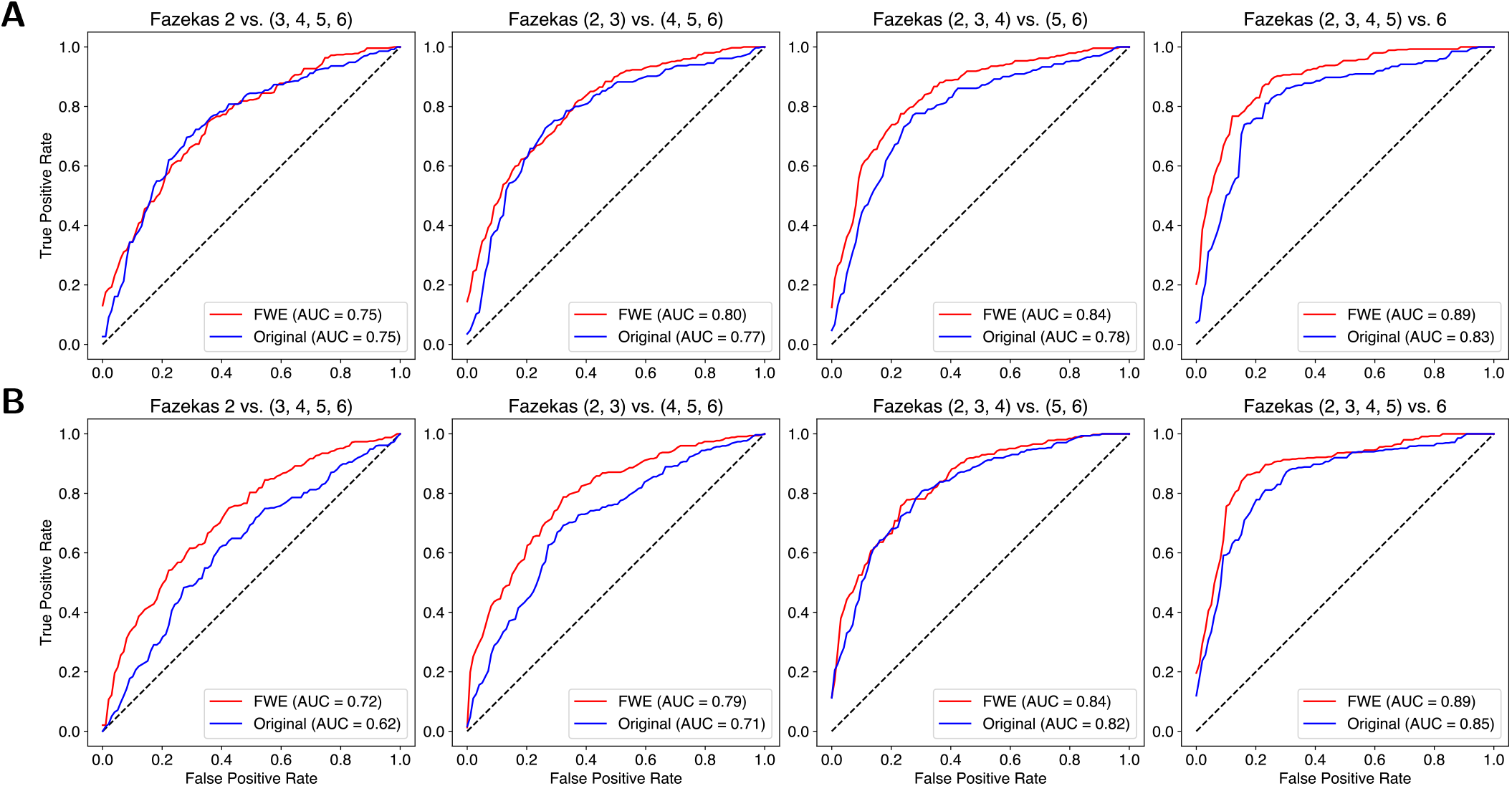
Fazekas score receiver operating characteristic (ROC) curves from whole-brain tract profiles. **(A)** The first row shows the AUC and ROC curves for the multi-shell data. **(B)** The second row shows the AUC and ROC curves for the single-shell data.

FWE tract profiles had similar or improved AUC discrimination performance in the ROC analysis for both the multi-shell (Figure 5A) and single-shell (Figure 5B) data. Differences were assessed for statistical significance using Delong’s test (DeLong et al., 1988; Sun & Xu, 2014). For the multi-shell data, FWE tract profiles significantly improved discrimination the highest Fazekas scores from all other scores: {2, 3, 4, 5} vs. 6 scores, *p* = 0.0076. The single-shell FWE tract profiles performed significantly better when discriminating 2 vs. {3, 4, 5, 6} scores, *p* = 0.0034, {2, 3} vs. {4, 5, 6} scores, *p* = 0.0002, and {2, 3, 4, 5} vs. 6 scores, *p* = 0.0155.

## 4 Control Analyses

One potential interpretation of the results is that FWE increases the number of streamlines that are produced in tractography, because it effectively lowers the FA threshold at which tractography terminates. A control analysis where the same analysis was conducted with 50% lowered FA threshold determined that this did not improve reliability (see Supplemental Material).

Another important question is whether these results generalize to populations in other age groups. An analysis of the Human Connectome Project Young Adult test-retest dataset (Glasser et al., 2016), showed that the benefits of FWE are substantially smaller in this young adult dataset, suggesting that FWE is particularly well-suited for analysis of older adults (see Supplemental Material).

## 5 Discussion

Tractometry is a powerful tool for studying human brain white matter connections, with applications across a range of different research questions, and potential applications in clinical settings. Here, we demonstrate that a straight-forward computational FWE procedure improves reliability in every step of the tractometry process in the brains of aging individuals. The most notable improvements were in fODF reliability throughout the white matter, with particularly large effects in WMH regions. These regions, and particularly periventricular WMH regions, are known to have high MD, low FA, and a lower magnetization transfer ratio (Bastin et al., 2009; Maniega et al., 2015), reflecting pathological processes of tissue degeneration, axonal loss, and demyelination. Nevertheless, even while tissue biophysics is significantly altered, there is a residual directional signal in these regions that was highly reproducible across splits of the data once free water effects were eliminated. The increased reliability in fODFs translates into more reliably delineated tracts. Particularly strong effects are observed in callosal bundles and the left uncinate. No effects, or even small detrimental effects are observed in the corticospinal tract and cingulum cingulate bundles, which may be positioned or organized such that they are less susceptible to free water contamination.

While improving reliability is beneficial on its own, it could also come at the expense of increased bias, as with many other denoising procedures (Kay, 2022). Therefore, it is important to validate that the observed reliability improvements also translate to improved accuracy. In the absence of independent measurements of the same brain tissue (e.g., histology), an effective way to assess improved accuracy is by demonstrating that the method improves inferences about individual differences in phenotypic variables, such as clinical status. Indeed, FWE tractometry improves the prediction of clinically-obtained Fazekas scores based solely on FLAIR images. Previous research demonstrated that WMH are associated with altered white matter tissue properties (Chang et al., 2023; Min et al., 2021; Wardlaw et al., 2015). Our work extends previous findings by linking these changes to specific anatomical tracts, demonstrating the potential to provide individualized assessments of WMH burden and its functional implications. Though the predictive performance is still limited (*R*^2^ = 0.21 in the best case), we anticipate improvements with the advancement of more sophisticated machine learning models, such as deep learning approaches (Kruper et al., 2024). Regardless of the limited accuracy demonstrated here, the improvement with FWE demonstrate the increased information captured through this method.

The FWDTI model has come under recent renewed scrutiny, particularly in the single-shell case, where it has been demonstrated that the single-shell FWDTI model biased FWE-corrected metrics, such as FA and MD, due to non-Gaussian diffusion effects (Correia et al., 2024). We acknowledge these issues, but emphasize that these issues are unrelated to the benefits we demonstrate here. In fact, our study avoids this pitfall, because in our approach, tractography is derived from the FWE-corrected signal, while tractometry metrics are derived from the original, non-FWE-corrected diffusion data. By doing so, we mitigate the risk of introducing bias into the diffusion metrics and preserve the benefits of FWE tractography, ensuring a more reliable and accurate analysis of white matter tract and tissue properties.

### 5.1 Conclusion

Our study demonstrates the benefits of applying FWE to improve the reliability and accuracy of tractometry in aging brains, particularly in regions affected by WMH. By mitigating the effects of free water contamination, we increase the reliability of all stages of tractometry: fODF estimation, tract delineation, and tract profiling. These improvements lead to greater accuracy in making diffusion tractometry inferences about individual differences in clinical and phenotypic measure of Fazekas scores. Our approach avoids the biases identified in FWDTI modeling, providing a processing pipeline that supports the study of white matter changes and its relationship to aging and disease.

## Supporting information

Supplemental Material to "Free-water elimination tractometry for aging brains"

## Data and Code Availability

Code to perform free water elimination is available on GitHub: https://github.com/nrdg/fwe. Data can be accessed through https://www.actagingresearch.org/

## Author Contributions

**Kelly Chang**: Conceptualization, Methodology, Software, Validation, Formal analysis, Investigation, Writing - Original Draft, Writing - Review & Editing, Visualization

**Luke Burke**: Investigation, Data Curation

**Nina LaPiana**: Investigation, Data Curation

**Bradley Howlett**: Investigation, Data Curation

**David Hunt**: Investigation, Data Curation

**Margaret Dezelar**: Project administration, Data Curation

**Jalal B. Andre**: Investigation, Data Curation, Writing - Review & Editing

**Patti Curl**: Investigation, Data Curation, Writing - Review & Editing

**James Ralston**: Investigation, Writing - Review & Editing

**Ariel Rokem**: Conceptualization, Methodology, Resources, Writing - Original Draft, Writing - Review & Editing, Supervision, Funding acquisition

**Christine MacDonald**: Investigation, Resources, Data Curation, Supervision, Project administration, Funding acquisition

## Acknowledgments

We thank the participants of the Adult Changes in Thought (ACT) study for the data they have provided and the many ACT investigators and staff who steward that data. You can learn more about ACT at: https://actagingstudy.org/.

## References

Andersson, J. L., Skare, S., & Ashburner, J. (2003). How to correct susceptibility distortions in spin-echo echo-planar images: Application to diffusion tensor imaging. NeuroImage, 20 (2), 870–888. 10.1016/S1053-8119(03)00336-7

Andersson, J. L., & Sotiropoulos, S. N. (2016). An integrated approach to correction for off-resonance effects and subject movement in diffusion MR imaging. NeuroImage, 125, 1063–1078. 10.1016/j.neuroimage.2015.10.019

Avants, B., Epstein, C., Grossman, M., & Gee, J. (2008). Symmetric diffeomorphic image registration with cross-correlation: Evaluating automated labeling of elderly and neurodegenerative brain. Medical Image Analysis, 12 (1), 26–41. 10.1016/j.media.2007.06.004

Basser, P. J., Mattiello, J., & LeBihan, D. (1994). MR diffusion tensor spectroscopy and imaging. Biophys. J., 66 (1), 259–267. 10.1016/S0006-3495(94)80775-1

Bastin, M. E., Clayden, J. D., Pattie, A., Gerrish, I. F., Wardlaw, J. M., & Deary, I. J. (2009). Diffusion tensor and magnetization transfer MRI measurements of periventricular white matter hyperintensities in old age. Neurobiol. Aging, 30 (1), 125–136. 10.1016/j.neurobiolaging.2007.05.013

Billot, B., Greve, D. N., Puonti, O., Thielscher, A., Van Leemput, K., Fischl, B., Dalca, A. V., & Iglesias, J. E. (2023). SynthSeg: Segmentation of brain MRI scans of any contrast and resolution without retraining. Med. Image Anal., 86, 102789. 10.1016/j.media.2023.102789

Billot, B., Magdamo, C., Cheng, Y., Arnold, S. E., Das, S., & Iglesias, J. E. (2023). Robust machine learning segmentation for large-scale analysis of heterogeneous clinical brain MRI datasets. Proceedings of the National Academy of Sciences, 120. 10.1073/pnas.2216399120

Chang, K., Burke, L., LaPiana, N., Howlett, B., Hunt, D., Dezelar, M., Andre, J. B., Ralston, J., Rokem, A., & Donald, C. M. (2023). Advanced diffusion MRI modeling sheds light on FLAIR white matter hyperintensities in an aging cohort. Computational Diffusion MRI, 192–203. 10.1007/978-3-031-47292-317

Cieslak, M., Cook, P. A., He, X., Yeh, F.-C., Dhollander, T., Adebimpe, A., Aguirre, G. K., Bassett, D. S., Betzel, R. F., Bourque, J., Cabral, L. M., Davatzikos, C., Detre, J. A., Earl, E., Elliott, M. A., Fadnavis, S., Fair, D. A., Foran, W., Fotiadis, P., … Satterthwaite, T. D. (2021). QSIPrep: An integrative platform for preprocessing and reconstructing diffusion MRI data. Nature Methods, 18 (7), 775–778. 10.1038/s41592-021-01185-5

Correia, M. M., Henriques, R. N., Golub, M., Winzeck, S., & Nunes, R. G. (2024). The trouble with free-water elimination using single-shell diffusion MRI data: A case study in ageing. Imaging Neuroscience, 2, 1–17. 10.1162/imaga00252

Cousineau, M., Jodoin, P.-M., Garyfallidis, E., Côté, M.-A., Morency, F. C., Rozanski, V., Grand’Maison, M., Bedell, B. J., & Descoteaux, M. (2017). A test-retest study on Parkinson’s PPMI dataset yields statistically significant white matter fascicles. NeuroImage: Clinical, 16, 222–233. 10.1016/j.nicl.2017.07.020

Cox, S. R., Ritchie, S. J., Tucker-Drob, E. M., Liewald, D. C., Hagenaars, S. P., Davies, G., Wardlaw, J. M., Gale, C. R., Bastin, M. E., & Deary, I. J. (2016). Ageing and brain white matter structure in 3,513 UK biobank participants. Nat. Commun., 7, 13629. 10.1038/ncomms13629

DeLong, E. R., DeLong, D. M., & Clarke-Pearson, D. L. (1988). Comparing the areas under two or more correlated receiver operating characteristic curves: A nonparametric approach. Biometrics, 44 (3), 837–845. 10.2307/2531595

Dougherty, R. F., Ben-Shachar, M., Deutsch, G. K., Hernandez, A., Fox, G. R., & Wandell, B. A. (2007). Temporal-callosal pathway diffusivity predicts phonological skills in children. Proc. Natl. Acad. Sci. U. S. A., 104 (20), 8556–8561.

Fazekas, F., Chawluk, J. B., Alavi, A., Hurtig, H. I., & Zimmerman, R. A. (1987). MR signal abnormalities at 1.5 T in Alzheimer’s dementia and normal aging. AJR Am. J. Roentgenol., 149 (2), 351–356. 10.2214/ajr.149.2.351

Fieremans, E., Jensen, J. H., & Helpern, J. A. (2011). White matter characterization with diffusional kurtosis imaging. NeuroImage, 58 (1), 177–188. 10.1016/j.neuroimage.2011.06.006

Fonov, V., Evans, A., McKinstry, R., Almli, C., & Collins, D. (2009). Unbiased nonlinear average age-appropriate brain templates from birth to adulthood. NeuroImage, 47, S102. 10.1016/S1053-8119(09)70884-5

Garyfallidis, E., Brett, M., Amirbekian, B., Rokem, A., Van Der Walt, S., Descoteaux, M., & Nimmo-Smith, I. (2014). Dipy, a library for the analysis of diffusion MRI data. Frontiers in Neuroinformatics, 8. 10.3389/fninf.2014.00008

Glasser, M. F., Smith, S. M., Marcus, D. S., Andersson, J. L. R., Auerbach, E. J., Behrens, T. E. J., Coalson, T. S., Harms, M. P., Jenkinson, M., Moeller, S., Robinson, E. C., Sotiropoulos, S. N.,Xu, J., Yacoub, E., Ugurbil, K., & Van Essen, D. C. (2016). The Human Connectome Project’s neuroimaging approach. Nat. Neurosci., 19 (9), 1175–1187. 10.1038/nn.4361

Golub, M., Neto Henriques, R., & Gouveia Nunes, R. (2021). Free-water DTI estimates from single b-value data might seem plausible but must be interpreted with care. Magn. Reson. Med., 85 (5), 2537–2551. 10.1002/mrm.28599

Henriques, R. N., Correia, M. M., Marrale, M., Huber, E., Kruper, J., Koudoro, S., Yeatman, J. D., Garyfallidis, E., & Rokem, A. (2021). Diffusional kurtosis imaging in the diffusion imaging in python project. Frontiers in Human Neuroscience, 15. 10.3389/fnhum.2021.675433

Henriques, R. N., Rokem, A., Garyfallidis, E., St-Jean, S., Peterson, E. T., & Correia, M. M. (2017). [Re] Optimization of a free water elimination two-compartment model for diffusion tensor imaging. ReScience, 3. 10.1101/108795

Hoopes, A., Mora, J. S., Dalca, A. V., Fischl, B., & Hoffmann, M. (2022). SynthStrip: Skull-stripping for any brain image. NeuroImage, 260, 119474. 10.1016/j.neuroimage.2022.119474

Hoy, A. R., Kecskemeti, S. R., & Alexander, A. L. (2015). Free water elimination diffusion tractography: A comparison with conventional and fluid-attenuated inversion recovery, diffusion tensor imaging acquisitions. J. Magn. Reson. Imaging, 42 (6), 1572–1581. 10.1002/jmri.24925

Hoy, A. R., Koay, C. G., Kecskemeti, S. R., & Alexander, A. L. (2014). Optimization of a free water elimination two-compartment model for diffusion tensor imaging. NeuroImage, 103, 323–333. 10.1016/j.neuroimage.2014.09.053

Irfanoglu, M. O., Modi, P., Nayak, A., Hutchinson, E. B., Sarlls, J., & Pierpaoli, C. (2015). DR-BUDDI (Diffeomorphic Registration for Blip-Up blip-Down Diffusion Imaging) method for correcting echo planar imaging distortions. NeuroImage, 106, 284–299. 10.1016/j.neuroimage.2014.11.042

Jensen, J. H., Helpern, J. A., Ramani, A., Lu, H., & Kaczynski, K. (2005). Diffusional kurtosis imaging: The quantification of non-gaussian water diffusion by means of magnetic resonance imaging. Magnetic Resonance in Medicine, 53 (6), 1432–1440. 10.1002/mrm.20508

Jeurissen, B., Descoteaux, M., Mori, S., & Leemans, A. (2019). Diffusion MRI fiber tractography of the brain. NMR Biomed., 32 (4), e3785. 10.1002/nbm.3785

Kay, K. (2022). The risk of bias in denoising methods: Examples from neuroimaging. PLoS One, 17 (7), e0270895. 10.1371/journal.pone.0270895

Kellner, E., Dhital, B., Kiselev, V. G., & Reisert, M. (2016). Gibbs-ringing artifact removal based on local subvoxel-shifts. Magn. Reson. Med., 76, 1574–1581. 10.1002/mrm.26054

Kruper, J., Richie-Halford, A., Benson, N. C., Caffarra, S., Owen, J., Wu, Y., Egan, C., Lee, A. Y., Lee, C.S., Yeatman, J. D., Rokem, A., & UK Biobank Eye and Vision Consortium. (2024). Convo-lutional neural network-based classification of glaucoma using optic radiation tissue properties. Commun. Med., 4 (1), 72. 10.1038/s43856-024-00496-w

Kruper, J., Yeatman, J. D., Richie-Halford, A., Bloom, D., Grotheer, M., Caffarra, S., Kiar, G., Karipidis, I.I., Roy, E., Chandio, B. Q., Garyfallidis, E., & Rokem, A. (2021). Evaluating the reliability of human brain white matter tractometry. Aperture Neuro, 1 (1). 10.52294/e6198273-b8e3-4b63-babb-6e6b0da10669

Kukull, W. A., Higdon, R., Bowen, J. D., McCormick, W. C., Teri, L., Schellenberg, G. D., van Belle, G., Jolley, L., & Larson, E. B. (2002). Dementia and Alzheimer Disease Incidence: A Prospective Cohort Study. Archives of Neurology, 59, 1737–1746. 10.1001/archneur.59.11.1737

Maniega, S. M., Valdés Hernández, M. C., Clayden, J. D., Royle, N. A., Murray, C., Morris, Z., Aribisala, B.S., Gow, A. J., Starr, J. M., Bastin, M. E., Deary, I. J., & Wardlaw, J. M. (2015). White matter hyperintensities and normal-appearing white matter integrity in the aging brain. Neurobiol. Aging, 36 (2), 909–918. 10.1016/j.neurobiolaging.2014.07.048

Min, Z.-G., Shan, H.-R., Xu, L., Yuan, D.-H., Sheng, X.-X., Xie, W.-C., Zhang, M., Niu, C., Shakir, T. M.,& Cao, Z.-H. (2021). Diffusion tensor imaging revealed different pathological processes of white matter hyperintensities. BMC Neurol., 21 (1), 128. 10.1186/s12883-021-02140-9

Mojiri Forooshani, P., Biparva, M., Ntiri, E. E., Ramirez, J., Boone, L., Holmes, M. F., Adamo, S., Gao, F., Ozzoude, M., Scott, C. J. M., Dowlatshahi, D., Lawrence-Dewar, J. M., Kwan, D., Lang, A.E., Marcotte, K., Leonard, C., Rochon, E., Heyn, C., Bartha, R., … Goubran, M. (2022). Deep bayesian networks for uncertainty estimation and adversarial resistance of white matter hyperintensity segmentation. Human Brain Mapping, 43 (7), 2089–2108. 10.1002/hbm.25784

Ould Ismail, A. A., Parker, D., Hernandez-Fernandez, M., Brem, S., Alexander, S., Pasternak, O., Caruyer, E., & Verma, R. (2019). Characterizing peritumoral tissue using DTI-based free water elimination. Brainlesion: Glioma, Multiple Sclerosis, Stroke and Traumatic Brain Injuries, 123–131. 10.1007/978-3-030-11723-812

Pasternak, O., Sochen, N., Gur, Y., Intrator, N., & Assaf, Y. (2009). Free water elimination and mapping from diffusion MRI. Magn. Reson. Med., 62 (3), 717–730. 10.1002/mrm.22055

Pedregosa, F., Varoquaux, G., Gramfort, A., Michel, V., Thirion, B., Grisel, O., Blondel, M., Prettenhofer, P., Weiss, R., Dubourg, V., Vanderplas, J., Passos, A., Cournapeau, D., Brucher, M., Perrot, M., & Duchesnay, E. (2011). Scikit-learn: Machine learning in Python. Journal of Machine Learning Research, 12, 2825–2830.

Pierpaoli, C., & Jones, D. K. (2004). Removing CSF contamination in brain DT-MRIs by using a two-compartment tensor model. Interntational Society for Magnetic Resonance in Medicine, 1215.

Richie-Halford, A., Cieslak, M., Ai, L., Caffarra, S., Covitz, S., Franco, A. R., Karipidis, I. I., Kruper, J., Milham, M., Avelar-Pereira, B., Roy, E., Sydnor, V. J., Yeatman, J. D., Fibr Community Science Consortium, Satterthwaite, T. D., & Rokem, A. (2022). An analysis-ready and quality controlled resource for pediatric brain white-matter research. Sci Data, 9 (1), 616. 10.1038/s41597-022-01695-7

Sun, X., & Xu, W. (2014). Fast implementation of DeLong’s algorithm for comparing the areas under correlated receiver operating characteristic curves. IEEE Signal Processing Letters, 21 (11), 1389– 1393. 10.1109/LSP.2014.2337313

Takemura, H., Kruper, J. A., Miyata, T., & Rokem, A. (2024). Tractometry of human visual white matter pathways in health and disease. Magnetic Resonance in Medical Sciences, 23 (3), 316– 340. 10.2463/mrms.rev.2024-0007

Tournier, J.-D., Calamante, F., & Connelly, A. (2007). Robust determination of the fibre orientation distribution in diffusion MRI: Non-negativity constrained super-resolved spherical deconvolution. NeuroImage, 35 (4), 1459–1472. 10.1016/j.neuroimage.2007.02.016

Tustison, N. J., Avants, B. B., Cook, P. A., Zheng, Y., Egan, A., Yushkevich, P. A., & Gee, J. C. (2010). N4ITK: Improved N3 bias correction. IEEE Transactions on Medical Imaging, 29 (6), 1310–1320. 10.1109/TMI.2010.2046908

Vallat, R. (2018). Pingouin: Statistics in python. Journal of Open Source Software, 3 (31), 1026. 10.21105/joss.01026

van der Walt, S., Schönberger, J. L., Nunez-Iglesias, J., Boulogne, F., Warner, J. D., Yager, N., Gouillart, E., Yu, T., & the scikit-image contributors. (2014). scikit-image: Image processing in Python. PeerJ, 2, e453. 10.7717/peerj.453

Veraart, J., Novikov, D. S., Christiaens, D., Ades-aron, B., Sijbers, J., & Fieremans, E. (2016). Denoising of diffusion MRI using random matrix theory. NeuroImage, 142, 394–406. 10.1016/j.neuroimage.2016.08.016

Wardlaw, J. M., Valdés Hernández, M. C., & Muñoz-Maniega, S. (2015). What are white matter hyperintensities made of? Relevance to vascular cognitive impairment. J. Am. Heart Assoc., 4 (6), 001140. 10.1161/JAHA.114.001140

Yeatman, J. D., Dougherty, R. F., Myall, N. J., Wandell, B. A., & Feldman, H. M. (2012). Tract profiles of white matter properties: Automating fiber-tract quantification. PLoS One, 7 (11), e49790. 10.1371/journal.pone.0049790

Yeh, F.-C., Zaydan, I. M., Suski, V. R., Lacomis, D., Richardson, R. M., Maroon, J. C., & Barrios-Martinez, J. (2019). Differential tractography as a track-based biomarker for neuronal injury. NeuroImage, 202, 116131. 10.1016/j.neuroimage.2019.116131

