## Supplemental Material to "Free-water elimination tractometry for aging brains" for "Free water elimination tractometry for aging brains"

November 10, 2024

### 1 Supplemental Analyses

#### 1.1 Lowered Threshold Original Diffusion

We performed the same analysis on the original dataset with a lowered stopping FA threshold (50% decrease) to evaluate the effect of FWE tractography. Overall, we found that lowering the stopping FA criterion increased the number of streamlines produced, but the results were similar to the original datasets when compared to FWE tractography.

##### 1.1.1 Tract delineation reliability

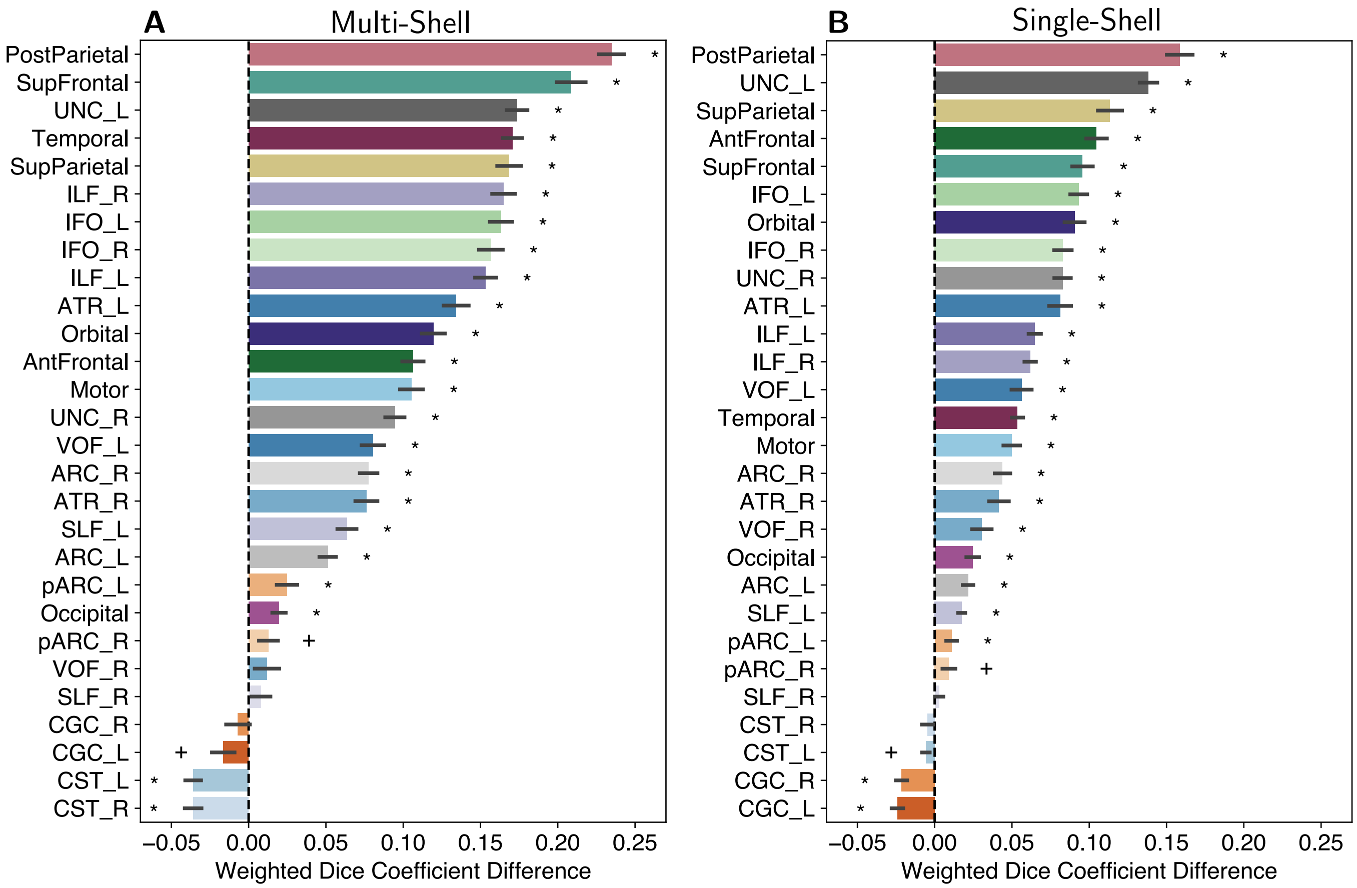


Figure 1: Tract weighted dice coefficient difference. Weighted dice coefficient differences shown for the split-half **(A)** multi-shell and **(B)** single-shell datasets. The difference was calculated as FWE - Original with lowered FA threshold. Error bars represent $\pm 1$ SEM. Asterisks represent tracts with weighted Dice coefficient differences significantly different (Bonferonni-corrected) from 0. Crosses represent tracts with weighted Dice coefficient difference (without Bonferonni correction) from 0.

##### 1.1.2 Tract profile reliability

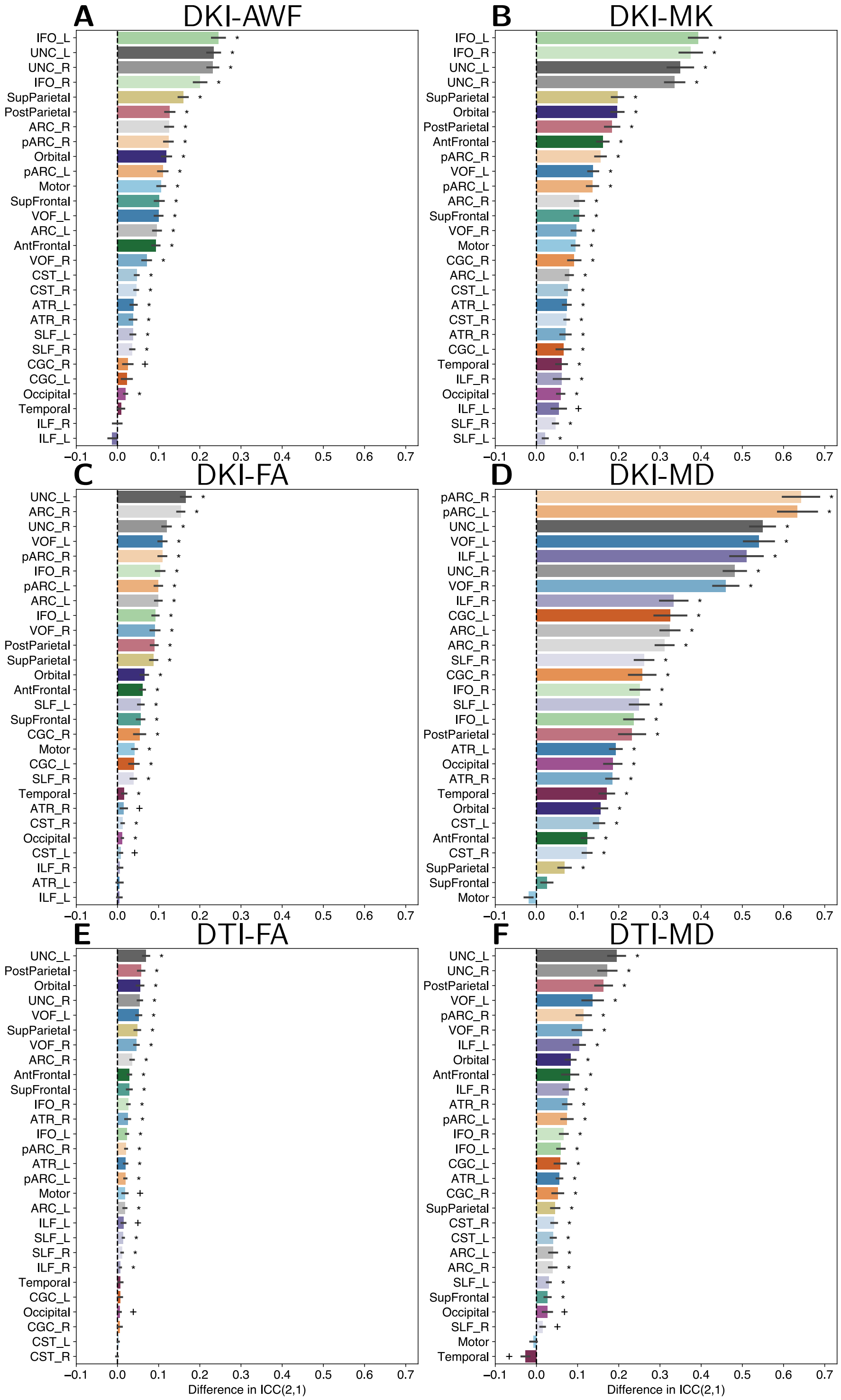


Figure 2: Differences in tract profile ICC(2,1). Tract profile ICC(2,1) differences are shown for split-half multi-shell **(A)** DKI-AWF, **(B)** DKI-MD, **(C)** DKI-FA, and **(D)** DKI-MD. Tract profile ICC(2,1) differences are shown split-half single-shell **(E)** DTI-FA and **(F)** DTI-MD. The difference was calculated as FWE - Original with lowered FA threshold. Error bars represent ±1 SEM.

##### 1.1.3 Fazekas score predictions

Fazekas score predictions improved for the multi-shell data by lowering the mean absolute error (MAE) from 0.98 to 0.85 points and increasing the variance explained from $R^2 = 0.16$ to $R^2 = 0.26$, an improvement of approximately $10\%$. A similar improved was shown for the FWE single-shell data, in which MAE decreased from 0.99 to 0.96 points and variance explained increased from $R^2 = 0.09$ to $R^2 = 0.14$, an improvement of approximately $5\%$

Differences in Fazekas score predictions were assessed for statistical significance using Delong's test (**DeLong1988-zc**; Sun & Xu, 2014). For the multi-shell data, FWE tract profiles significantly improved discrimination the lowest Fazekas scores from all other scores: 2 vs. {3, 4, 5, 6} scores, $p = 0.0004$, and {2, 3} vs. {4, 5, 6} scores, $p = 0.0361$. The single-shell FWE tract profiles performed significantly better when discriminating {2, 3} vs. {4, 5, 6} scores, $p = 0.0406$.

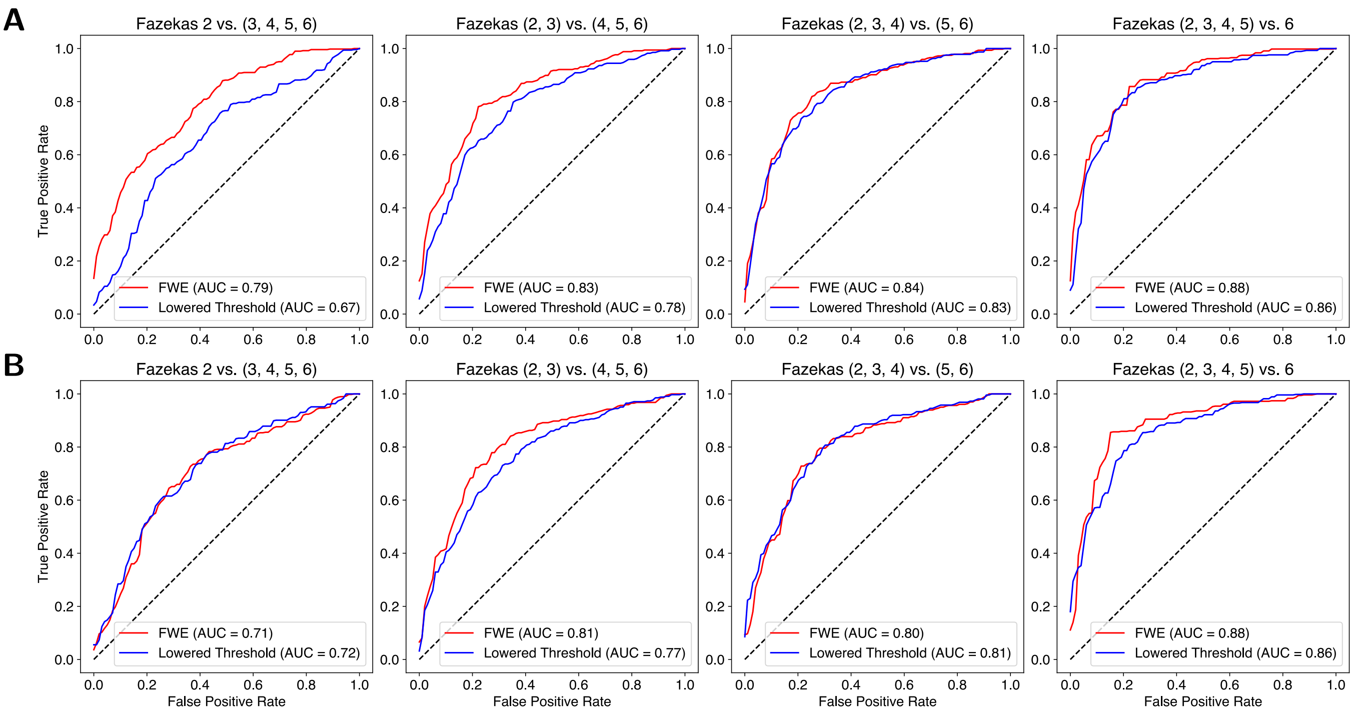


Figure 3: Fazekas score receiver operating characteristic (ROC) curves from whole-brain tract profiles. **(A)** The first row shows the AUC and ROC curves for the multi-shell data. **(B)** The second row shows the AUC and ROC curves for the single-shell data. The red lines represent Fazekas score prediction from FWE tract profiles and blue lines represent the Fazekas score prediction from the original tract profiles with lowered FA threshold.

#### 1.2 HCP Test-Retest dataset

We performed the same analysis with the HCP Test-Retest dataset to evaluate the importance of age for FWE tractography. Overall, we found that the benefits of FWE on tractography were minimal.

##### 1.2.1 fODF reliability

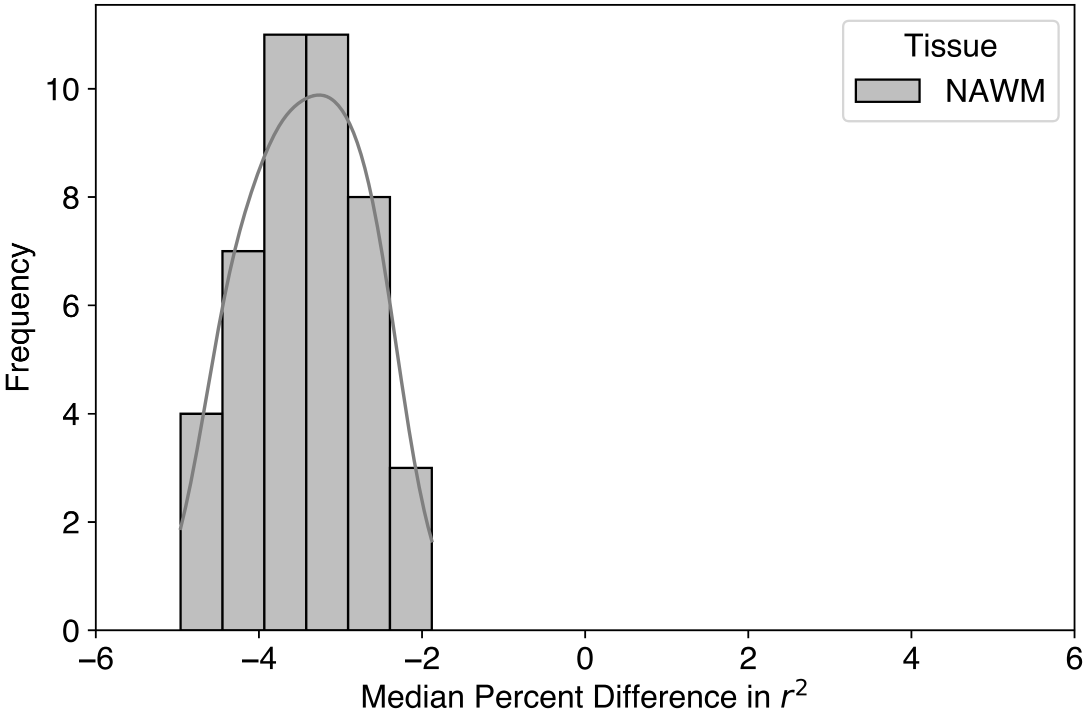


Figure 4: fODF split-half reliability for the HCP Test-Retest dataset. Histogram of split-half fODF reliability percent difference in normal appearing white matter (NAWM).

##### 1.2.2 Tract delineation reliability

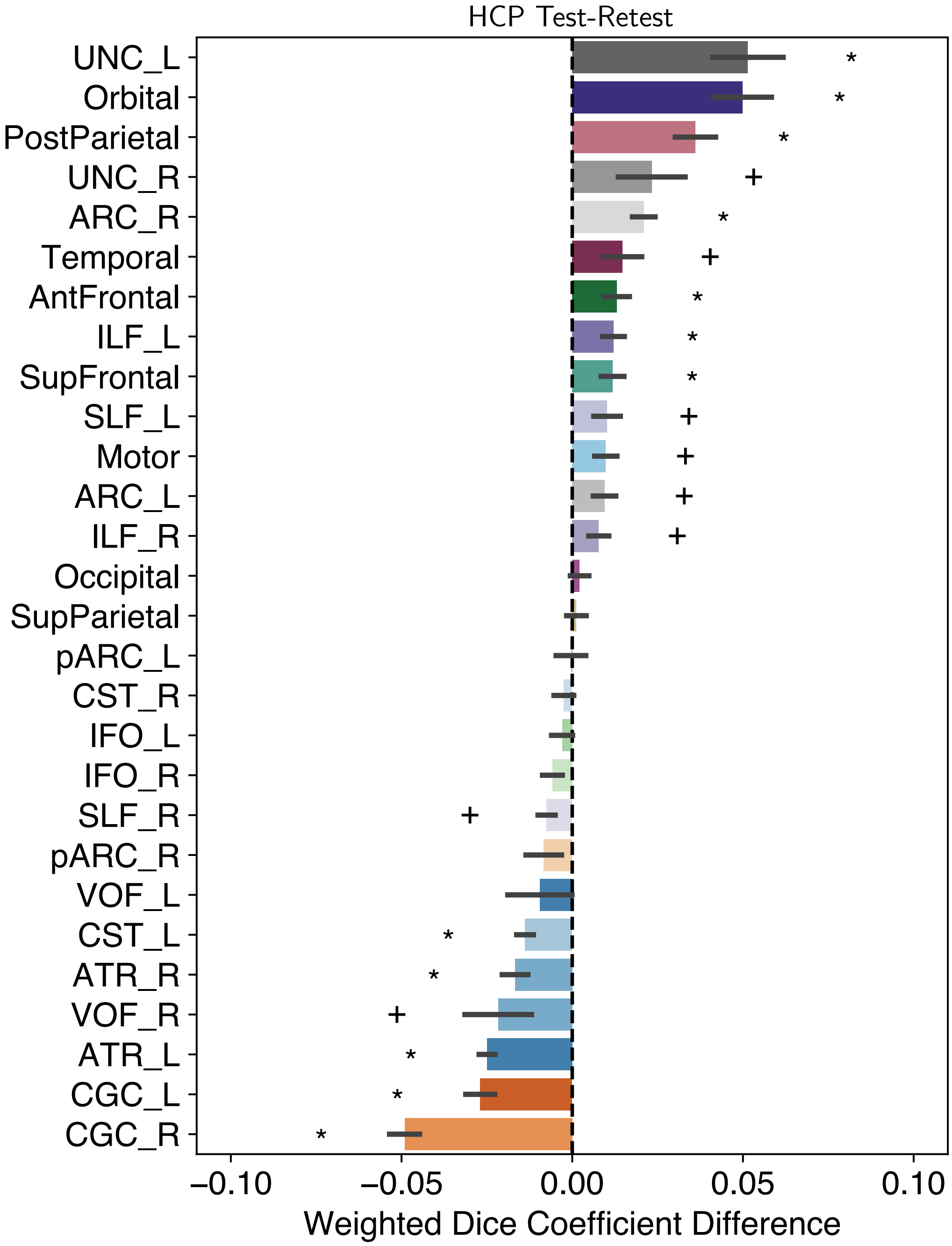


Figure 5: Tract weighted dice coefficient difference for the HCP Test-Retest dataset. The difference was calculated as FWE - Original. Error bars represent $\pm 1$ SEM. Asterisks represent tracts with weighted Dice coefficient differences significantly different (Bonferonni-corrected) from 0. Crosses represent tracts with weighted Dice coefficient difference (without Bonferonni correction) from 0.

##### 1.2.3 Tract profile reliability

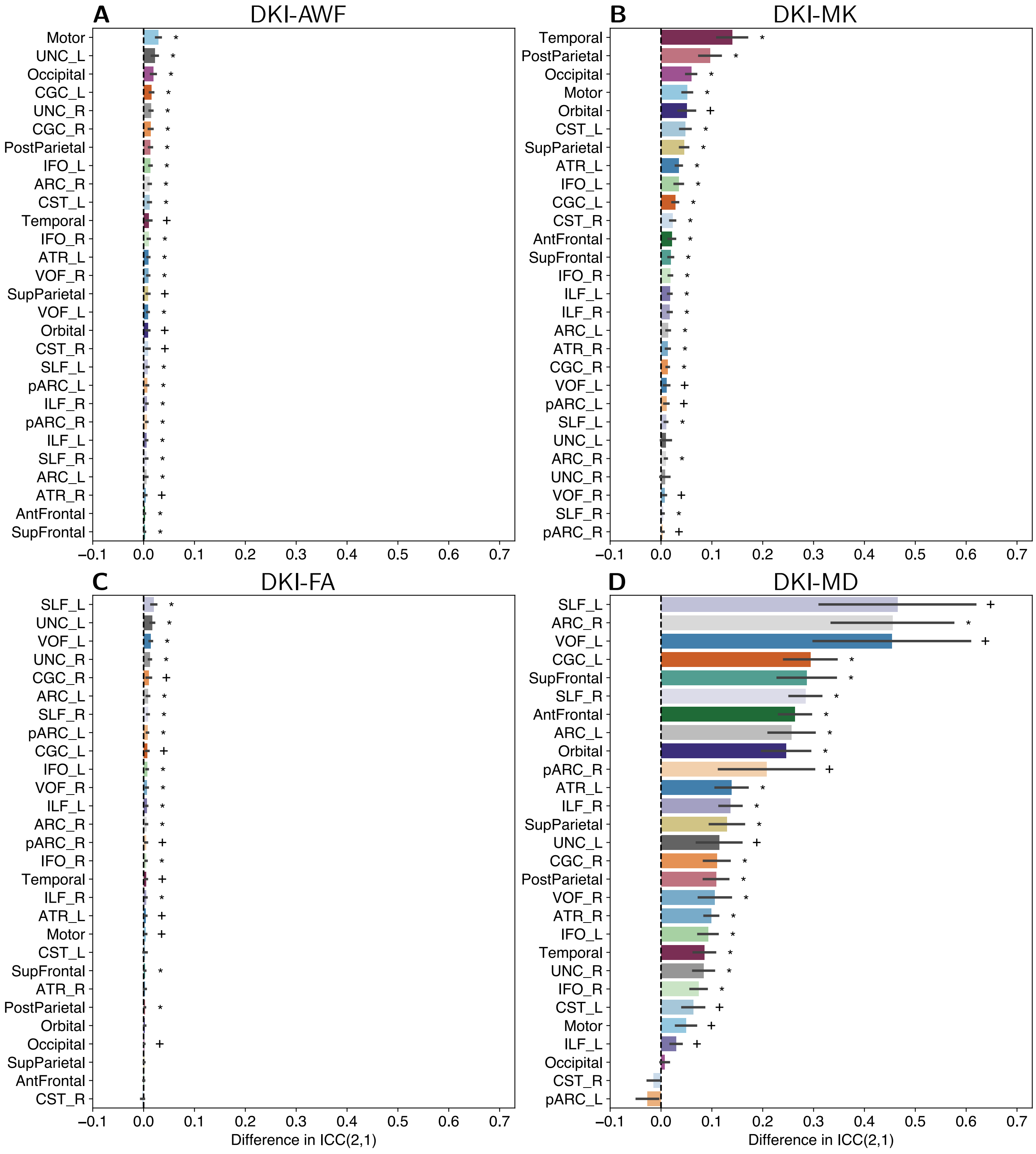


Figure 6: Differences in tract profile ICC(2,1) for the HCP Test-Retest dataset. Tract profile ICC(2,1) differences are shown for split-half multi-shell **(A)** DKI-AWF, **(B)** DKI-MD, **(C)** DKI-FA, and **(D)** DKI-MD. The difference was calculated as FWE - Original. Error bars represent $\pm 1$ SEM.

#### 1.3 Tract Profiles

We provide the tract profiles for each tract and metric from the multi-shell and single-shell datasets.

##### 1.3.1 Multi-shell dataset

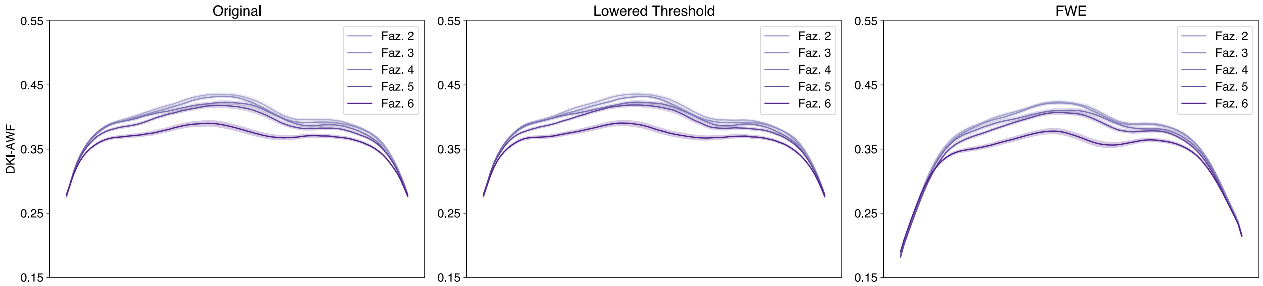


Figure 7: Multi-Shell Left Arcuate DKI-AWF profiles.

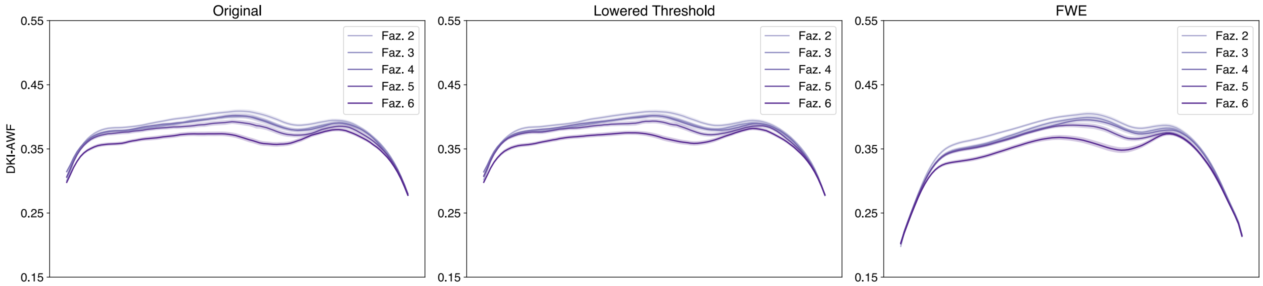


Figure 8: Multi-Shell Right Arcuate DKI-AWF profiles.

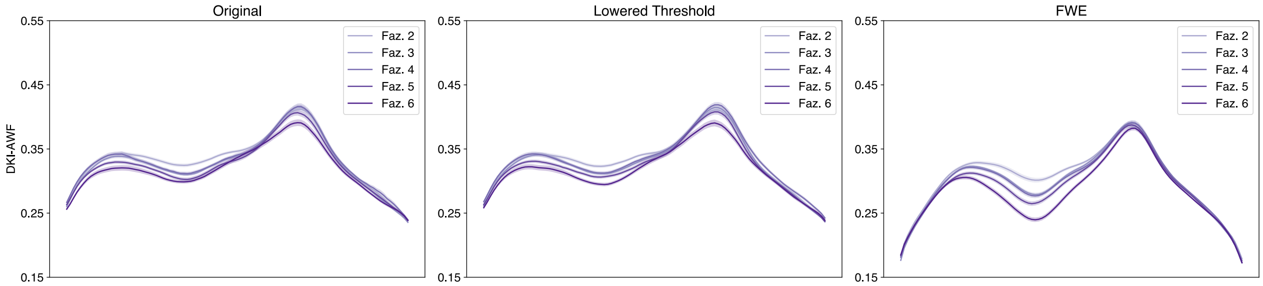


Figure 9: Multi-Shell Left Anterior Thalamic DKI-AWF profiles.

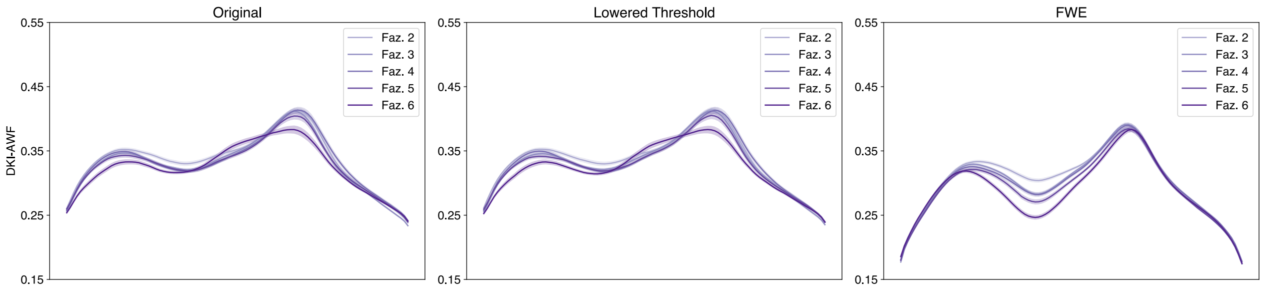


Figure 10: Multi-Shell Right Anterior Thalamic DKI-AWF profiles.

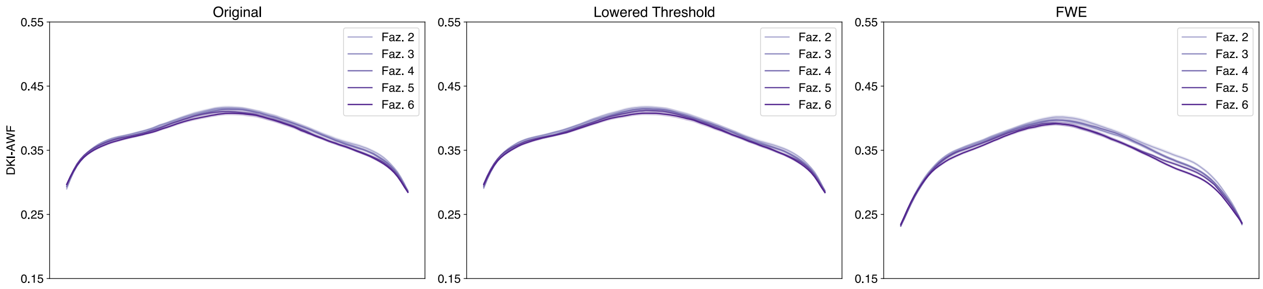


Figure 11: Multi-Shell Left Cingulum Cingulate DKI-AWF profiles.

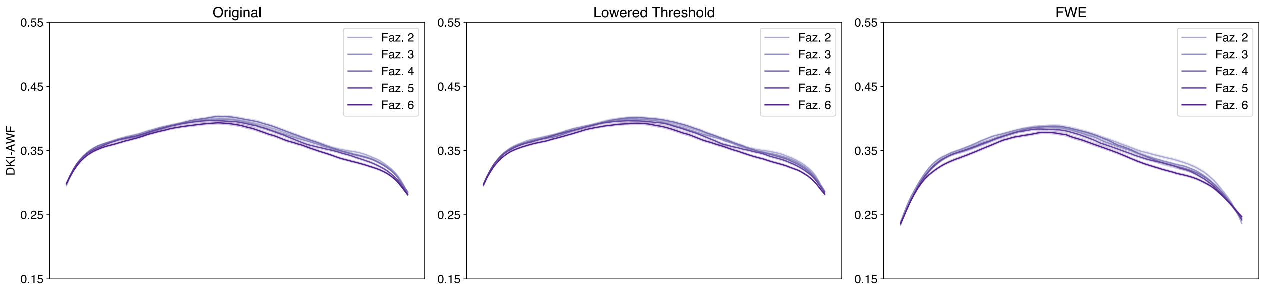


Figure 12: Multi-Shell Right Cingulum Cingulate DKI-AWF profiles.

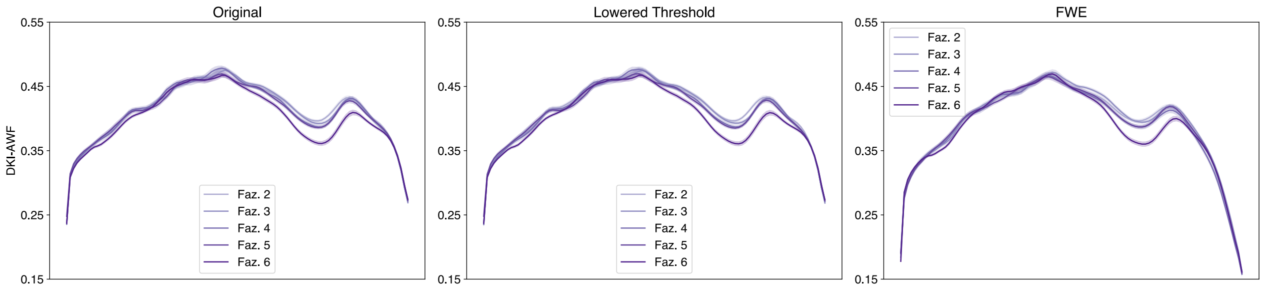


Figure 13: Multi-Shell Left Corticospinal DKI-AWF profiles.

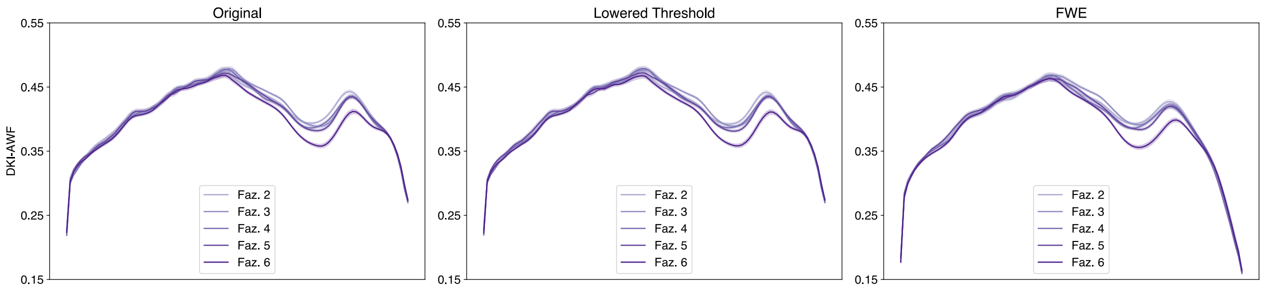


Figure 14: Multi-Shell Right Corticospinal DKI-AWF profiles.

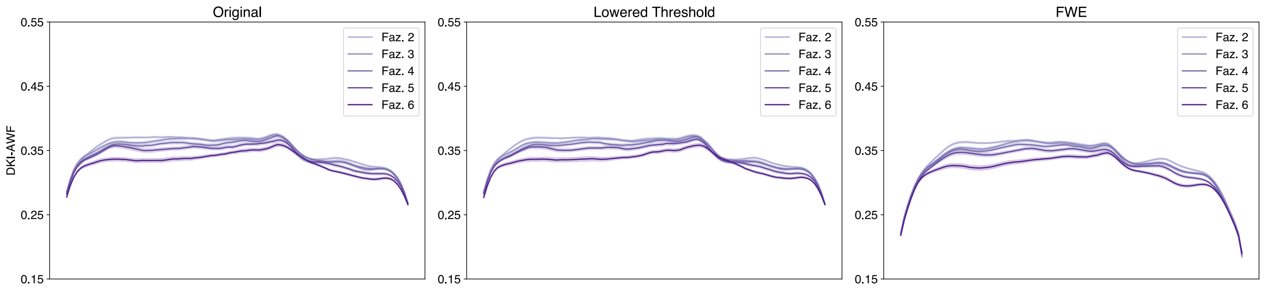


Figure 15: Multi-Shell Left Inferior Fronto-Occipital DKI-AWF profiles.

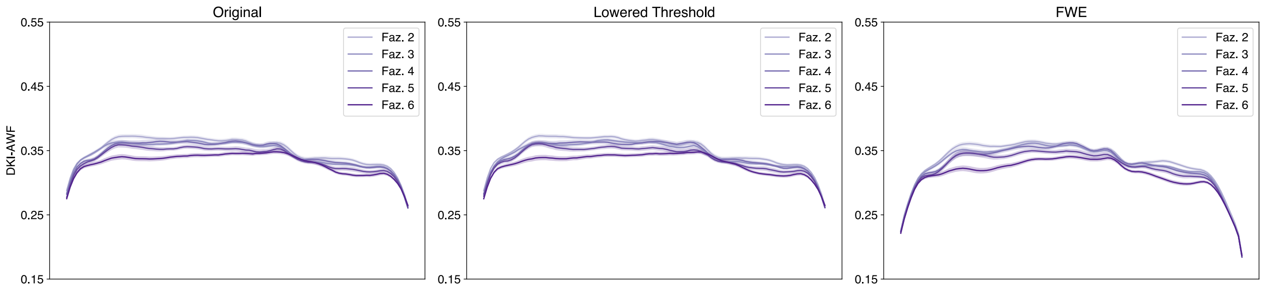


Figure 16: Multi-Shell Right Inferior Fronto-Occipital DKI-AWF profiles.

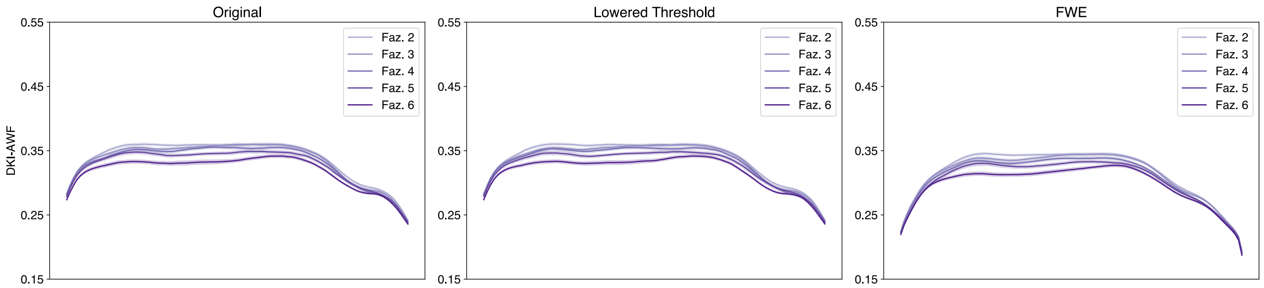


Figure 17: Multi-Shell Left Inferior Longitudinal DKI-AWF profiles.

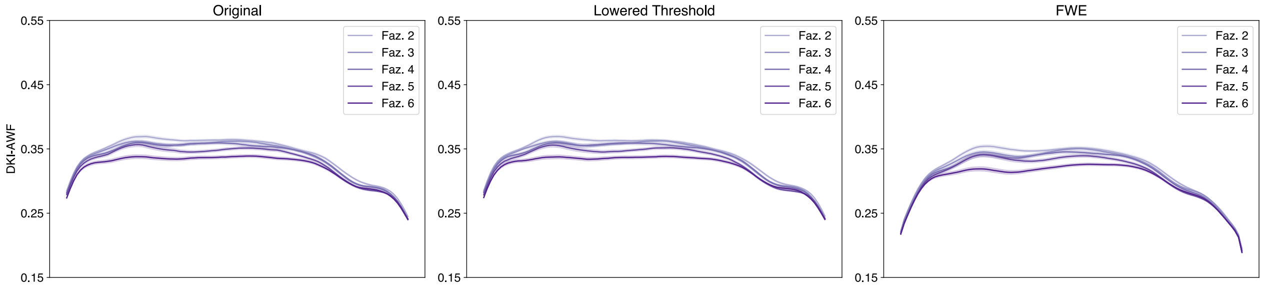


Figure 18: Multi-Shell Right Inferior Longitudinal DKI-AWF profiles.

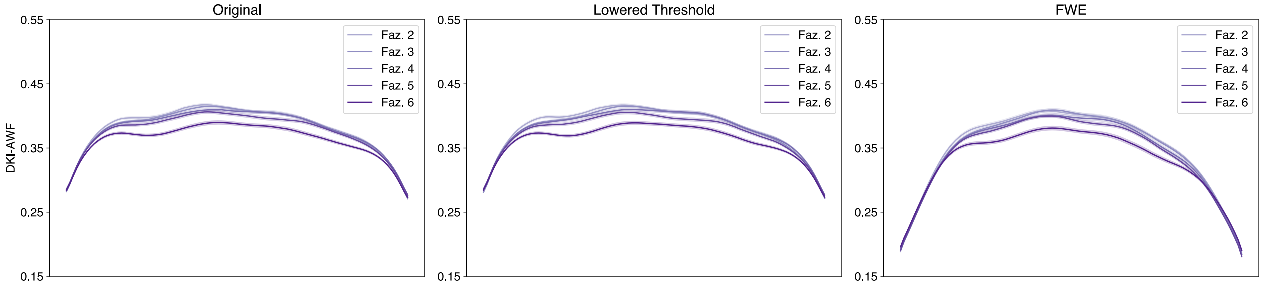


Figure 19: Multi-Shell Left Superior Longitudinal DKI-AWF profiles.

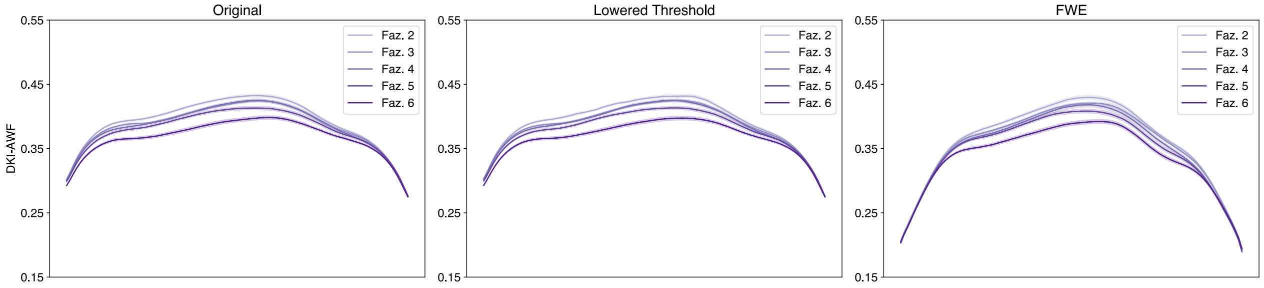


Figure 20: Multi-Shell Right Superior Longitudinal DKI-AWF profiles.

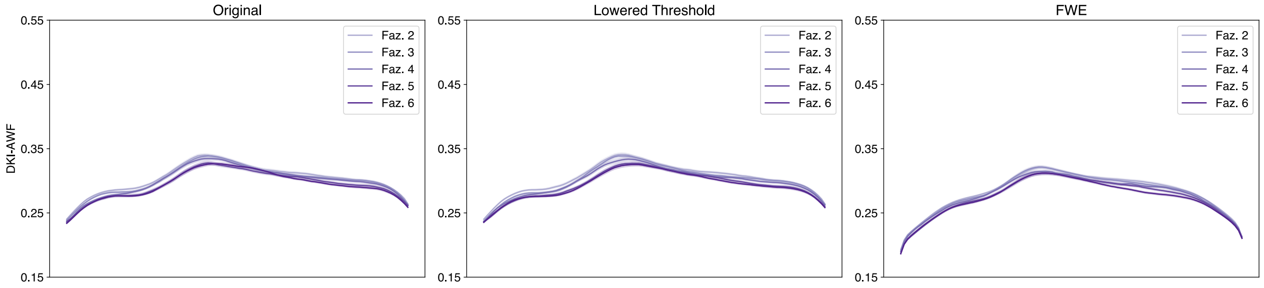


Figure 21: Multi-Shell Left Uncinate DKI-AWF profiles.

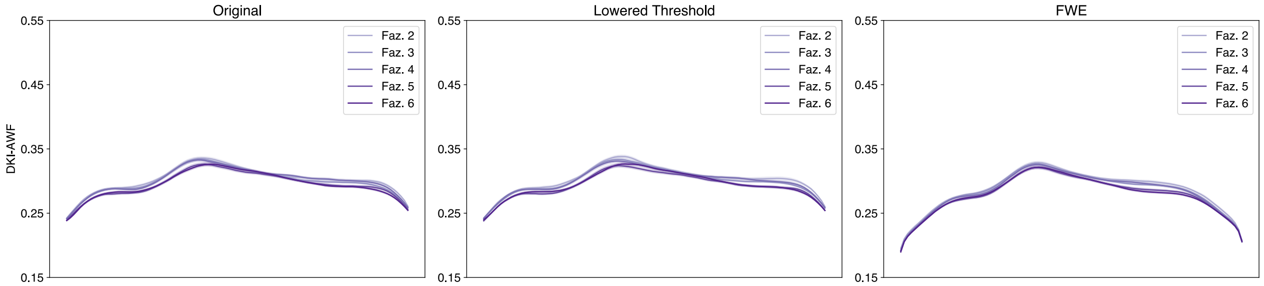


Figure 22: Multi-Shell Right Uncinate DKI-AWF profiles.

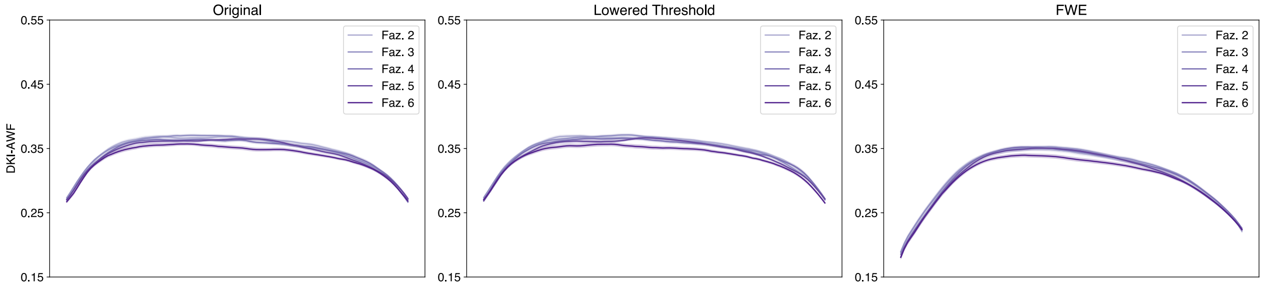


Figure 23: Multi-Shell Left Vertical Occipital DKI-AWF profiles.

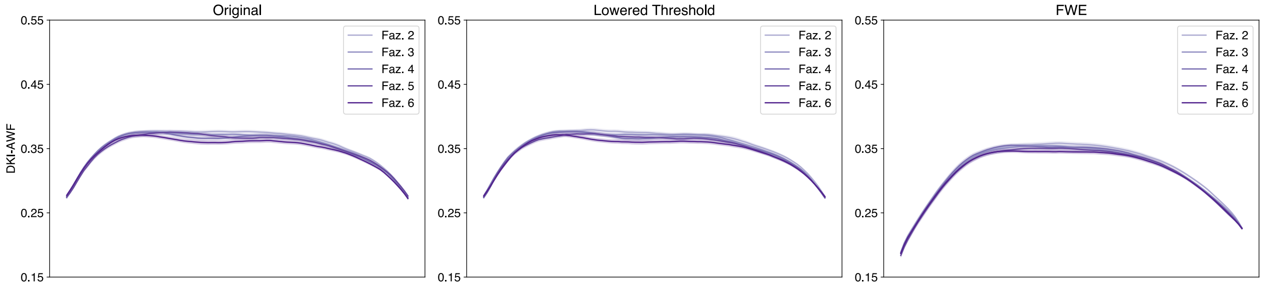


Figure 24: Multi-Shell Right Vertical Occipital DKI-AWF profiles.

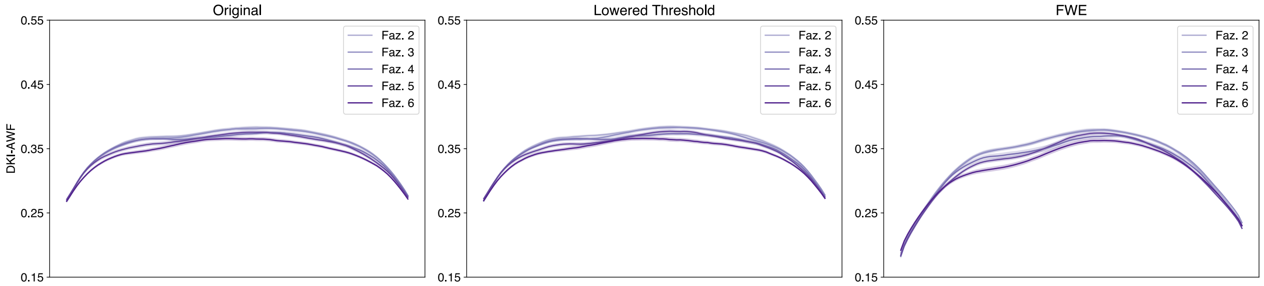


Figure 25: Multi-Shell Left Posterior Arcuate DKI-AWF profiles.

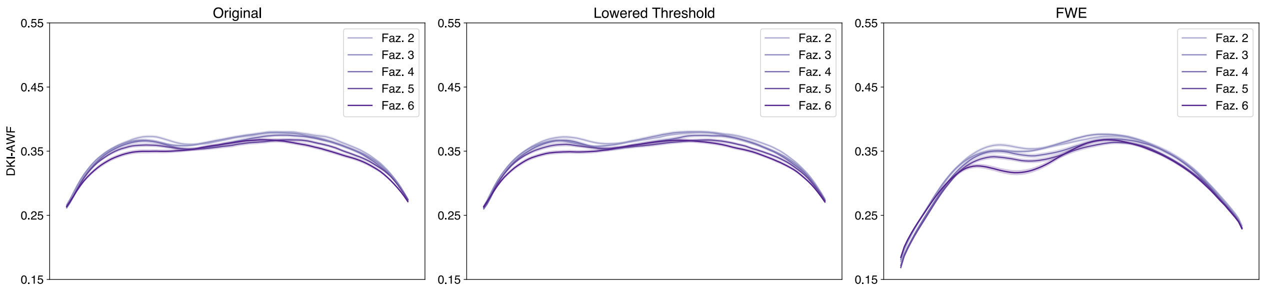


Figure 26: Multi-Shell Right Posterior Arcuate DKI-AWF profiles.

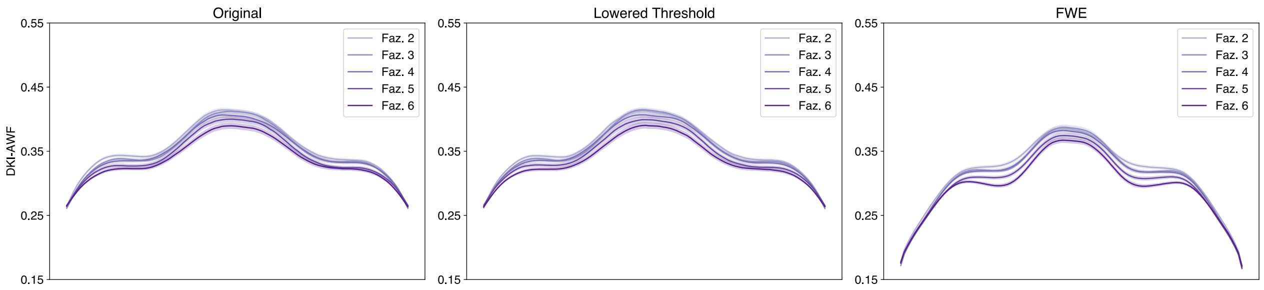


Figure 27: Multi-Shell Anterior Frontal Callosum DKI-AWF profiles.

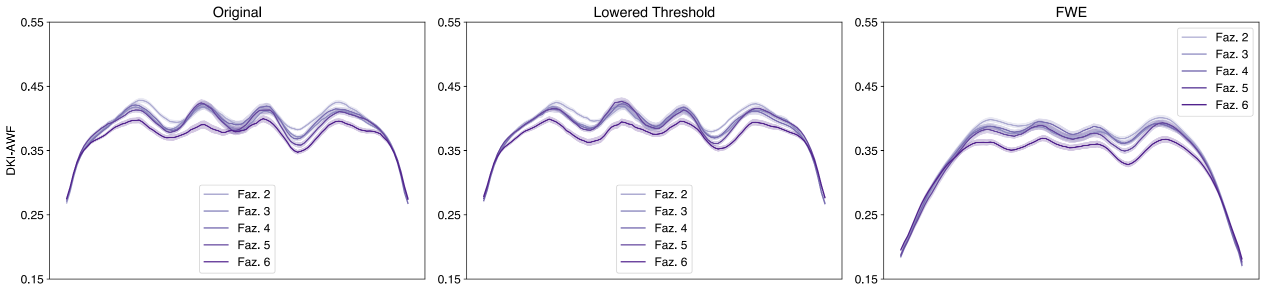


Figure 28: Multi-Shell Motor Corpus Callosum DKI-AWF profiles.

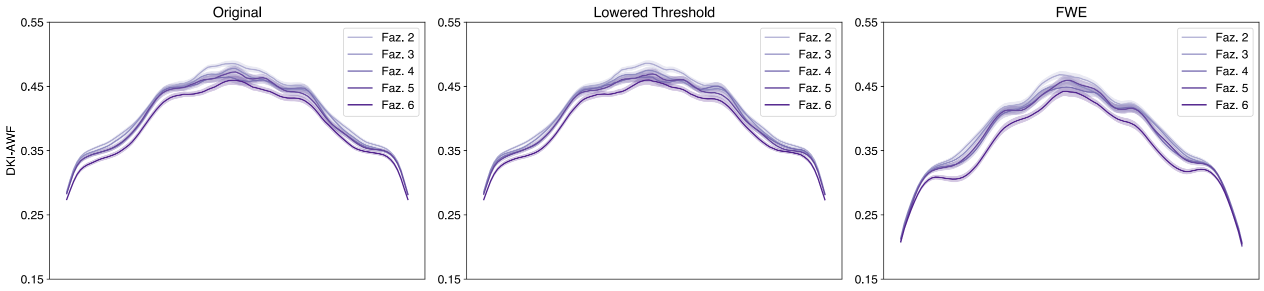


Figure 29: Multi-Shell Occipital Corpus Callosum DKI-AWF profiles.

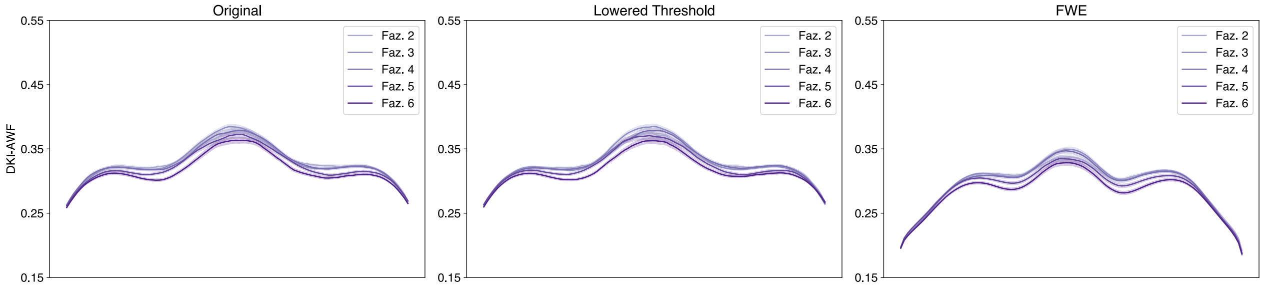


Figure 30: Multi-Shell Orbital Corpus Callosum DKI-AWF profiles.

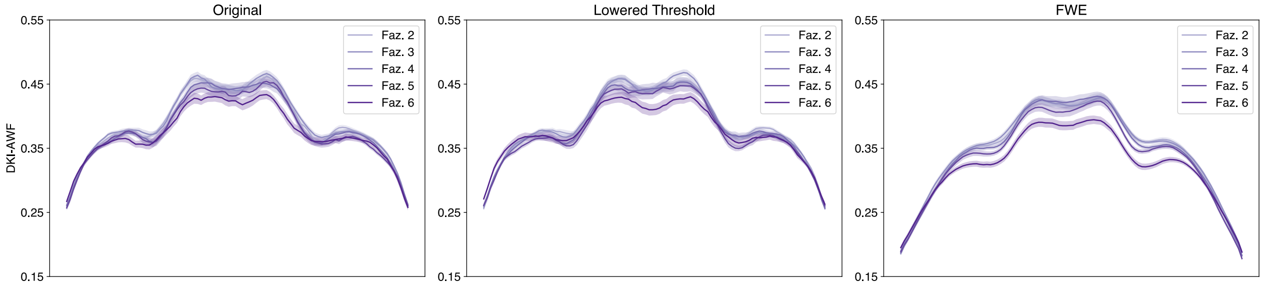


Figure 31: Multi-Shell Posterior Parietal Corpus Callosum DKI-AWF profiles.

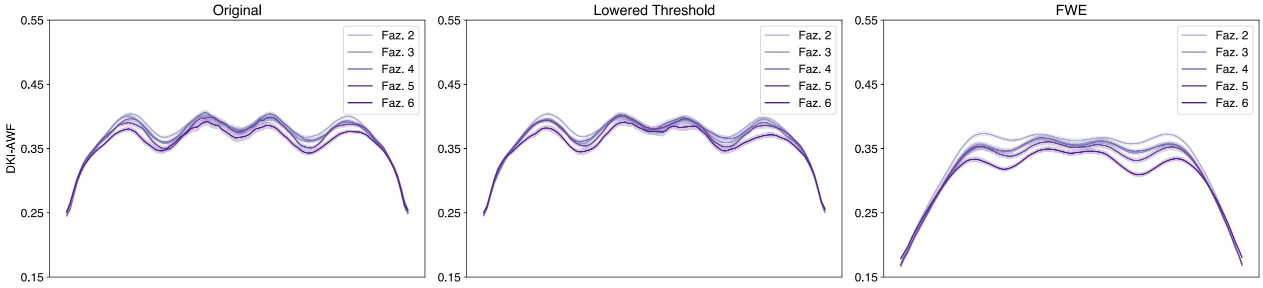


Figure 32: Multi-Shell Superior Frontal Callosum DKI-AWF profiles.

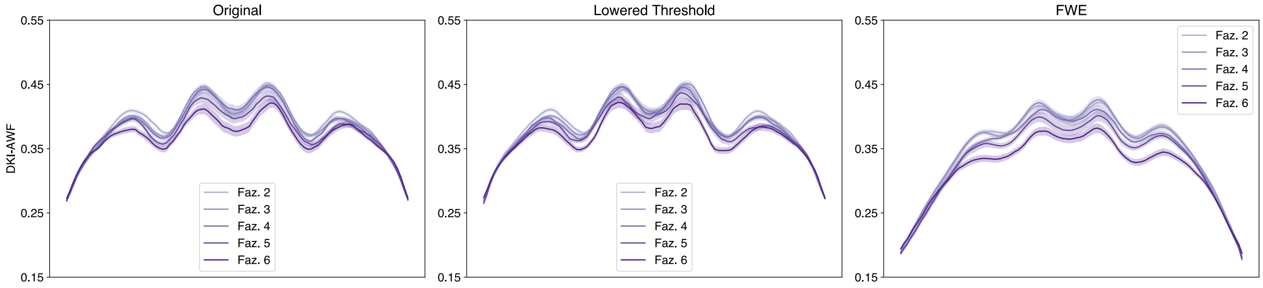


Figure 33: Multi-Shell Superior Parietal Corpus Callosum DKI-AWF profiles.

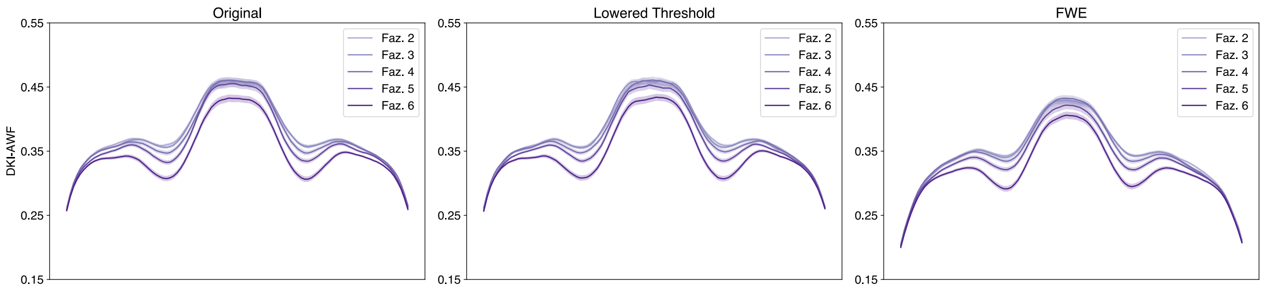


Figure 34: Multi-Shell Temporal Corpus Callosum DKI-AWF profiles.

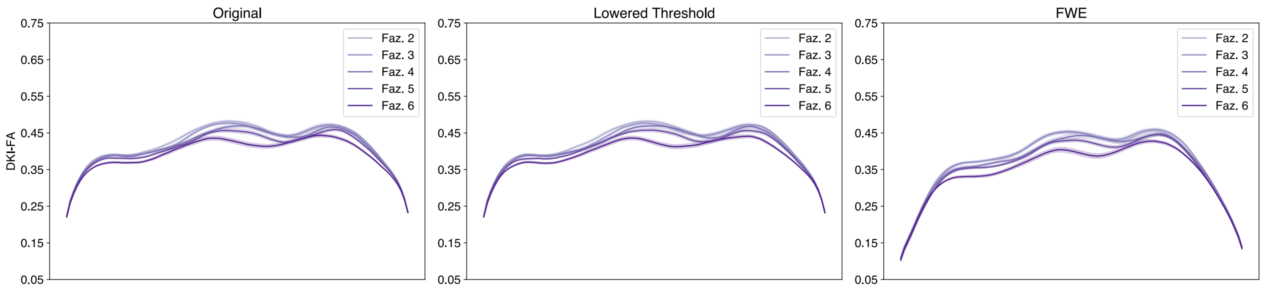


Figure 35: Multi-Shell Left Arcuate DKI-FA profiles.

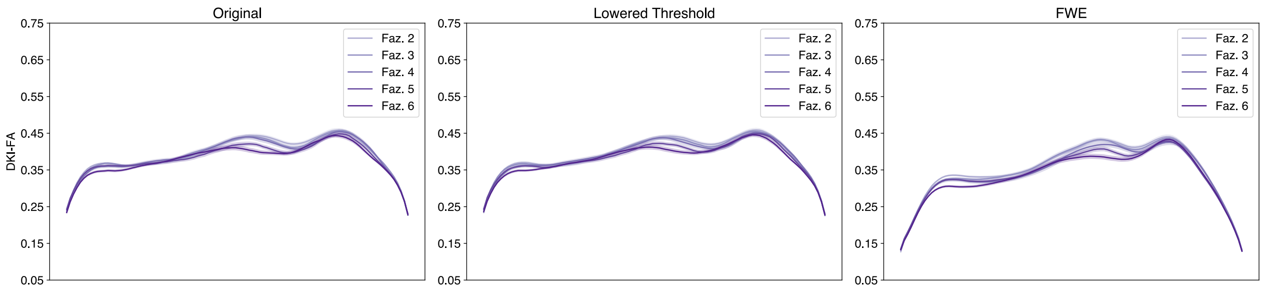


Figure 36: Multi-Shell Right Arcuate DKI-FA profiles.

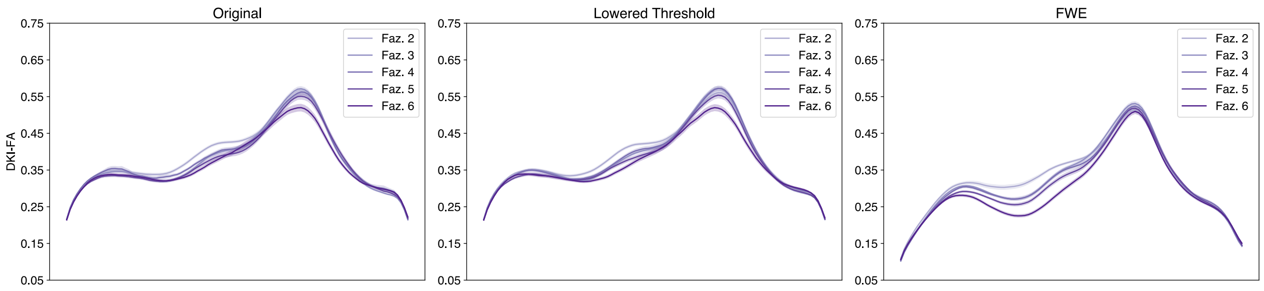


Figure 37: Multi-Shell Left Anterior Thalamic DKI-FA profiles.

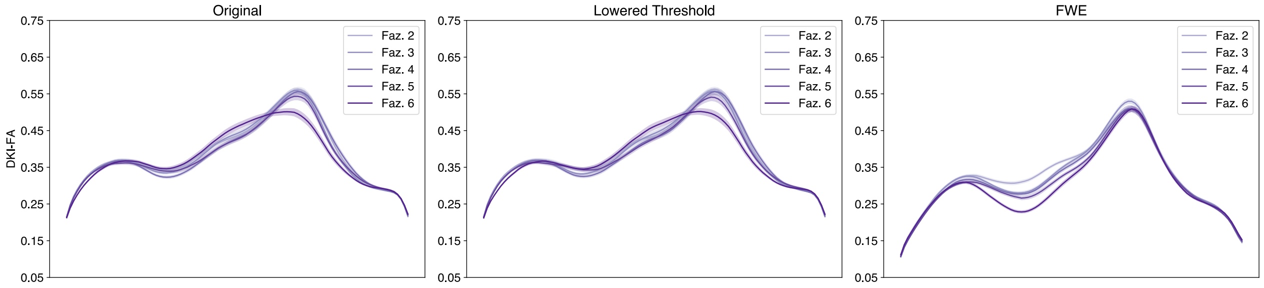


Figure 38: Multi-Shell Right Anterior Thalamic DKI-FA profiles.

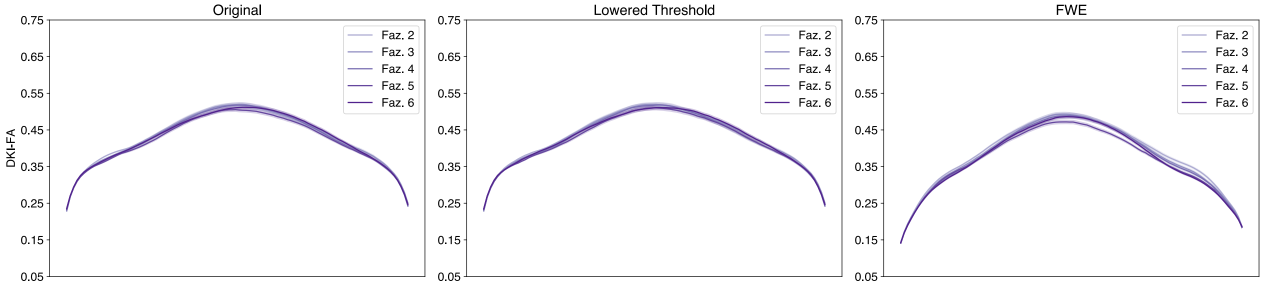


Figure 39: Multi-Shell Left Cingulum Cingulate DKI-FA profiles.

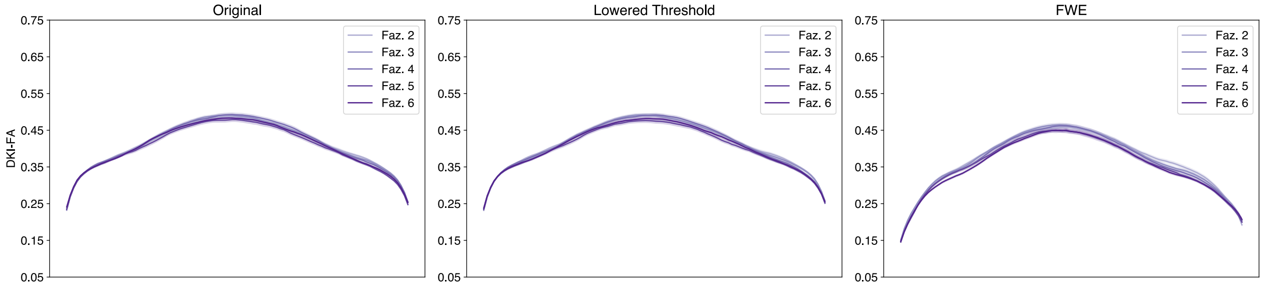


Figure 40: Multi-Shell Right Cingulum Cingulate DKI-FA profiles.

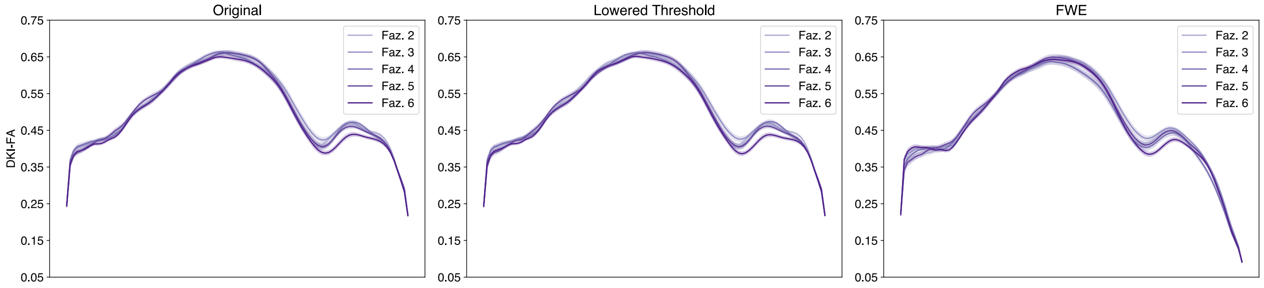


Figure 41: Multi-Shell Left Corticospinal DKI-FA profiles.

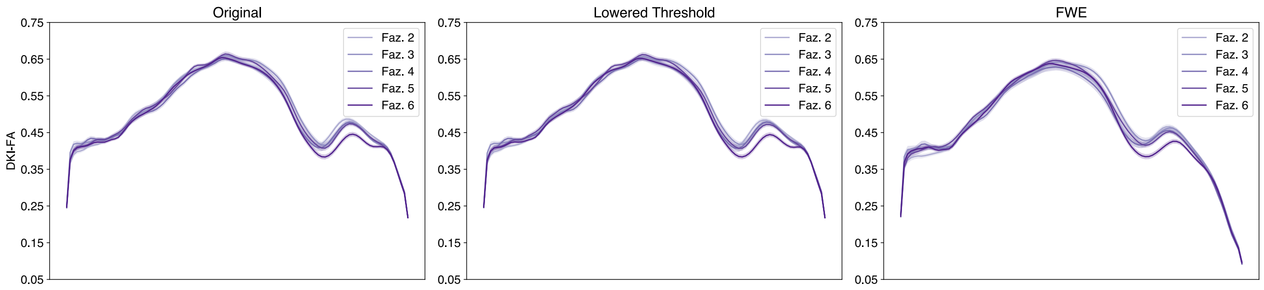


Figure 42: Multi-Shell Right Corticospinal DKI-FA profiles.

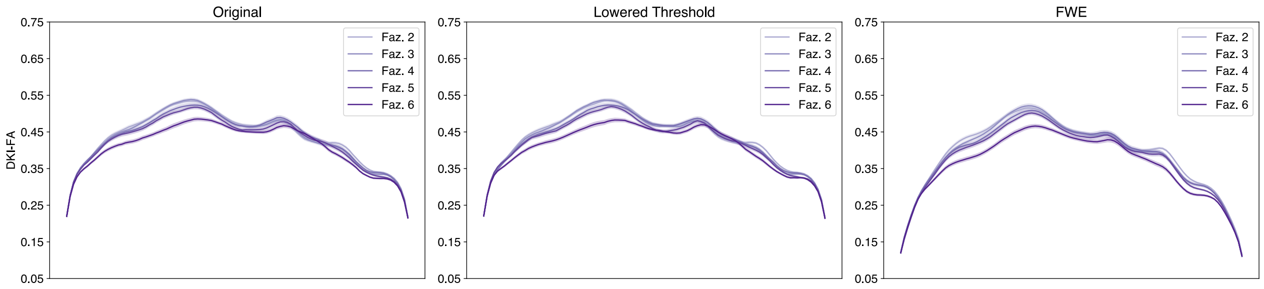


Figure 43: Multi-Shell Left Inferior Fronto-Occipital DKI-FA profiles.

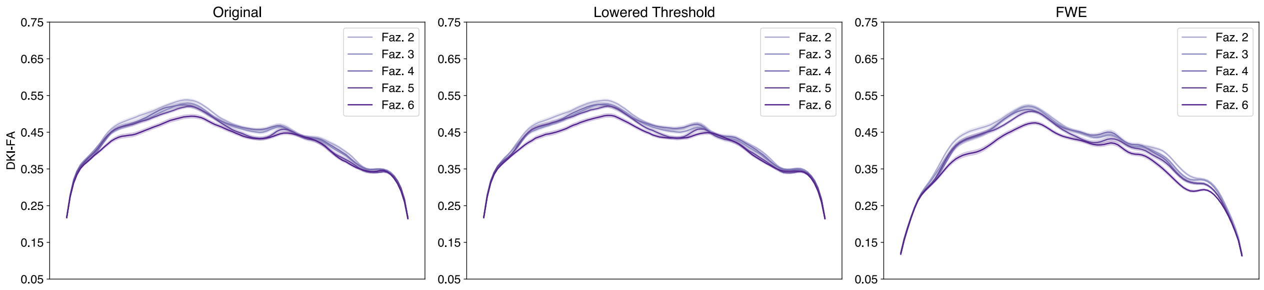


Figure 44: Multi-Shell Right Inferior Fronto-Occipital DKI-FA profiles.

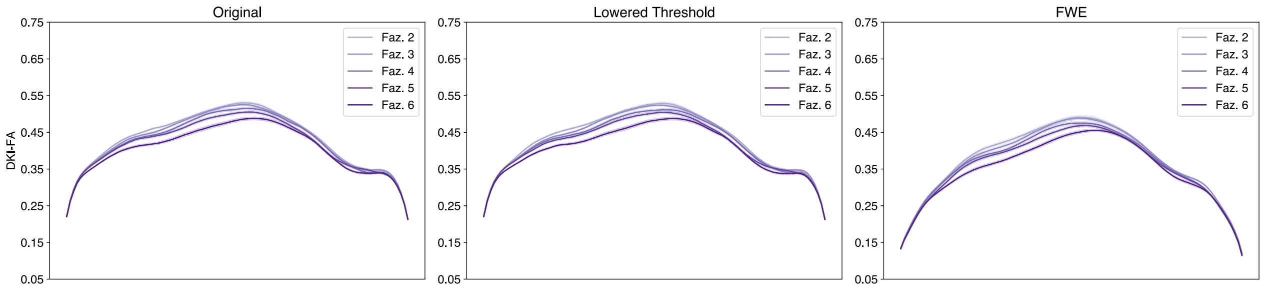


Figure 45: Multi-Shell Left Inferior Longitudinal DKI-FA profiles.

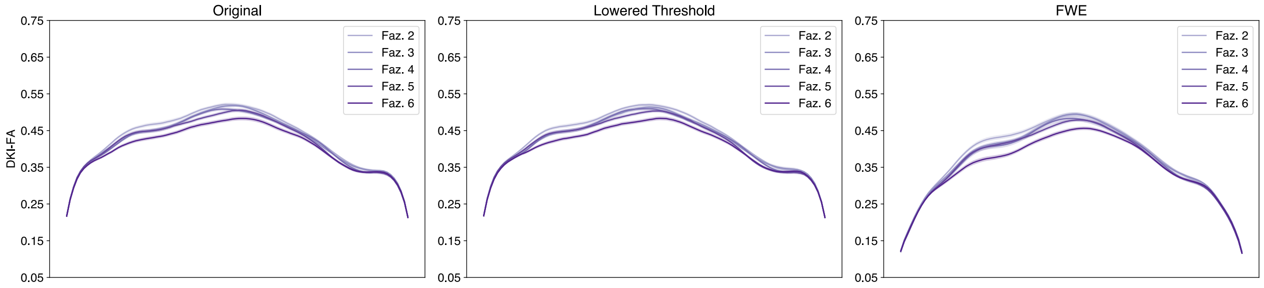


Figure 46: Multi-Shell Right Inferior Longitudinal DKI-FA profiles.

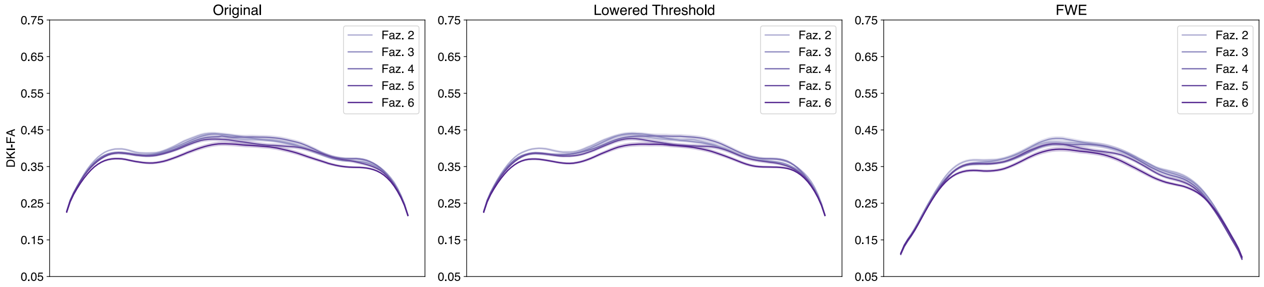


Figure 47: Multi-Shell Left Superior Longitudinal DKI-FA profiles.

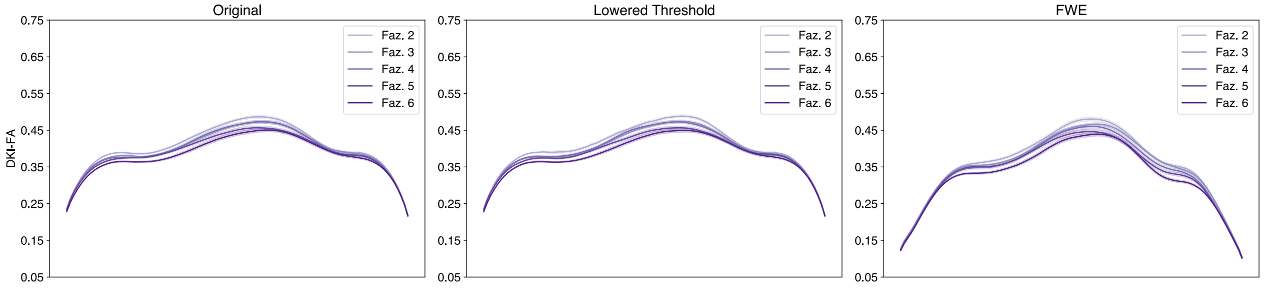


Figure 48: Multi-Shell Right Superior Longitudinal DKI-FA profiles.

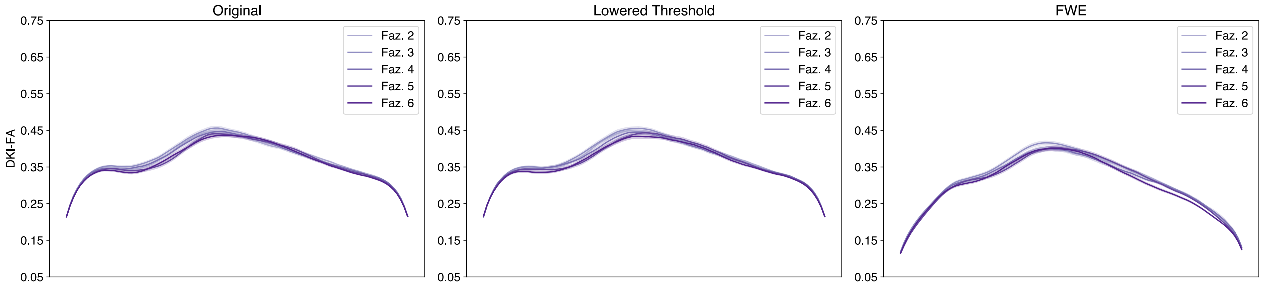


Figure 49: Multi-Shell Left Uncinate DKI-FA profiles.

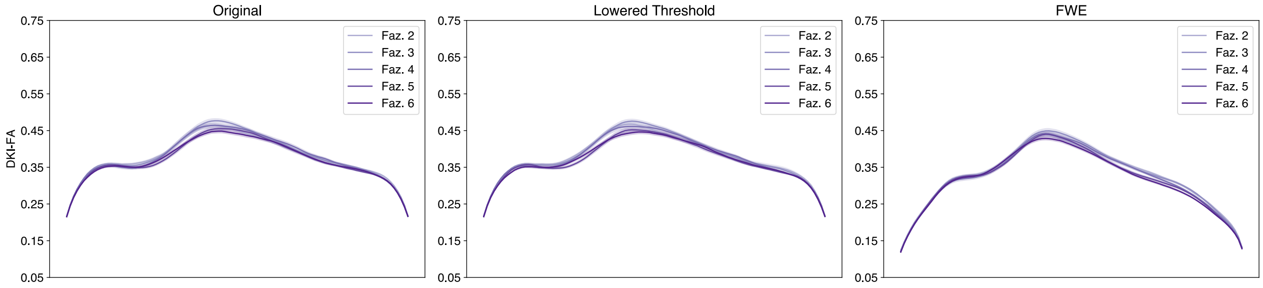

Figure 50: Multi-Shell Right Uncinate DKI-FA profiles.

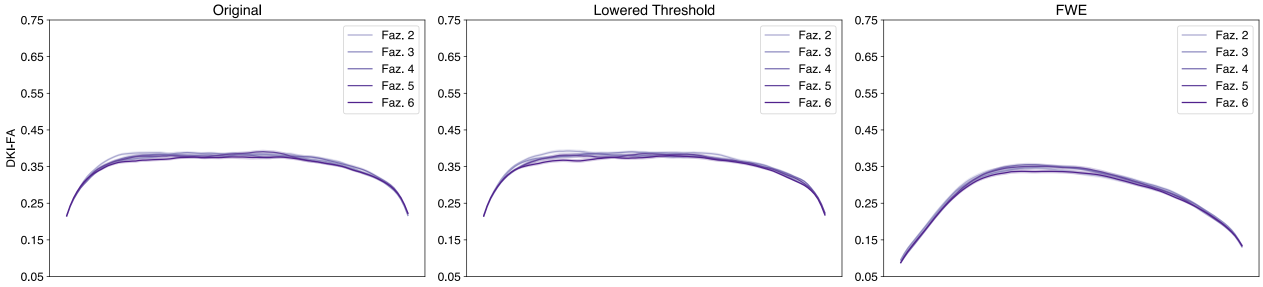

Figure 51: Multi-Shell Left Vertical Occipital DKI-FA profiles.

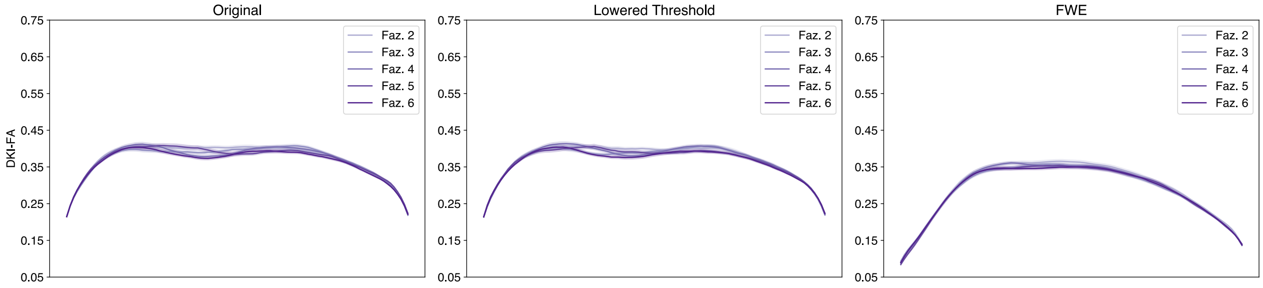

Figure 52: Multi-Shell Right Vertical Occipital DKI-FA profiles.

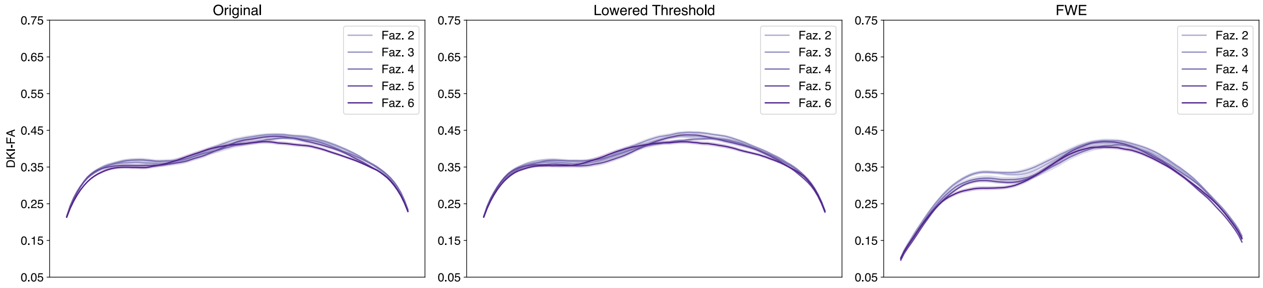

Figure 53: Multi-Shell Left Posterior Arcuate DKI-FA profiles.

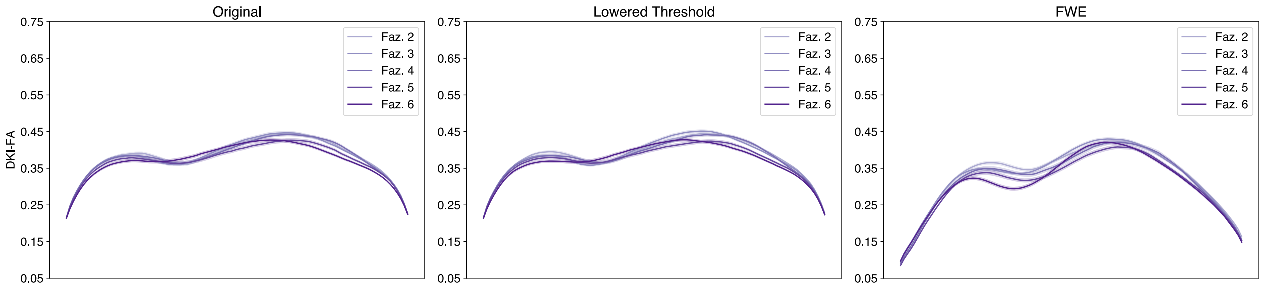


Figure 54: Multi-Shell Right Posterior Arcuate DKI-FA profiles.

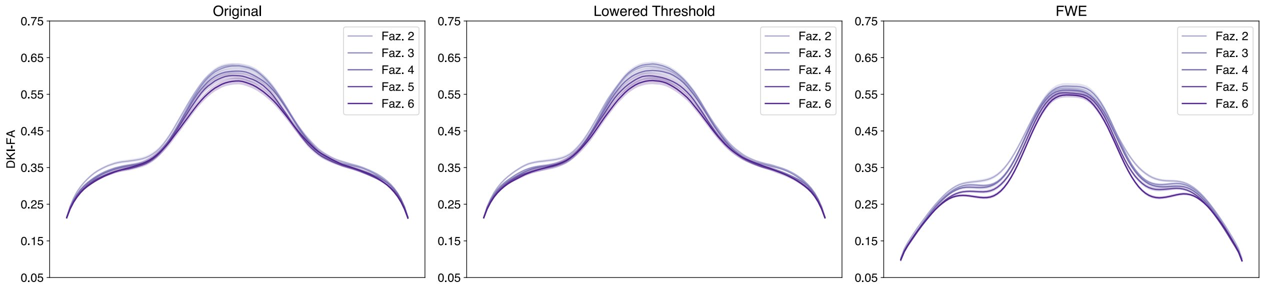


Figure 55: Multi-Shell Anterior Frontal Callosum DKI-FA profiles.

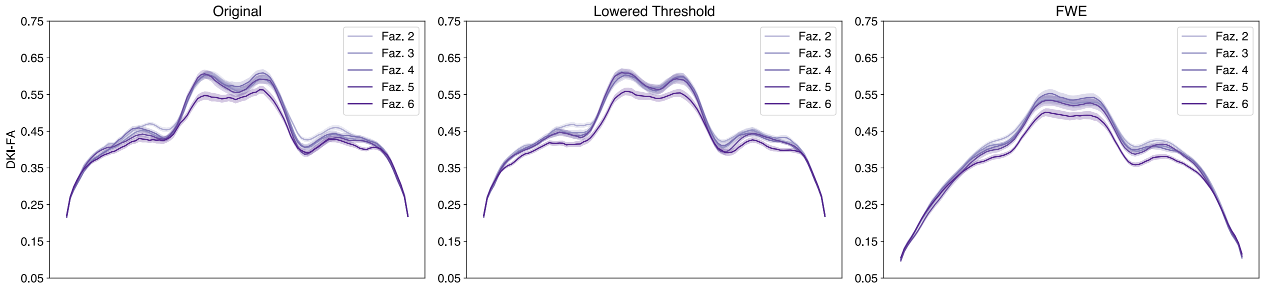


Figure 56: Multi-Shell Motor Corpus Callosum DKI-FA profiles.

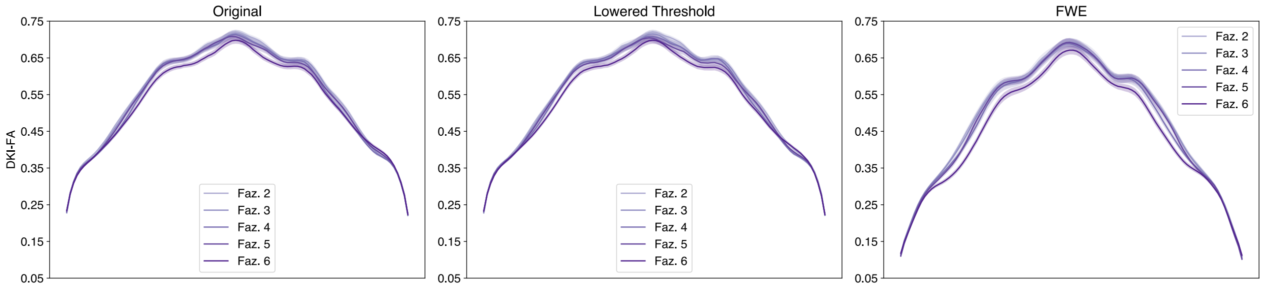


Figure 57: Multi-Shell Occipital Corpus Callosum DKI-FA profiles.

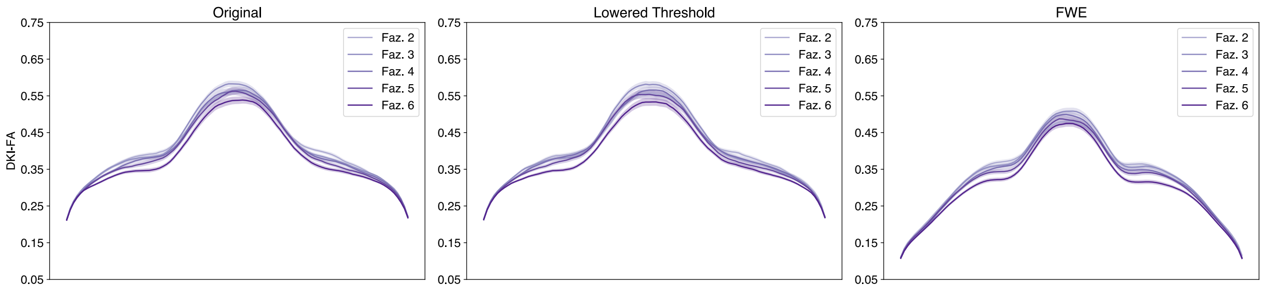


Figure 58: Multi-Shell Orbital Corpus Callosum DKI-FA profiles.

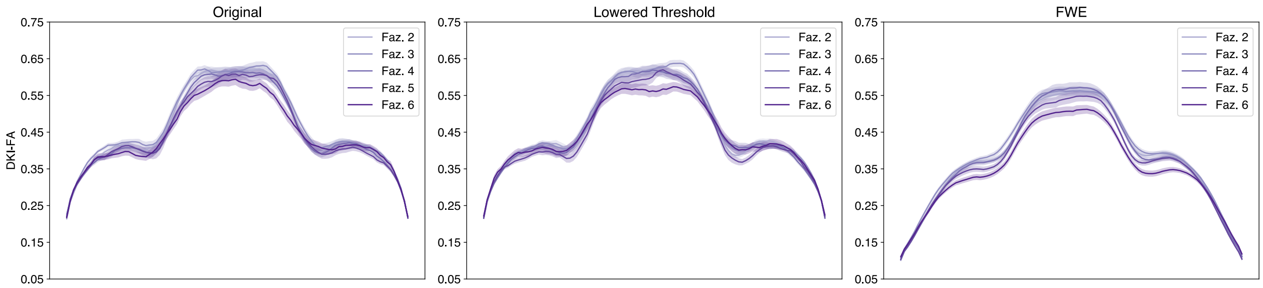


Figure 59: Multi-Shell Posterior Parietal Corpus Callosum DKI-FA profiles.

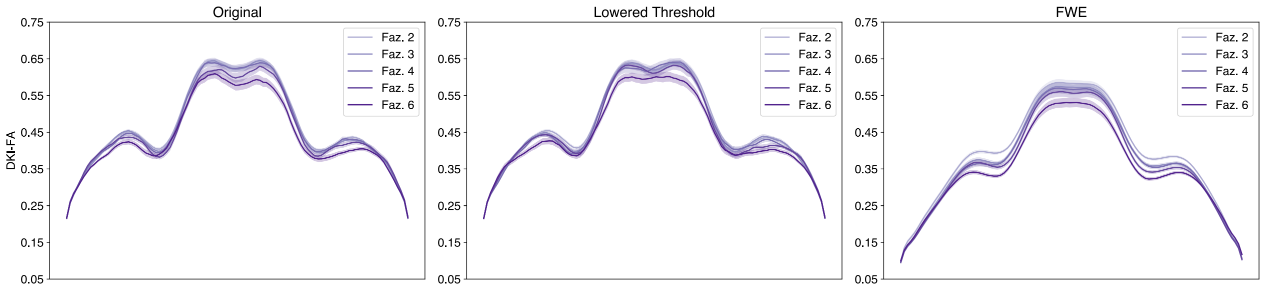


Figure 60: Multi-Shell Superior Frontal Callosum DKI-FA profiles.

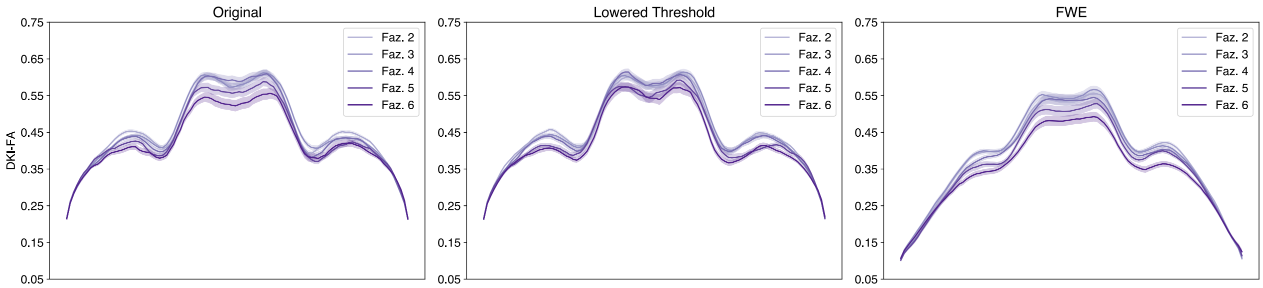


Figure 61: Multi-Shell Superior Parietal Corpus Callosum DKI-FA profiles.

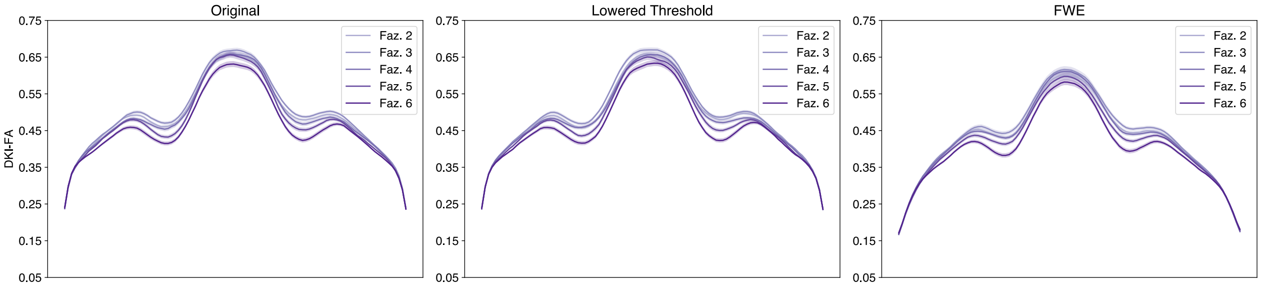


Figure 62: Multi-Shell Temporal Corpus Callosum DKI-FA profiles.

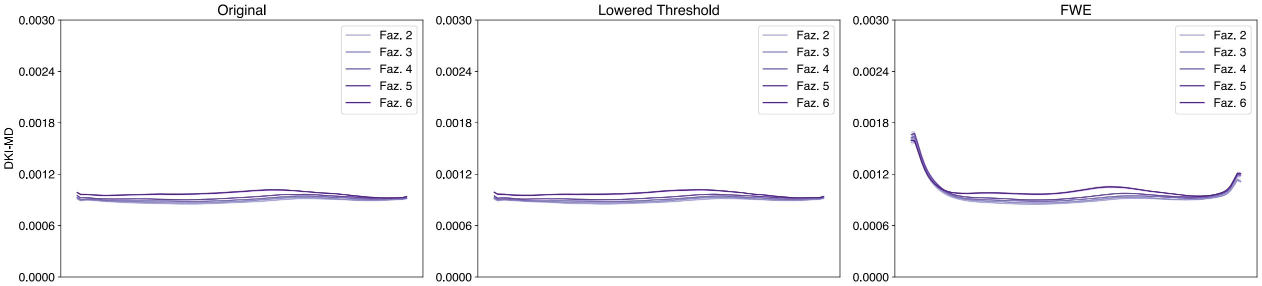


Figure 63: Multi-Shell Left Arcuate DKI-MD profiles.

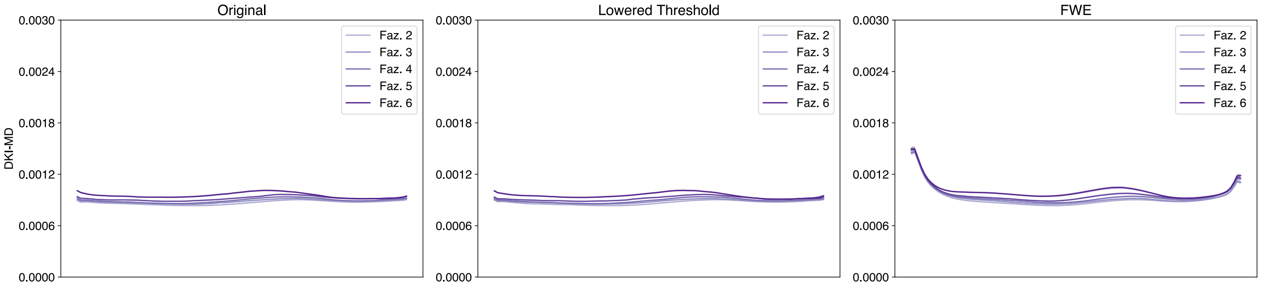


Figure 64: Multi-Shell Right Arcuate DKI-MD profiles.

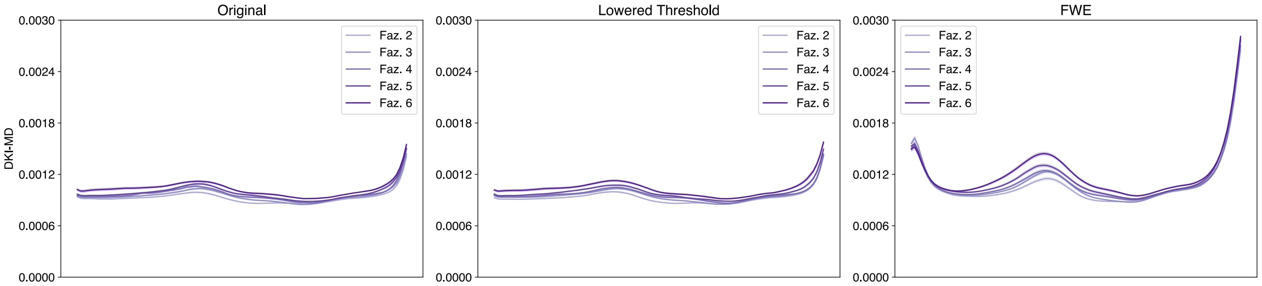


Figure 65: Multi-Shell Left Anterior Thalamic DKI-MD profiles.

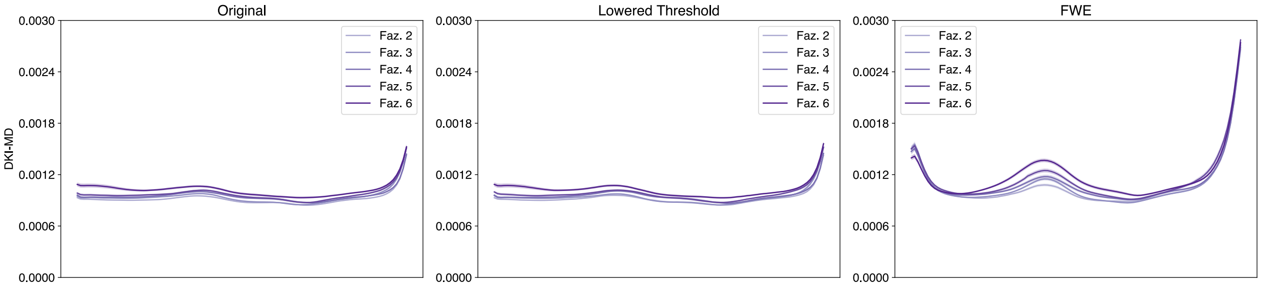


Figure 66: Multi-Shell Right Anterior Thalamic DKI-MD profiles.

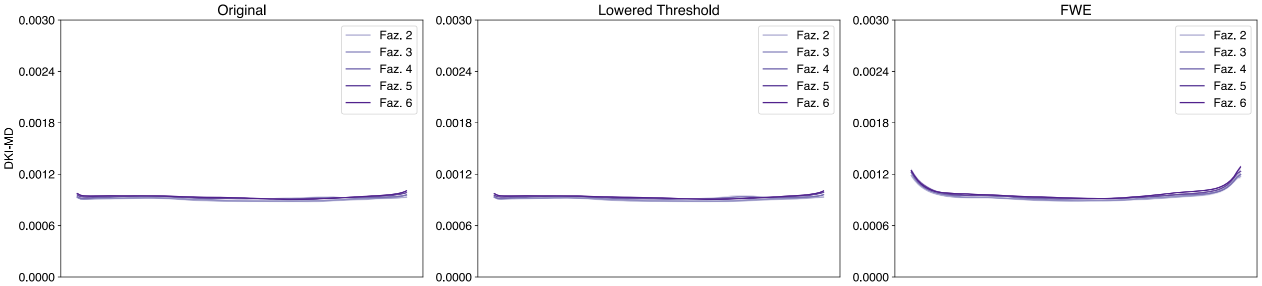


Figure 67: Multi-Shell Left Cingulum Cingulate DKI-MD profiles.

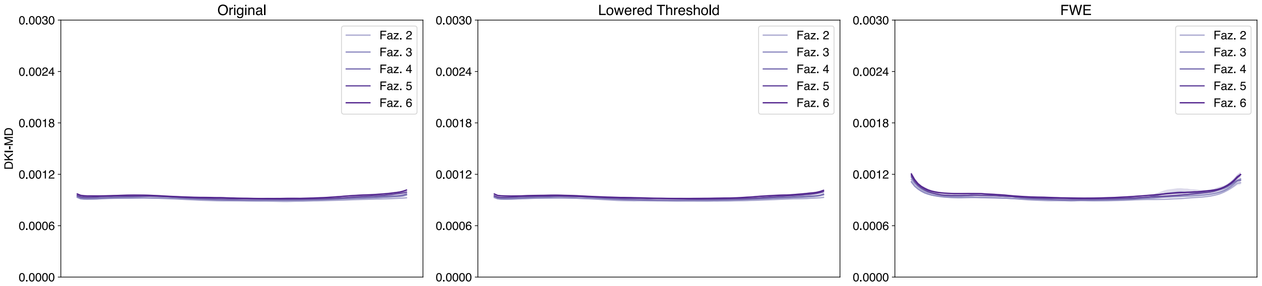


Figure 68: Multi-Shell Right Cingulum Cingulate DKI-MD profiles.

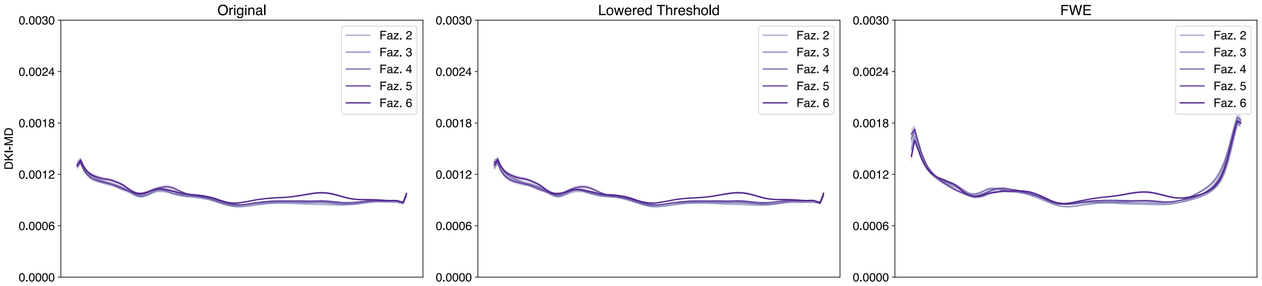


Figure 69: Multi-Shell Left Corticospinal DKI-MD profiles.

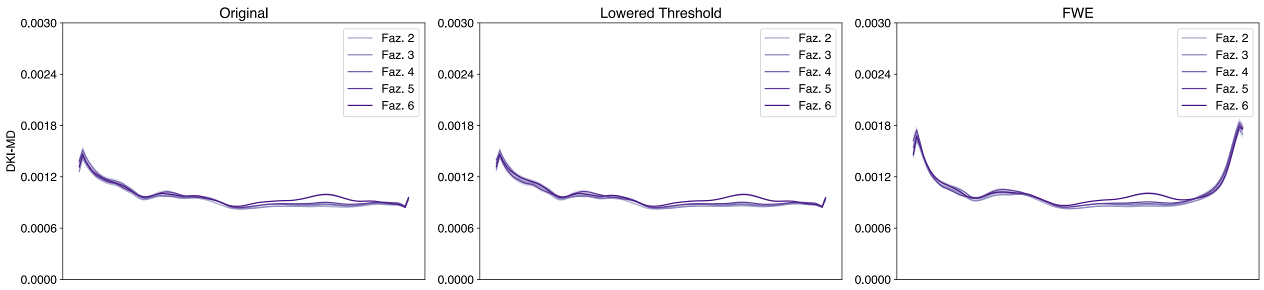


Figure 70: Multi-Shell Right Corticospinal DKI-MD profiles.

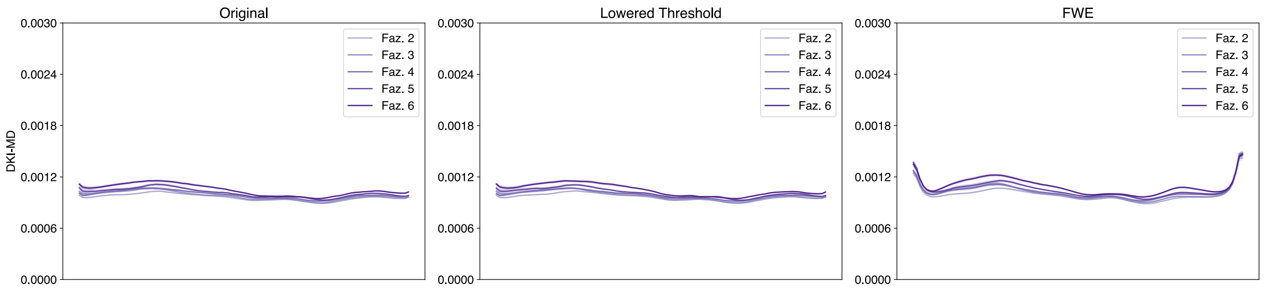


Figure 71: Multi-Shell Left Inferior Fronto-Occipital DKI-MD profiles.

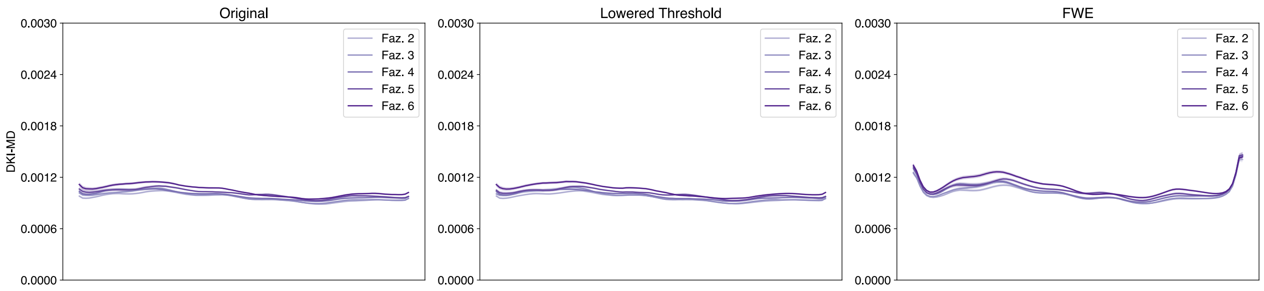


Figure 72: Multi-Shell Right Inferior Fronto-Occipital DKI-MD profiles.

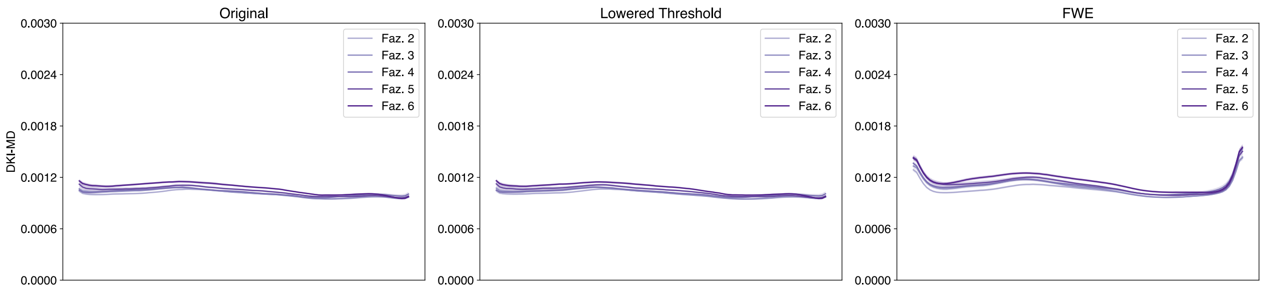


Figure 73: Multi-Shell Left Inferior Longitudinal DKI-MD profiles.

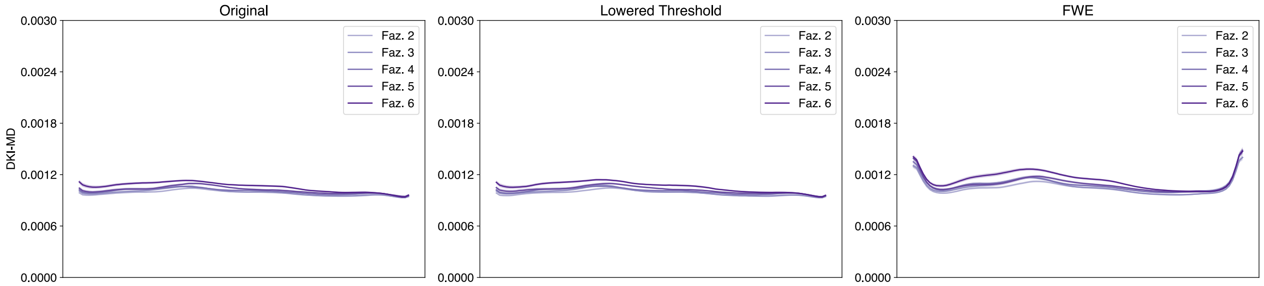


Figure 74: Multi-Shell Right Inferior Longitudinal DKI-MD profiles.

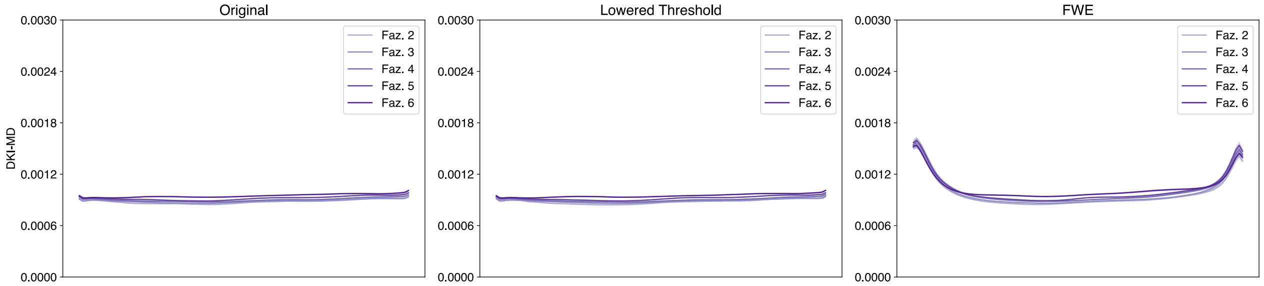


Figure 75: Multi-Shell Left Superior Longitudinal DKI-MD profiles.

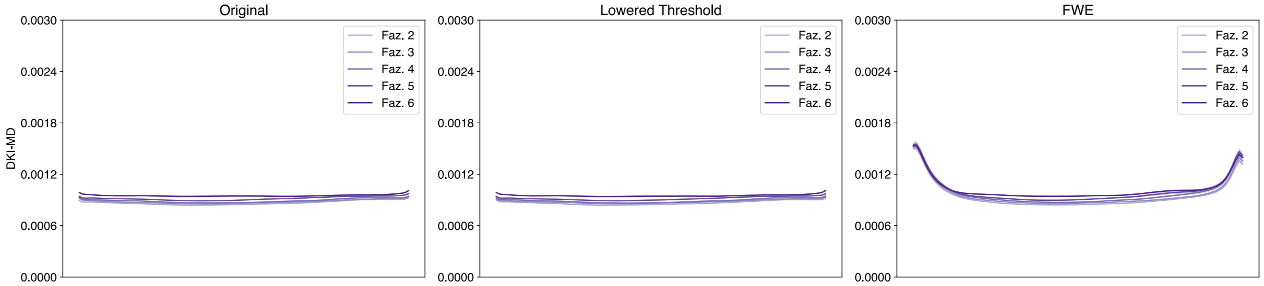


Figure 76: Multi-Shell Right Superior Longitudinal DKI-MD profiles.

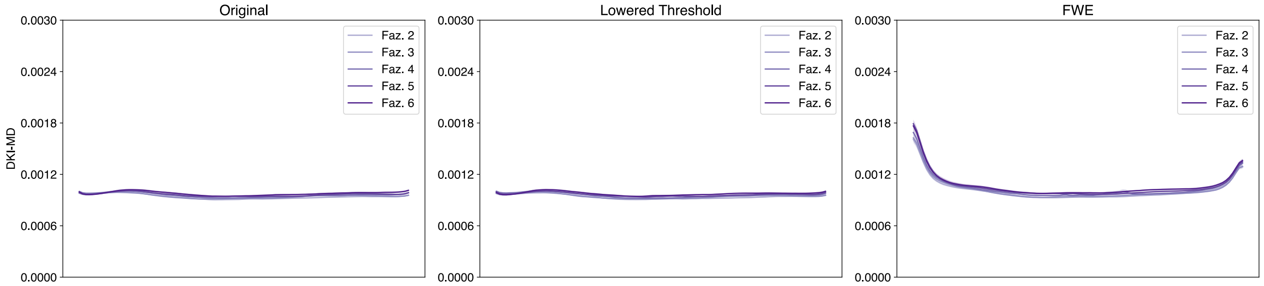


Figure 77: Multi-Shell Left Uncinate DKI-MD profiles.

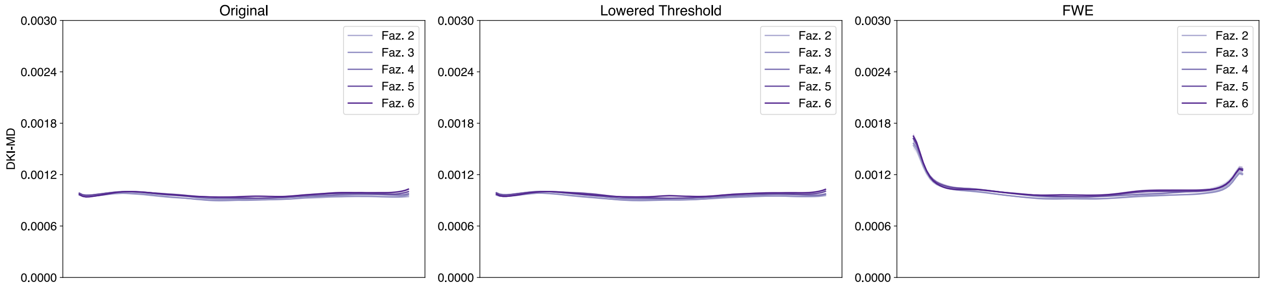


Figure 78: Multi-Shell Right Uncinate DKI-MD profiles.

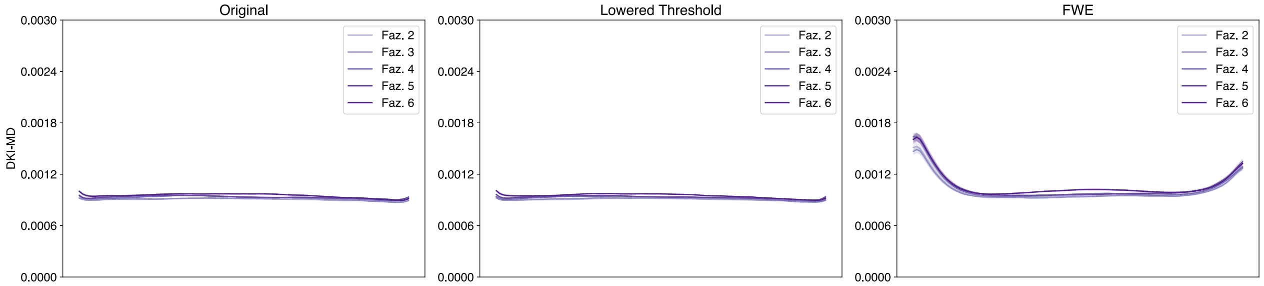


Figure 79: Multi-Shell Left Vertical Occipital DKI-MD profiles.

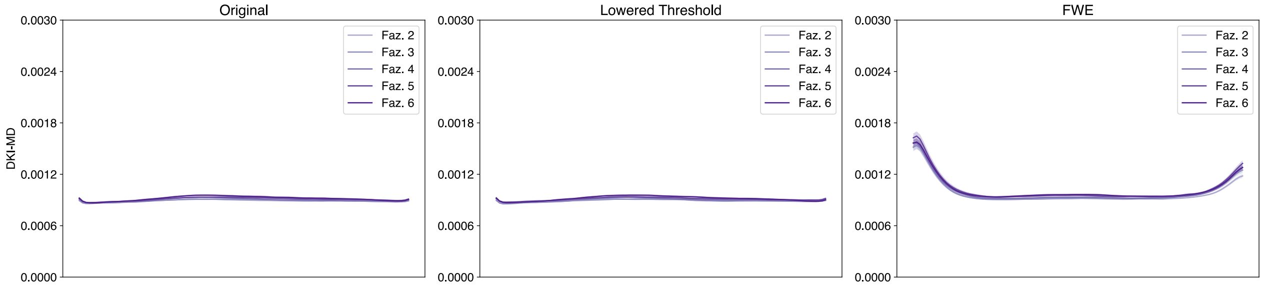


Figure 80: Multi-Shell Right Vertical Occipital DKI-MD profiles.

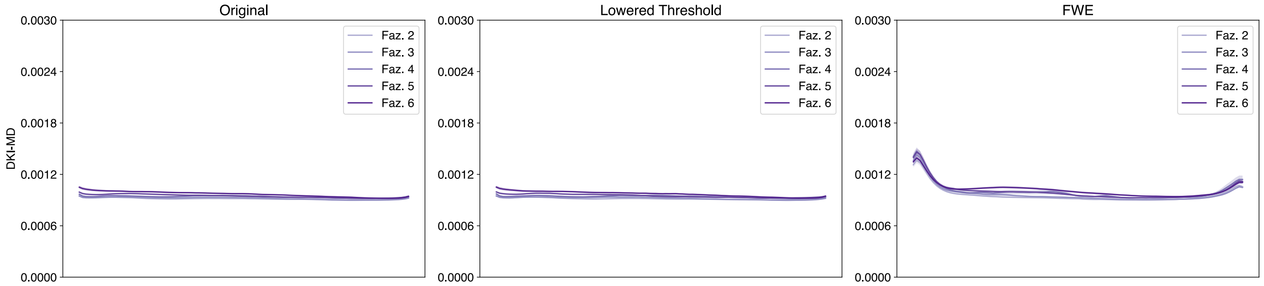


Figure 81: Multi-Shell Left Posterior Arcuate DKI-MD profiles.

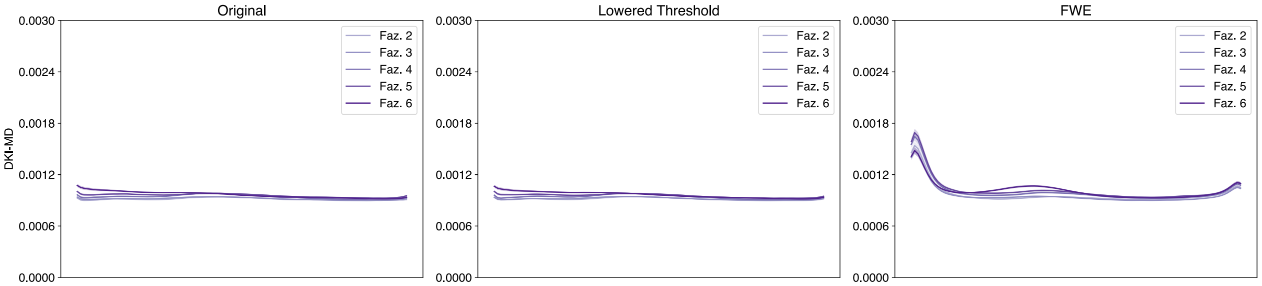


Figure 82: Multi-Shell Right Posterior Arcuate DKI-MD profiles.

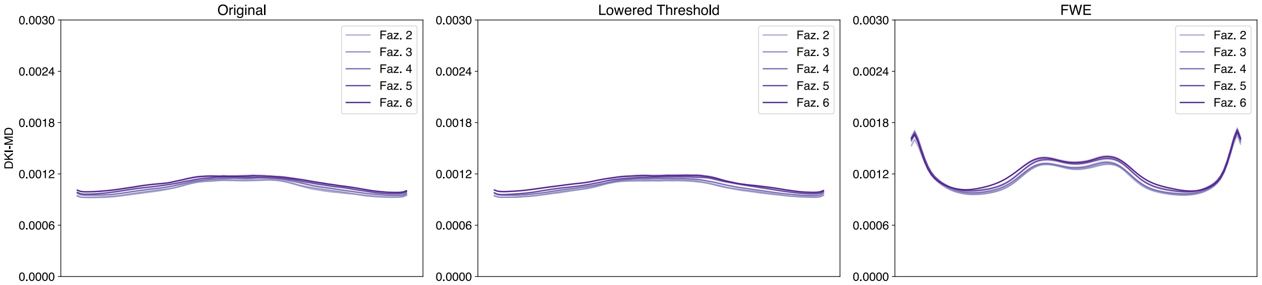


Figure 83: Multi-Shell Anterior Frontal Callosum DKI-MD profiles.

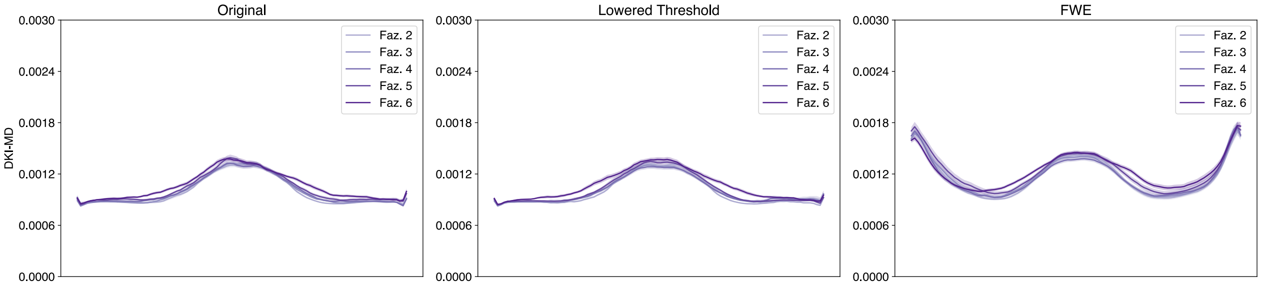


Figure 84: Multi-Shell Motor Corpus Callosum DKI-MD profiles.

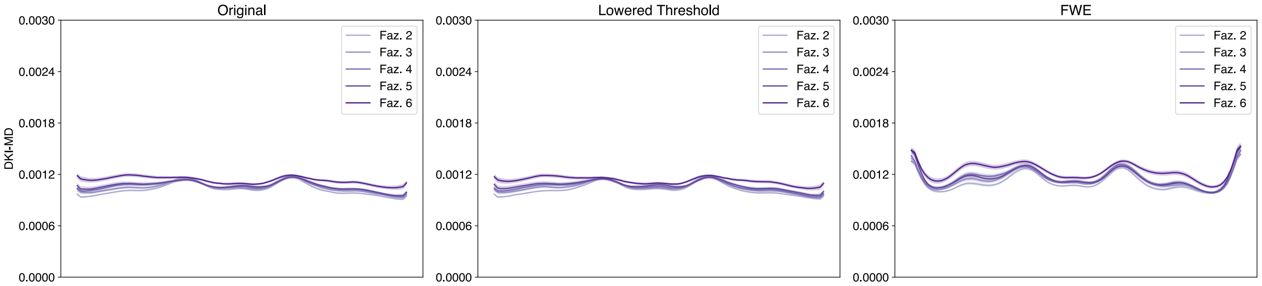


Figure 85: Multi-Shell Occipital Corpus Callosum DKI-MD profiles.

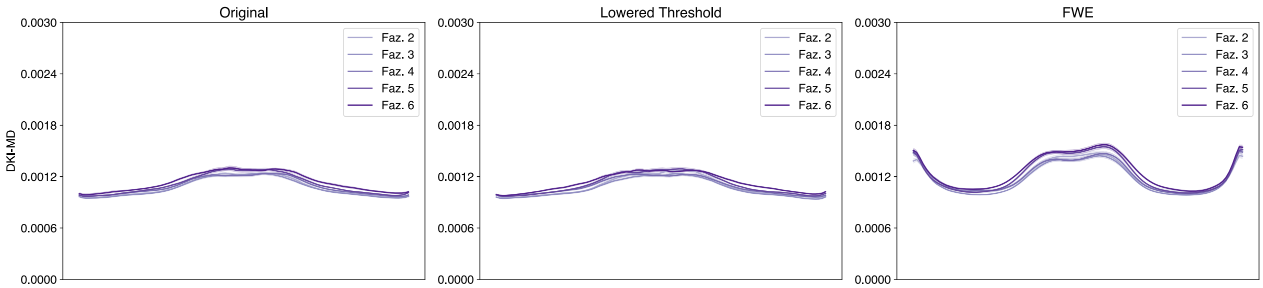


Figure 86: Multi-Shell Orbital Corpus Callosum DKI-MD profiles.

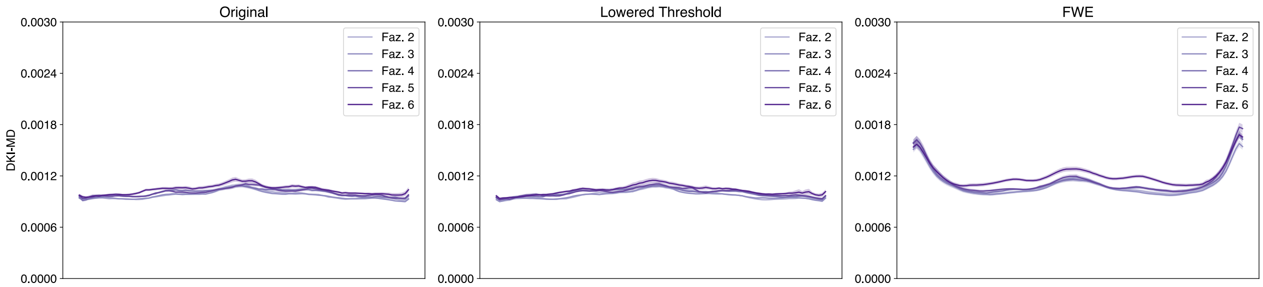


Figure 87: Multi-Shell Posterior Parietal Corpus Callosum DKI-MD profiles.

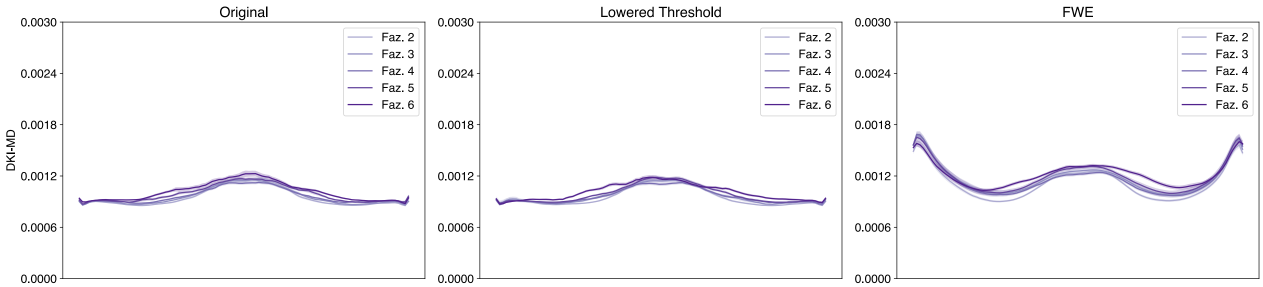


Figure 88: Multi-Shell Superior Frontal Callosum DKI-MD profiles.

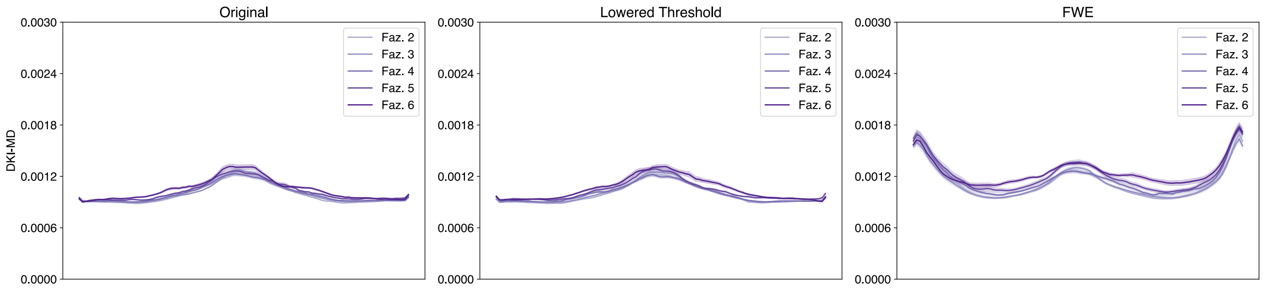


Figure 89: Multi-Shell Superior Parietal Corpus Callosum DKI-MD profiles.

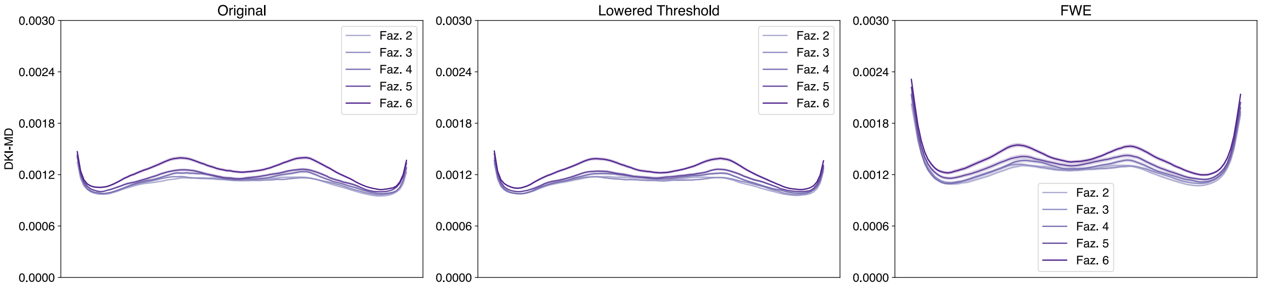


Figure 90: Multi-Shell Temporal Corpus Callosum DKI-MD profiles.

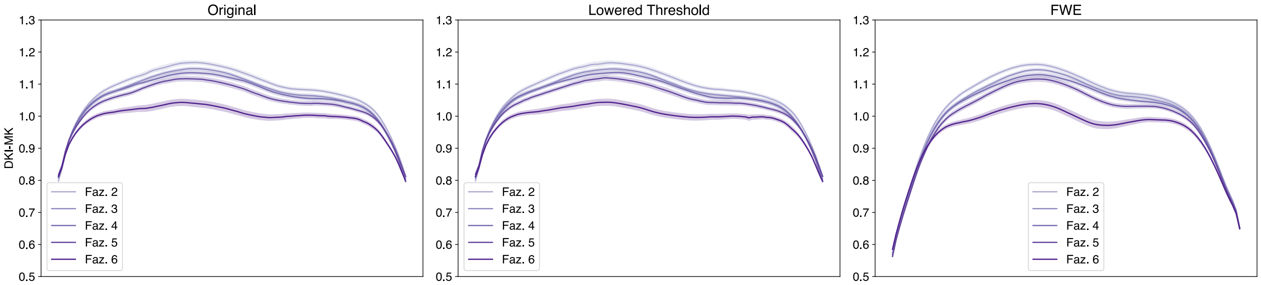


Figure 91: Multi-Shell Left Arcuate DKI-MK profiles.

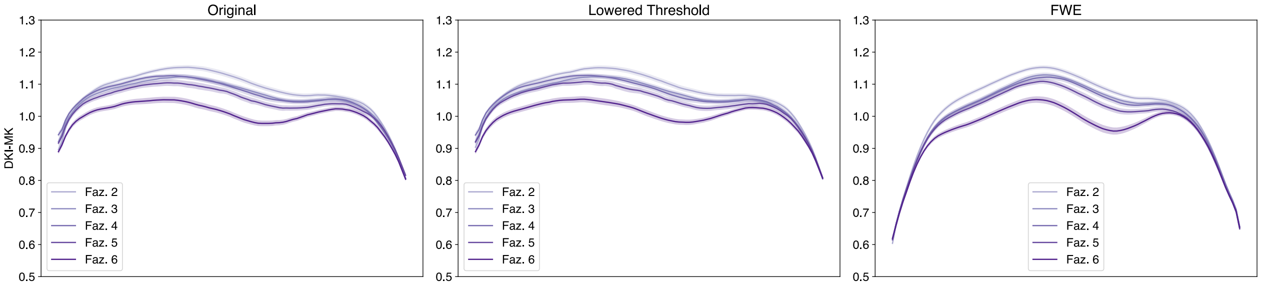


Figure 92: Multi-Shell Right Arcuate DKI-MK profiles.

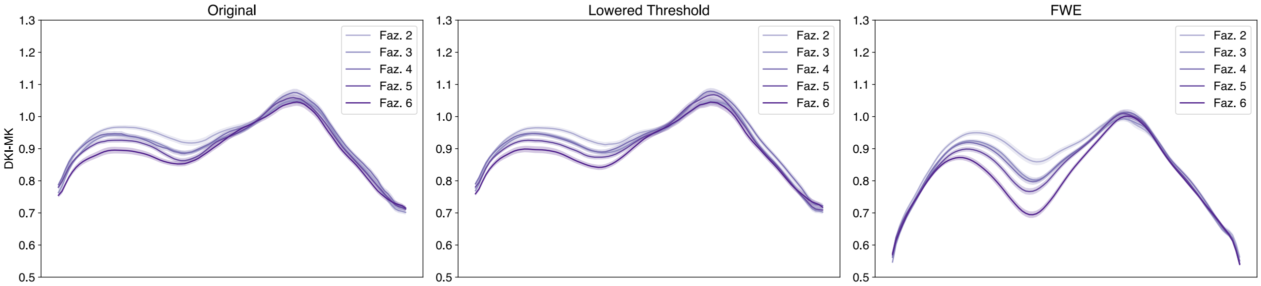


Figure 93: Multi-Shell Left Anterior Thalamic DKI-MK profiles.

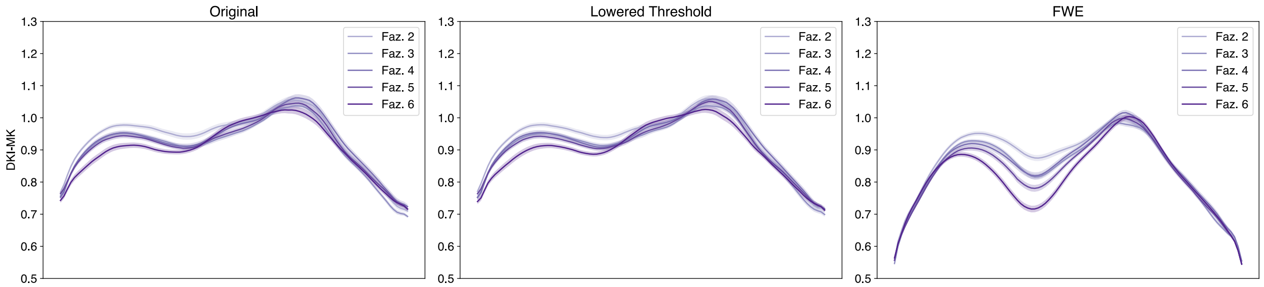


Figure 94: Multi-Shell Right Anterior Thalamic DKI-MK profiles.

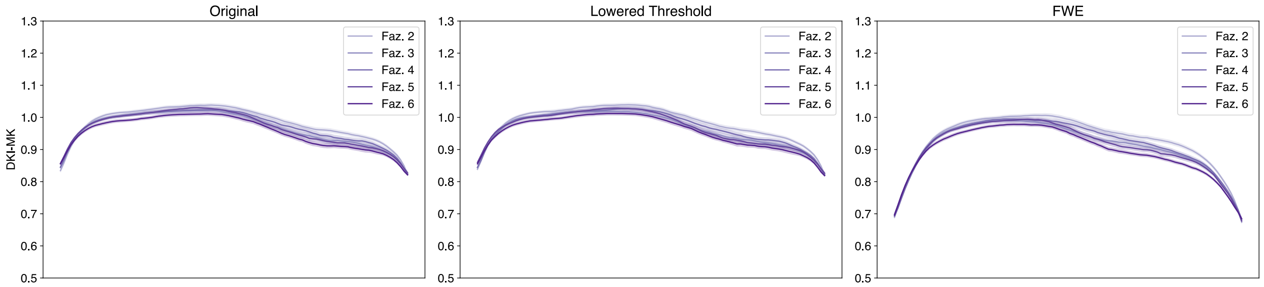


Figure 95: Multi-Shell Left Cingulum Cingulate DKI-MK profiles.

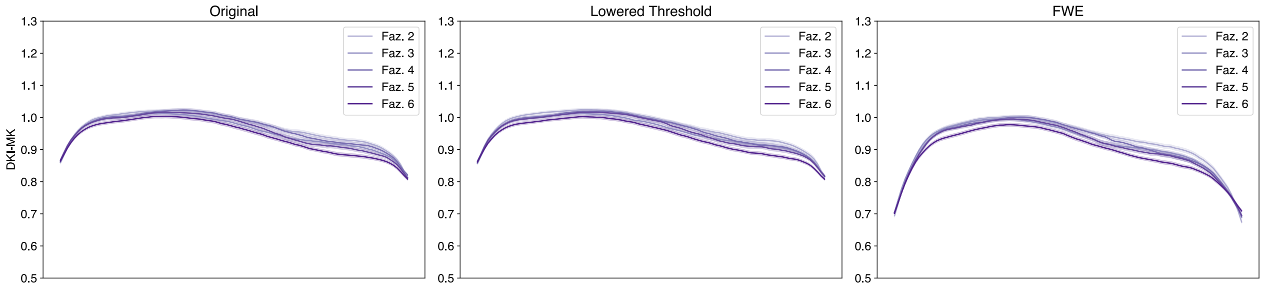


Figure 96: Multi-Shell Right Cingulum Cingulate DKI-MK profiles.

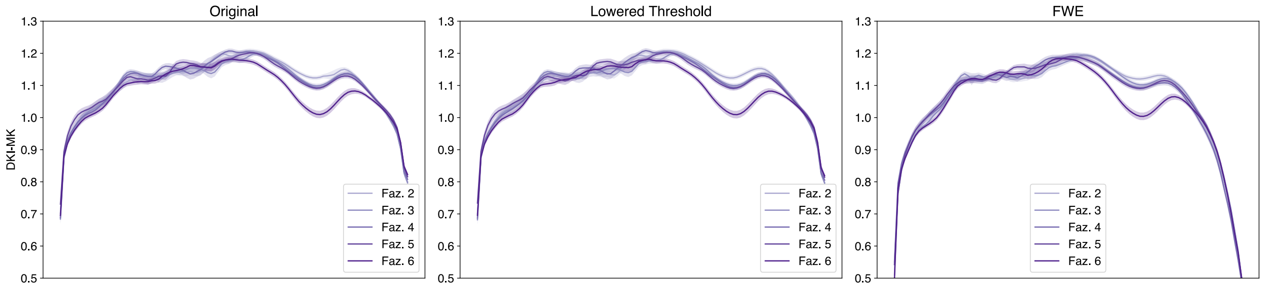


Figure 97: Multi-Shell Left Corticospinal DKI-MK profiles.

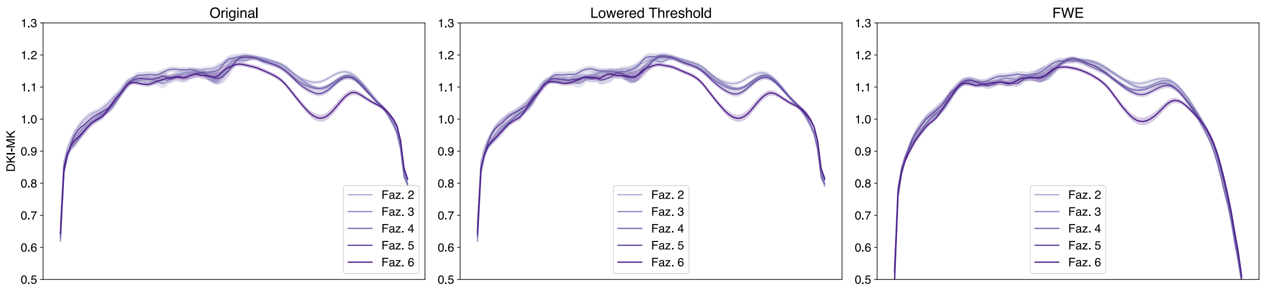


Figure 98: Multi-Shell Right Corticospinal DKI-MK profiles.

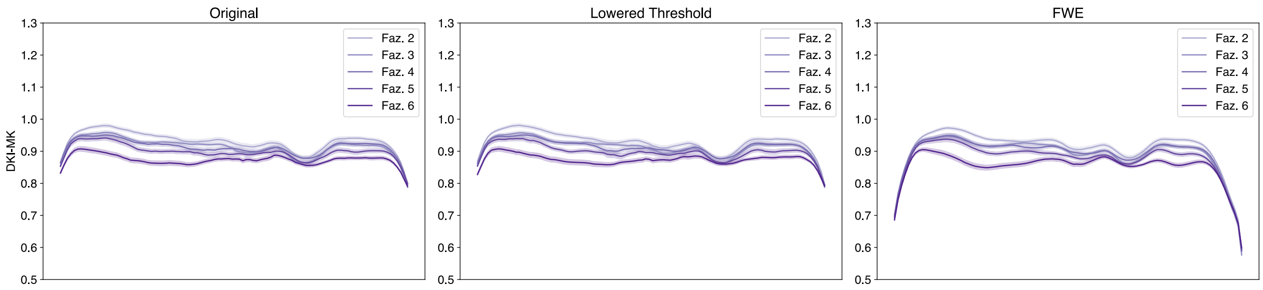


Figure 99: Multi-Shell Left Inferior Fronto-Occipital DKI-MK profiles.

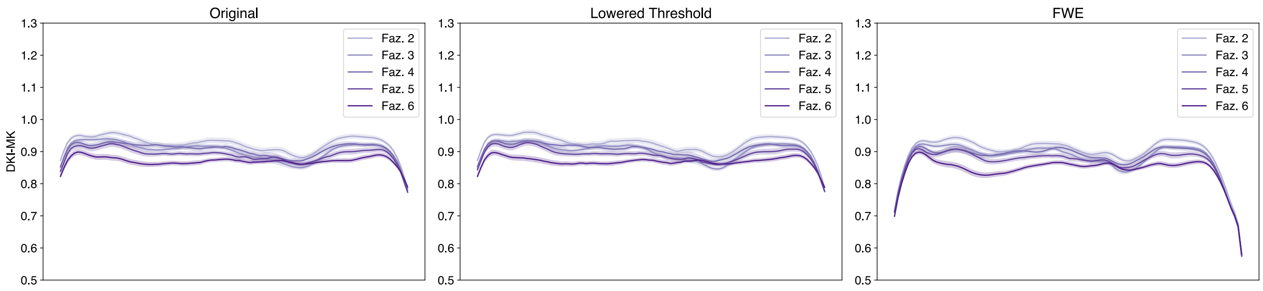


Figure 100: Multi-Shell Right Inferior Fronto-Occipital DKI-MK profiles.

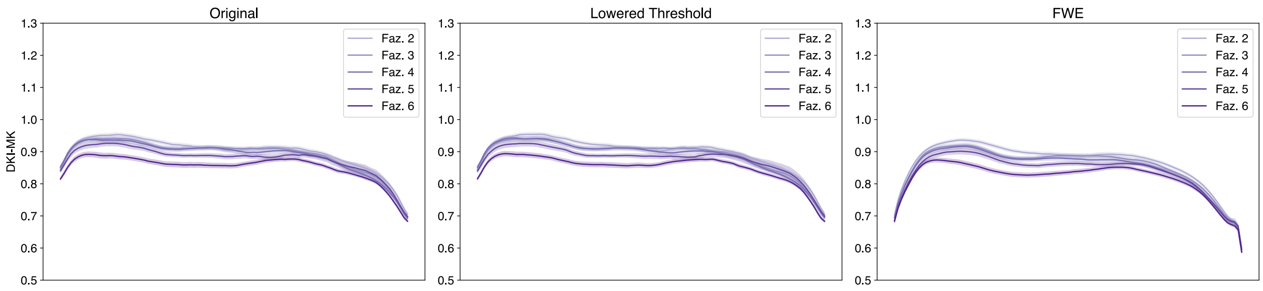


Figure 101: Multi-Shell Left Inferior Longitudinal DKI-MK profiles.

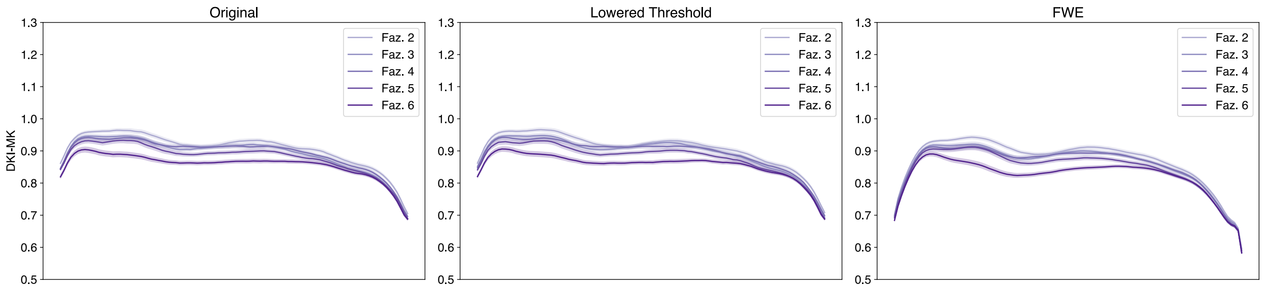


Figure 102: Multi-Shell Right Inferior Longitudinal DKI-MK profiles.

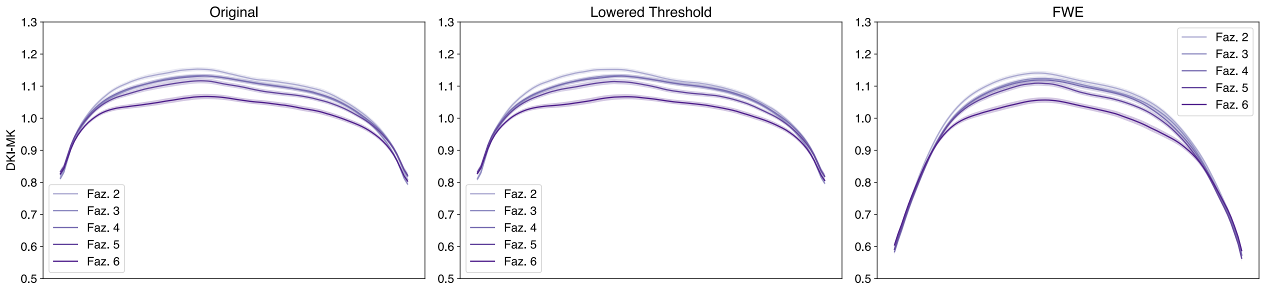


Figure 103: Multi-Shell Left Superior Longitudinal DKI-MK profiles.

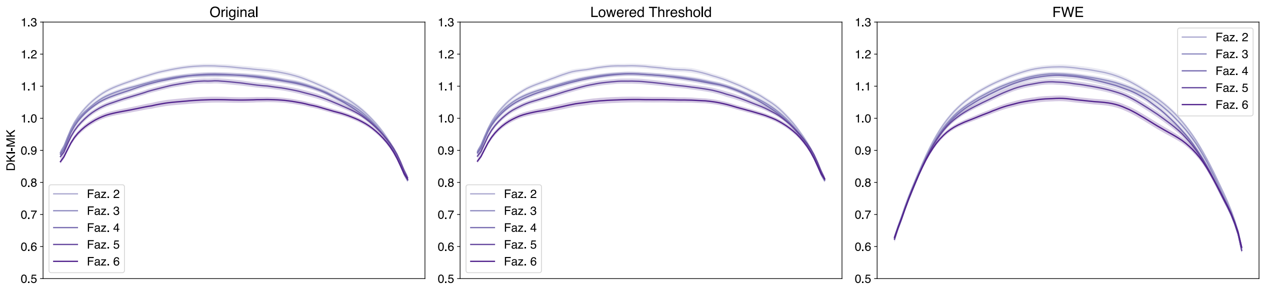


Figure 104: Multi-Shell Right Superior Longitudinal DKI-MK profiles.

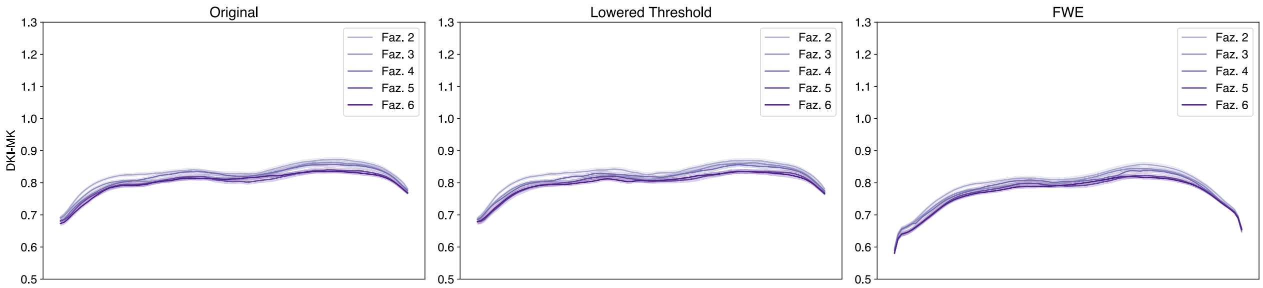


Figure 105: Multi-Shell Left Uncinate DKI-MK profiles.

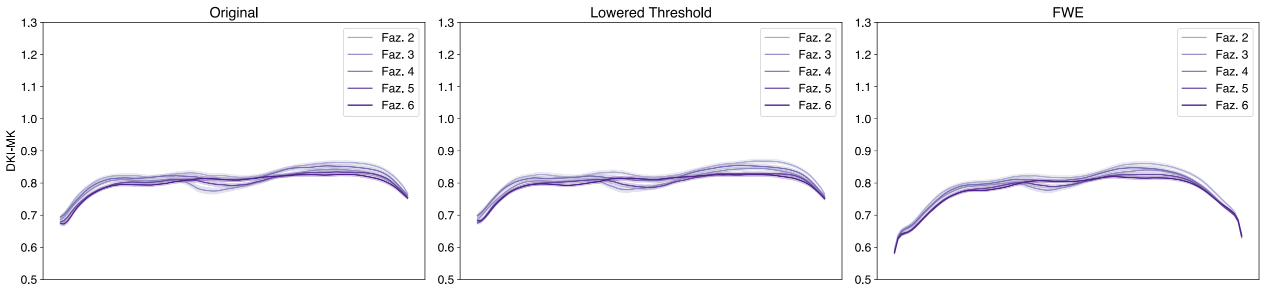


Figure 106: Multi-Shell Right Uncinate DKI-MK profiles.

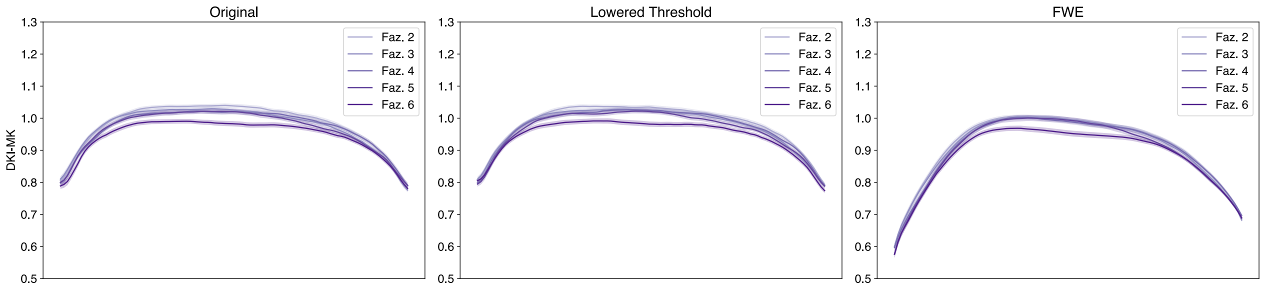


Figure 107: Multi-Shell Left Vertical Occipital DKI-MK profiles.

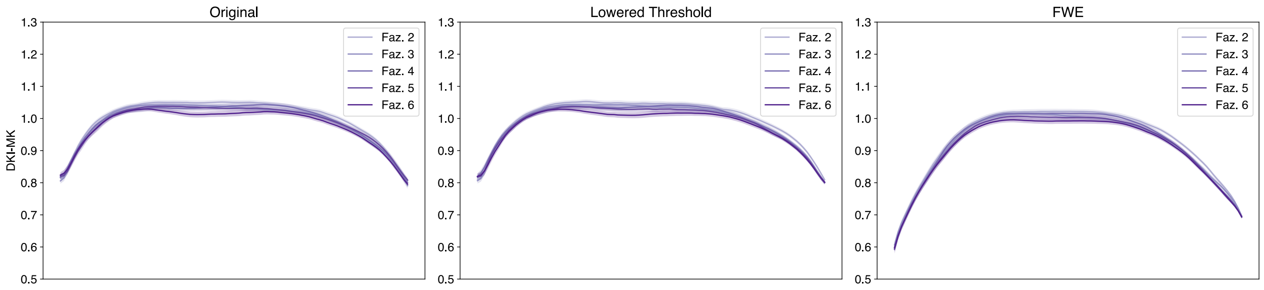


Figure 108: Multi-Shell Right Vertical Occipital DKI-MK profiles.

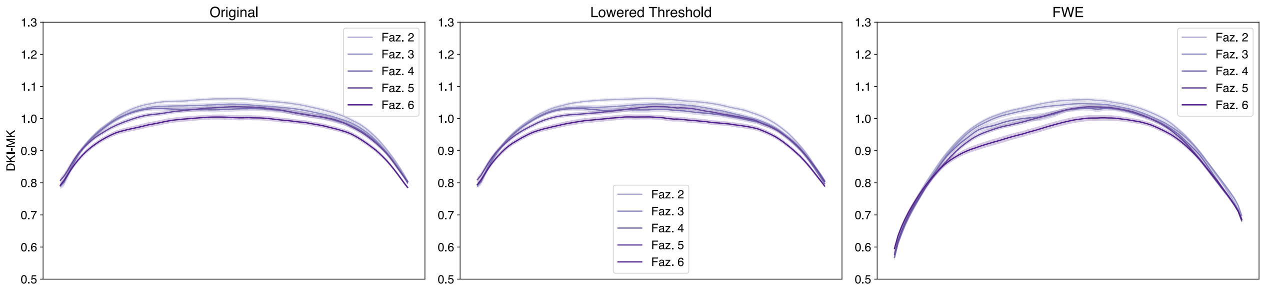


Figure 109: Multi-Shell Left Posterior Arcuate DKI-MK profiles.

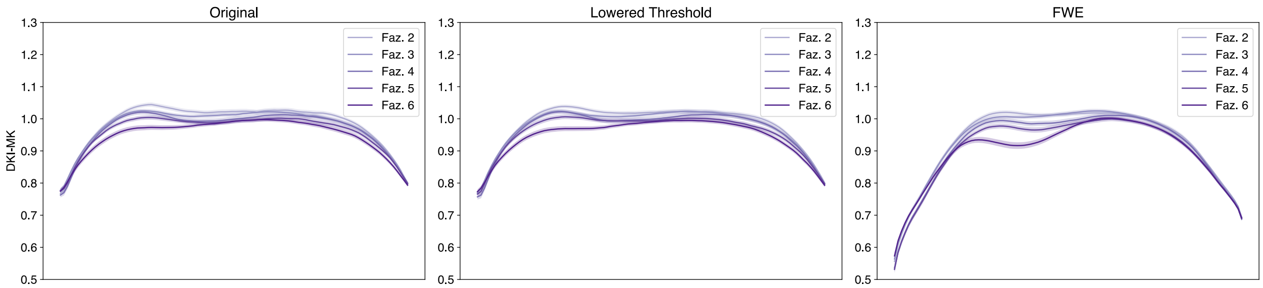


Figure 110: Multi-Shell Right Posterior Arcuate DKI-MK profiles.

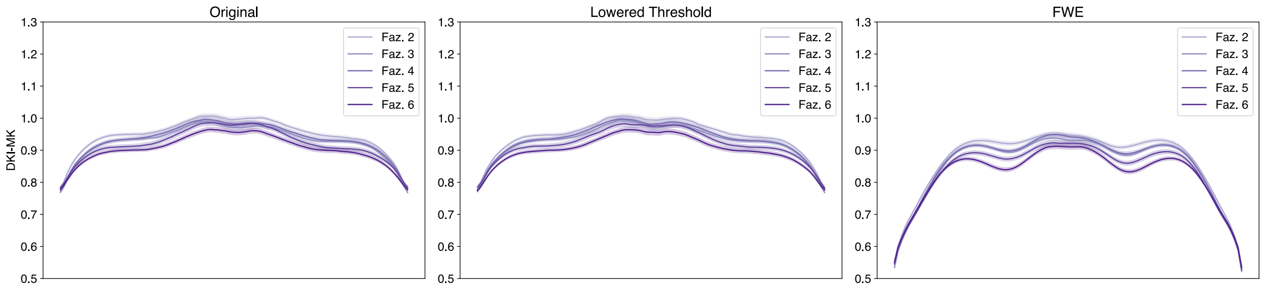


Figure 111: Multi-Shell Anterior Frontal Callosum DKI-MK profiles.

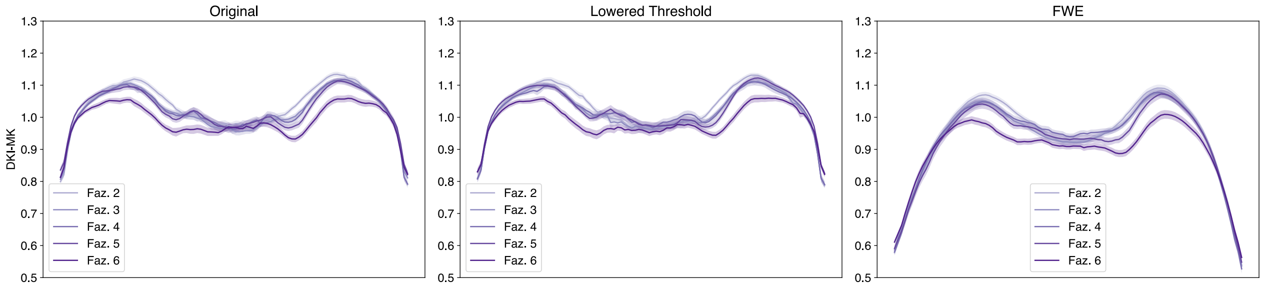


Figure 112: Multi-Shell Motor Corpus Callosum DKI-MK profiles.

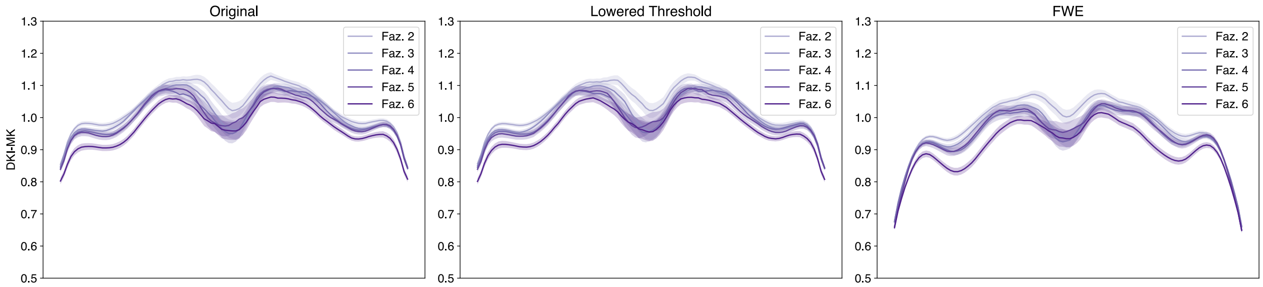


Figure 113: Multi-Shell Occipital Corpus Callosum DKI-MK profiles.

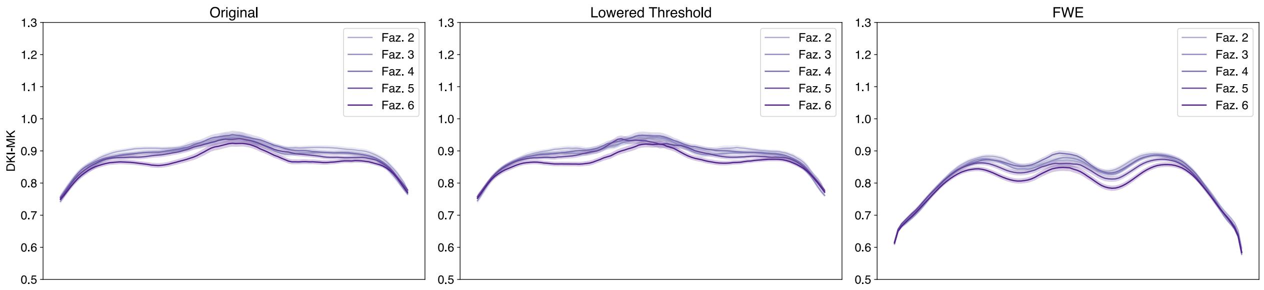


Figure 114: Multi-Shell Orbital Corpus Callosum DKI-MK profiles.

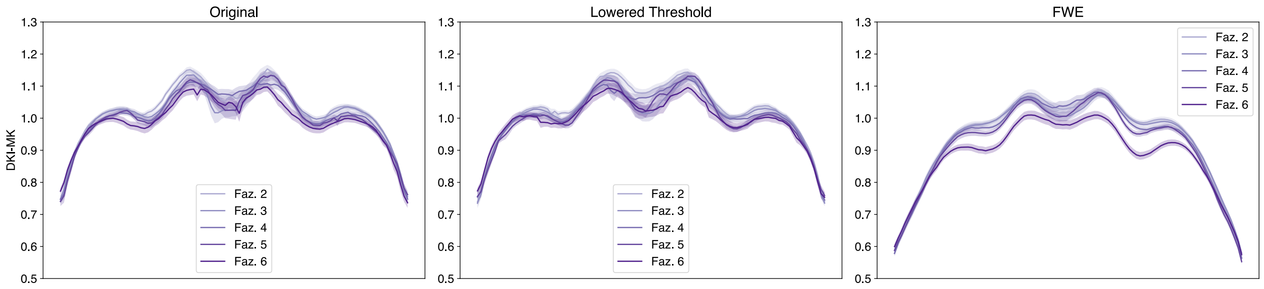


Figure 115: Multi-Shell Posterior Parietal Corpus Callosum DKI-MK profiles.

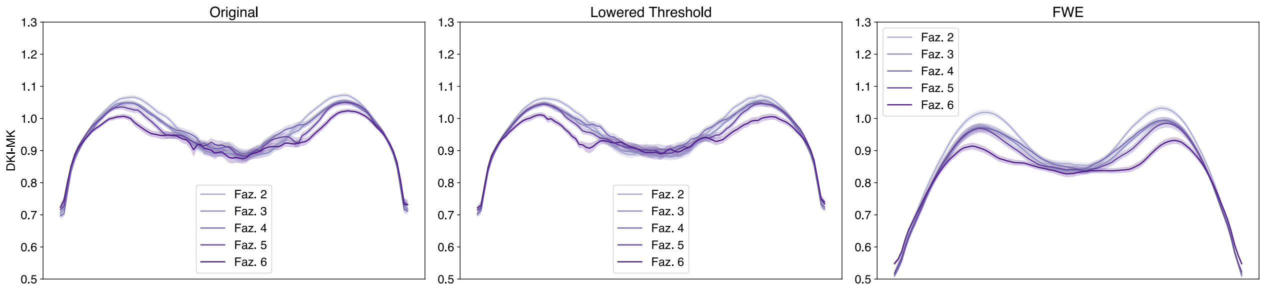


Figure 116: Multi-Shell Superior Frontal Callosum DKI-MK profiles.

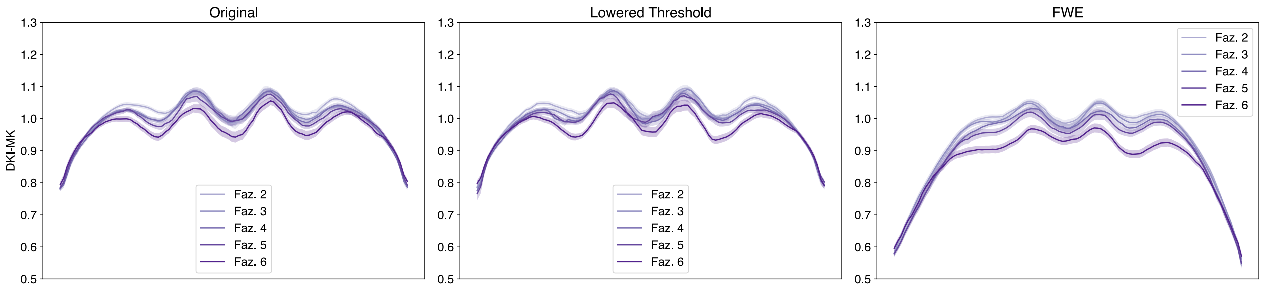


Figure 117: Multi-Shell Superior Parietal Corpus Callosum DKI-MK profiles.

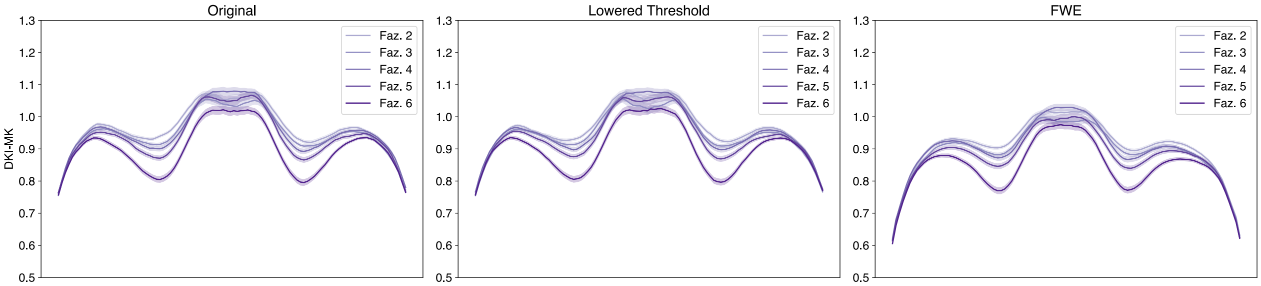


Figure 118: Multi-Shell Temporal Corpus Callosum DKI-MK profiles.

##### 1.3.2 Single-shell dataset

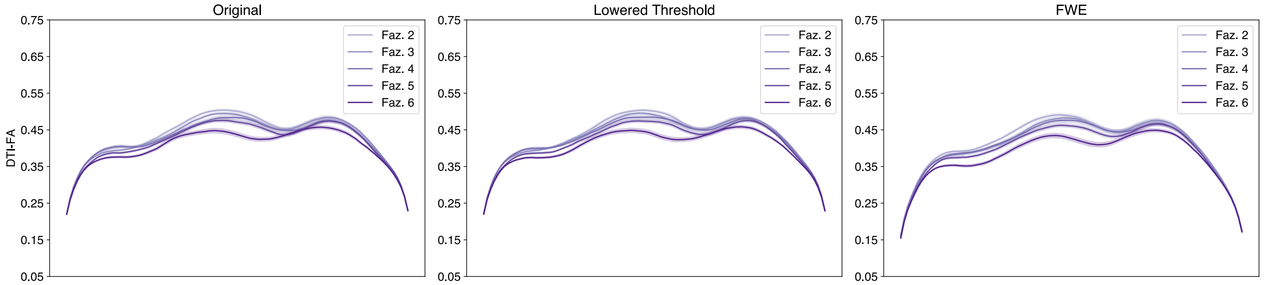


Figure 119: Single-Shell Left Arcuate DTI-FA profiles.

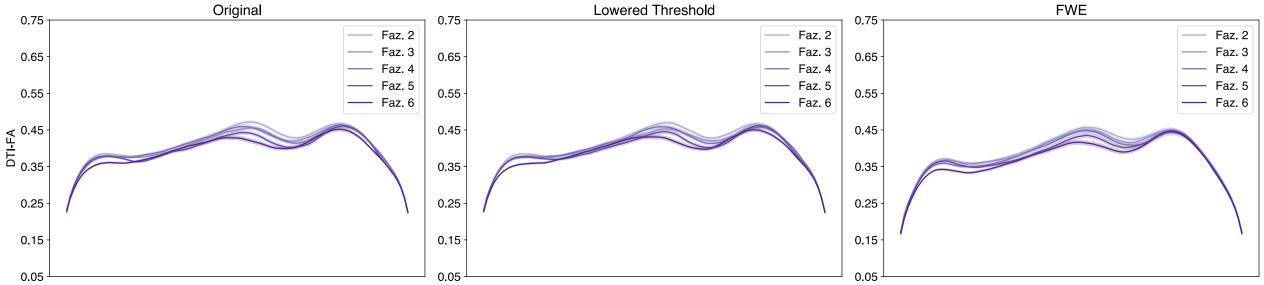


Figure 120: Single-Shell Right Arcuate DTI-FA profiles.

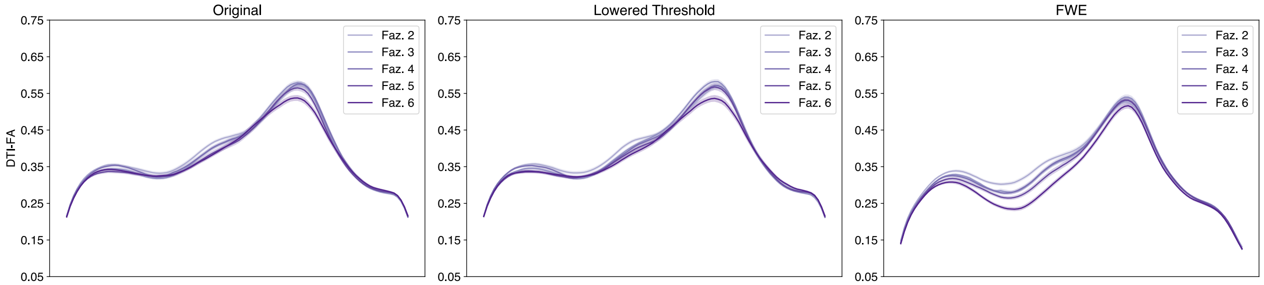


Figure 121: Single-Shell Left Anterior Thalamic DTI-FA profiles.

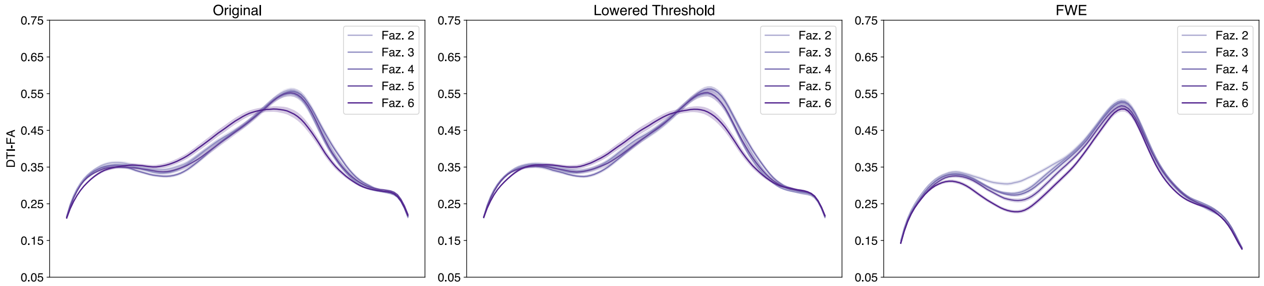


Figure 122: Single-Shell Right Anterior Thalamic DTI-FA profiles.

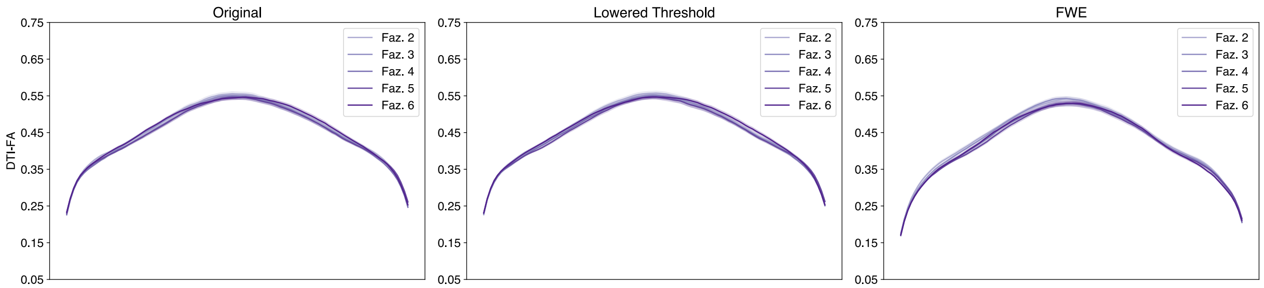


Figure 123: Single-Shell Left Cingulum Cingulate DTI-FA profiles.

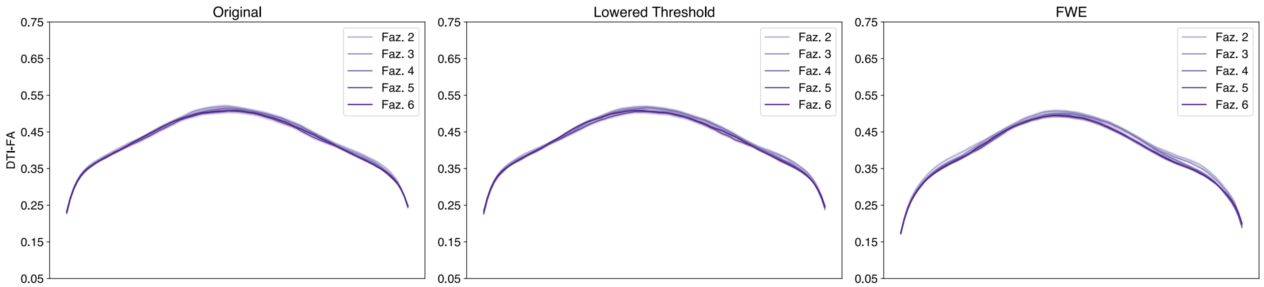


Figure 124: Single-Shell Right Cingulum Cingulate DTI-FA profiles.

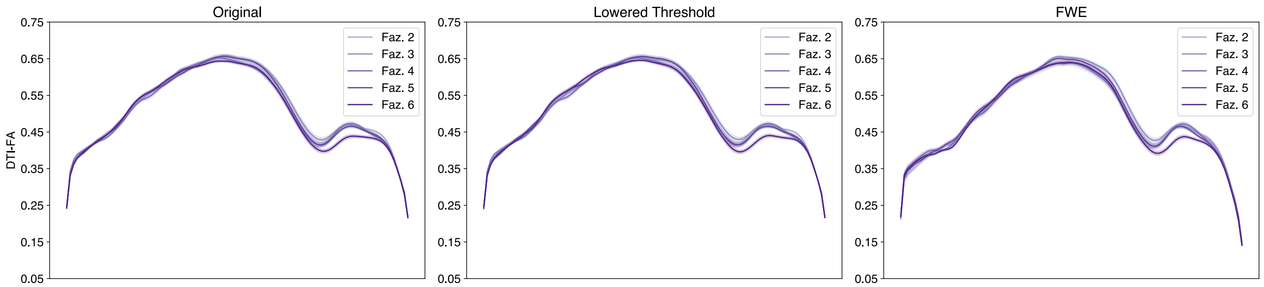


Figure 125: Single-Shell Left Corticospinal DTI-FA profiles.

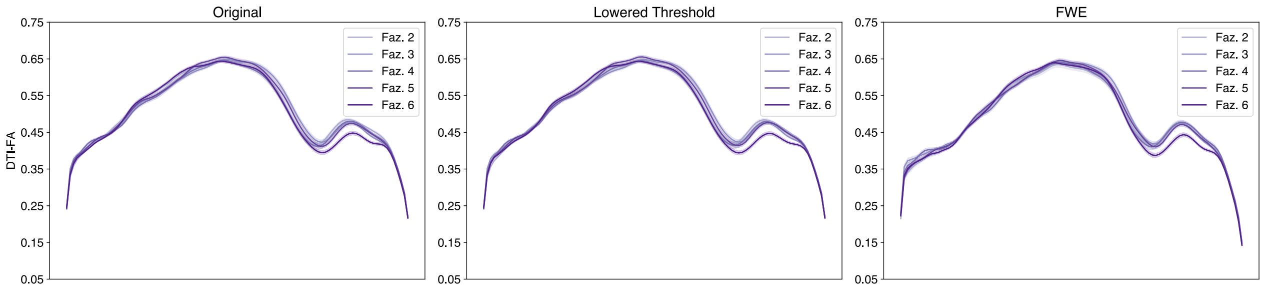


Figure 126: Single-Shell Right Corticospinal DTI-FA profiles.

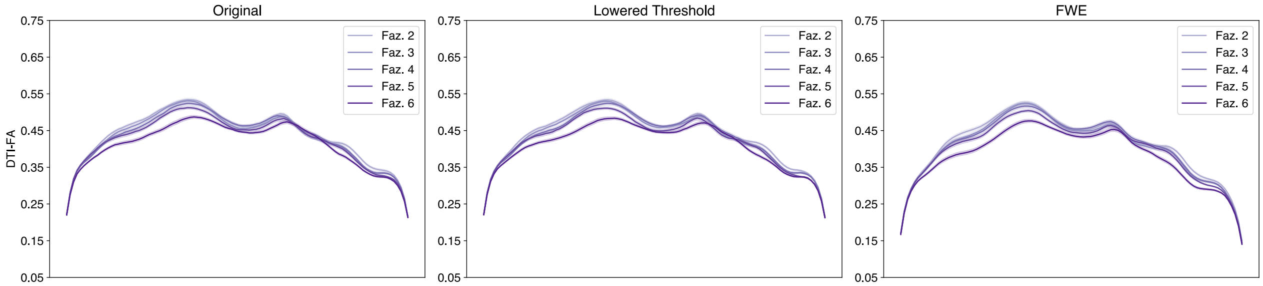


Figure 127: Single-Shell Left Inferior Fronto-Occipital DTI-FA profiles.

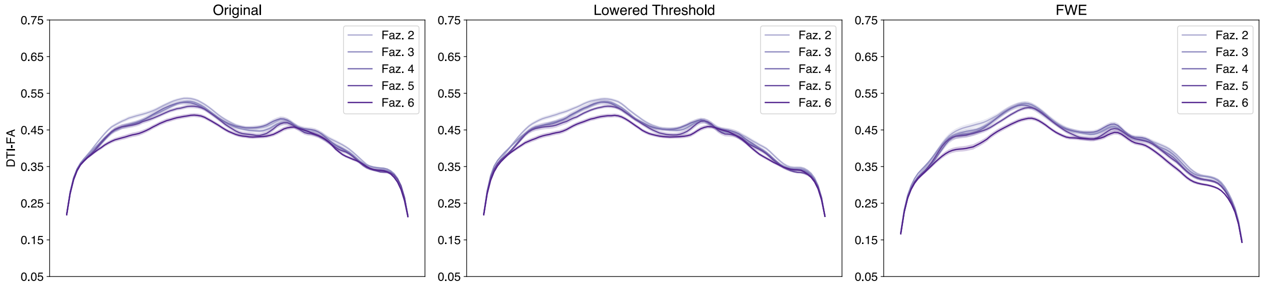


Figure 128: Single-Shell Right Inferior Fronto-Occipital DTI-FA profiles.

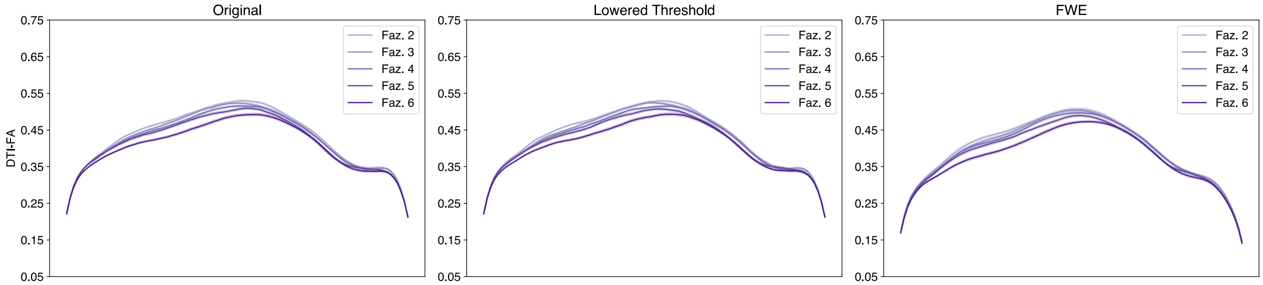


Figure 129: Single-Shell Left Inferior Longitudinal DTI-FA profiles.

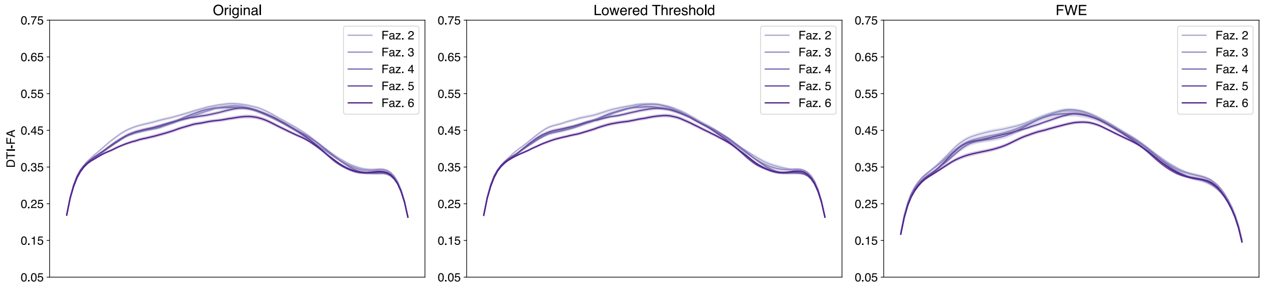


Figure 130: Single-Shell Right Inferior Longitudinal DTI-FA profiles.

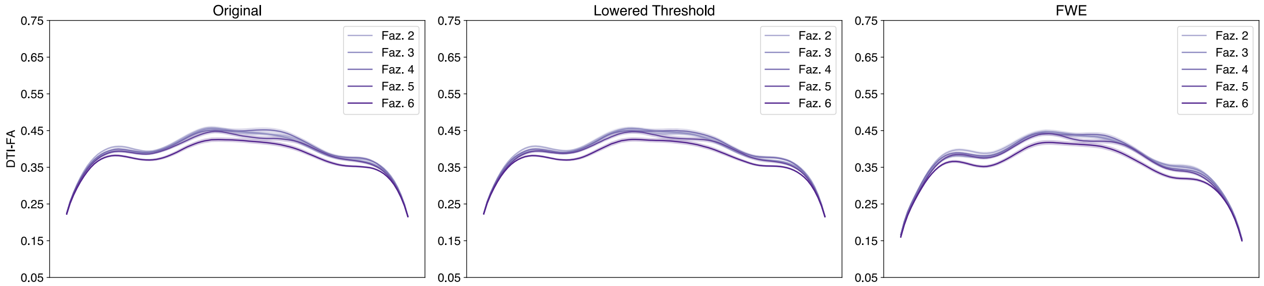


Figure 131: Single-Shell Left Superior Longitudinal DTI-FA profiles.

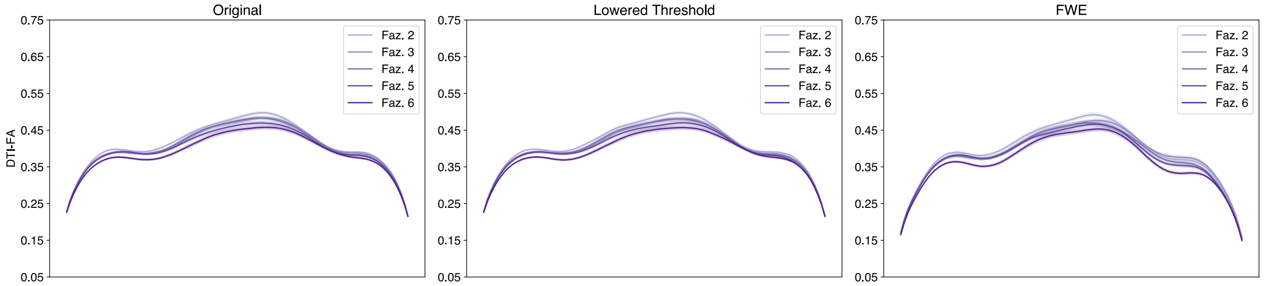


Figure 132: Single-Shell Right Superior Longitudinal DTI-FA profiles.

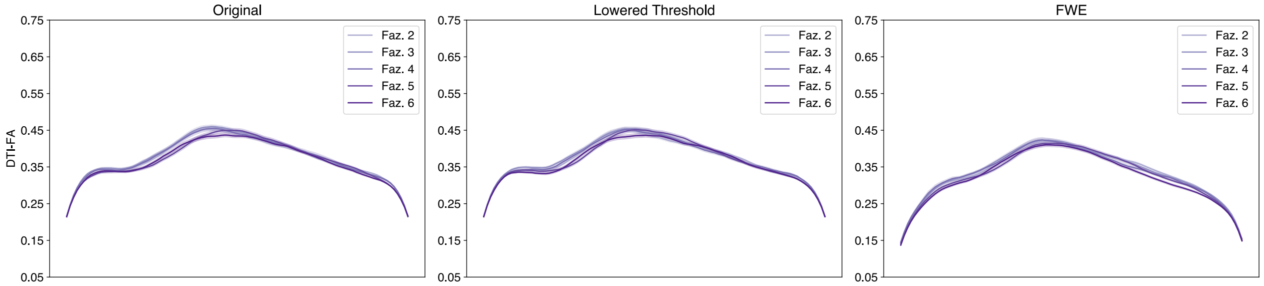


Figure 133: Single-Shell Left Uncinate DTI-FA profiles.

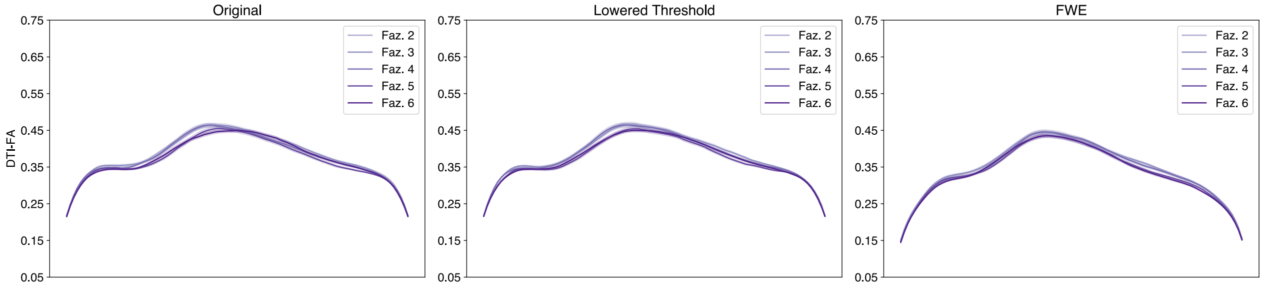


Figure 134: Single-Shell Right Uncinate DTI-FA profiles.

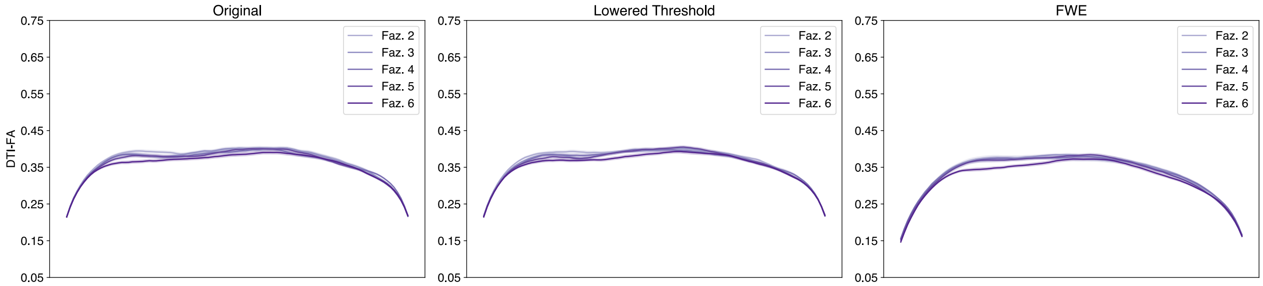


Figure 135: Single-Shell Left Vertical Occipital DTI-FA profiles.

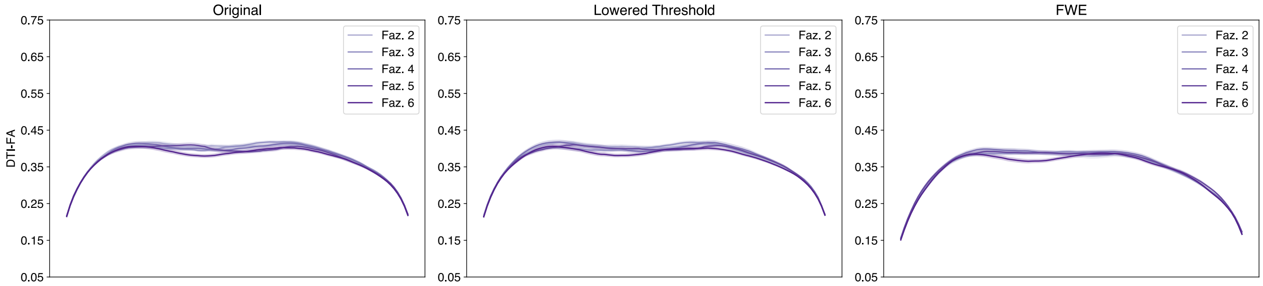


Figure 136: Single-Shell Right Vertical Occipital DTI-FA profiles.

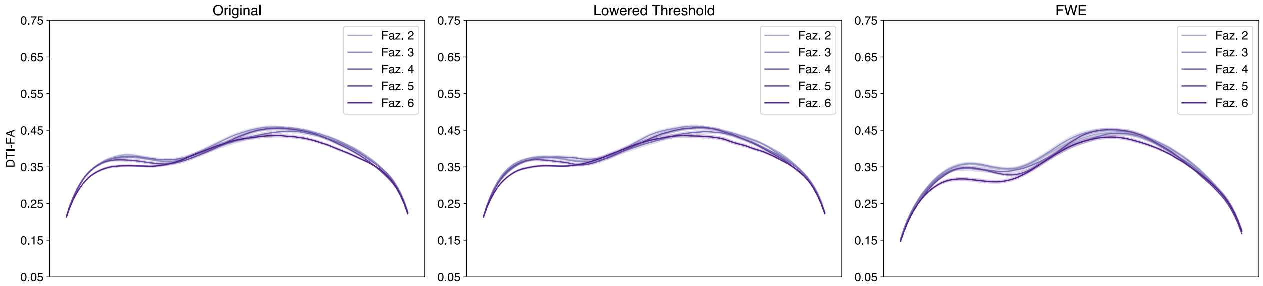


Figure 137: Single-Shell Left Posterior Arcuate DTI-FA profiles.

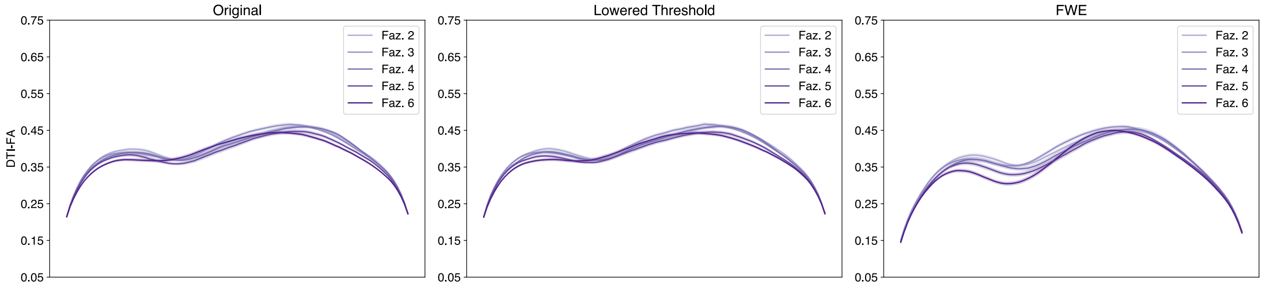


Figure 138: Single-Shell Right Posterior Arcuate DTI-FA profiles.

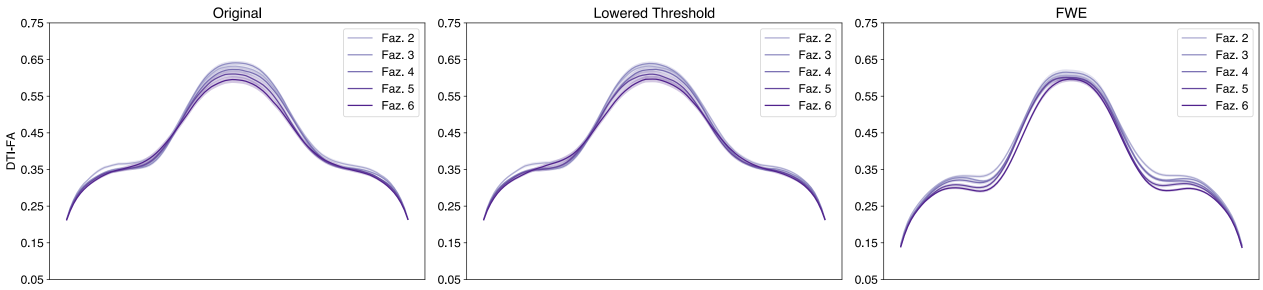


Figure 139: Single-Shell Anterior Frontal Callosum DTI-FA profiles.

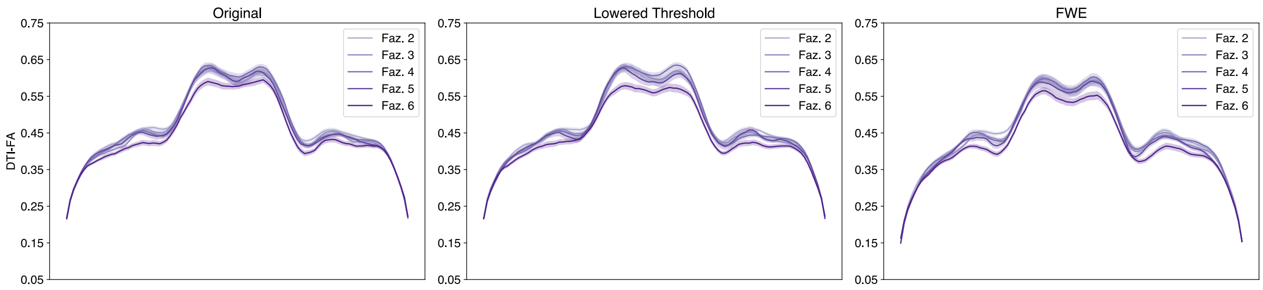


Figure 140: Single-Shell Motor Corpus Callosum DTI-FA profiles.

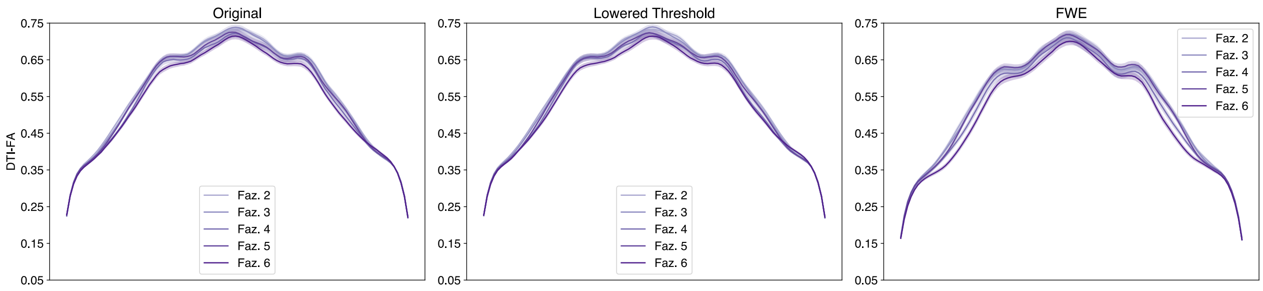


Figure 141: Single-Shell Occipital Corpus Callosum DTI-FA profiles.

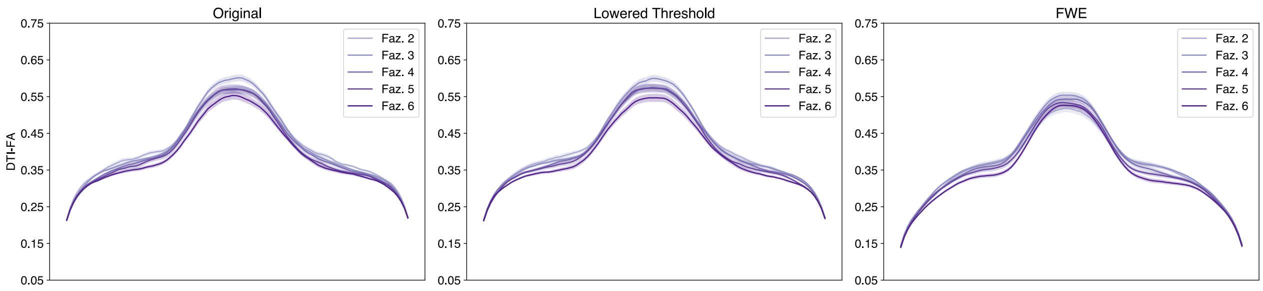


Figure 142: Single-Shell Orbital Corpus Callosum DTI-FA profiles.

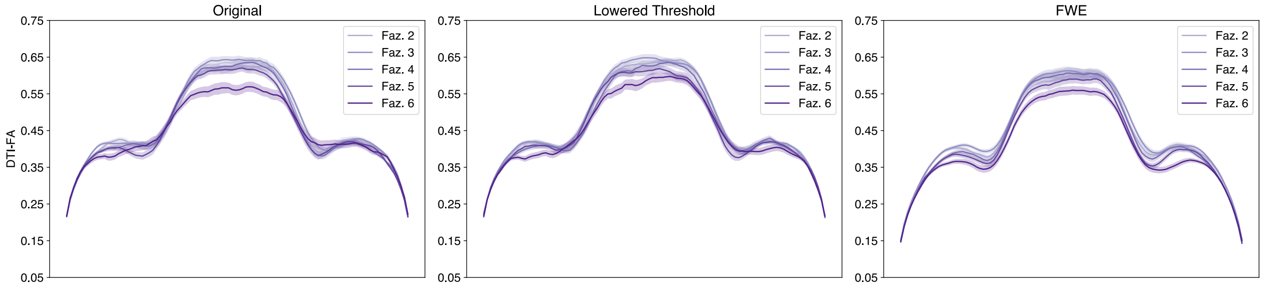


Figure 143: Single-Shell Posterior Parietal Corpus Callosum DTI-FA profiles.

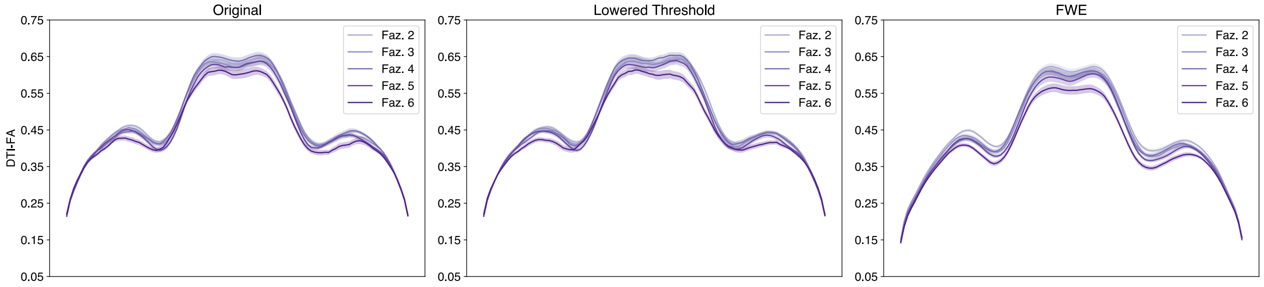


Figure 144: Single-Shell Superior Frontal Callosum DTI-FA profiles.

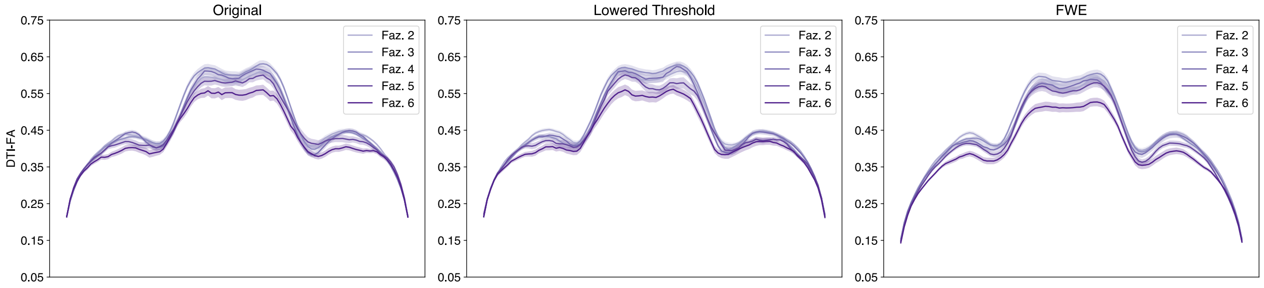


Figure 145: Single-Shell Superior Parietal Corpus Callosum DTI-FA profiles.

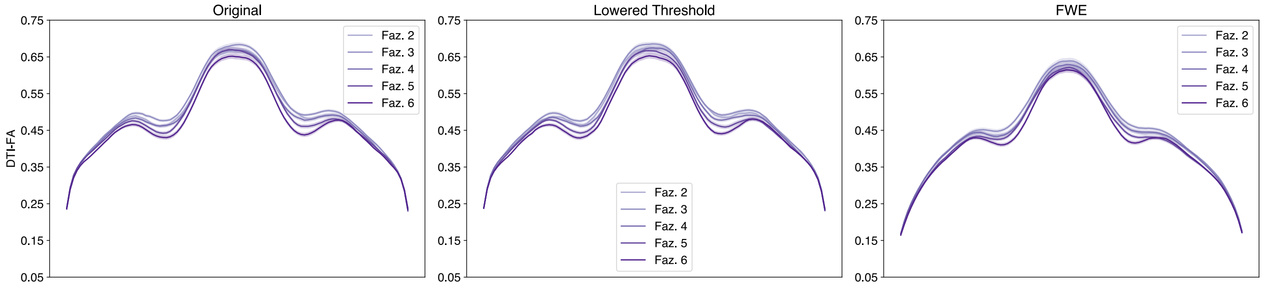


Figure 146: Single-Shell Temporal Corpus Callosum DTI-FA profiles.

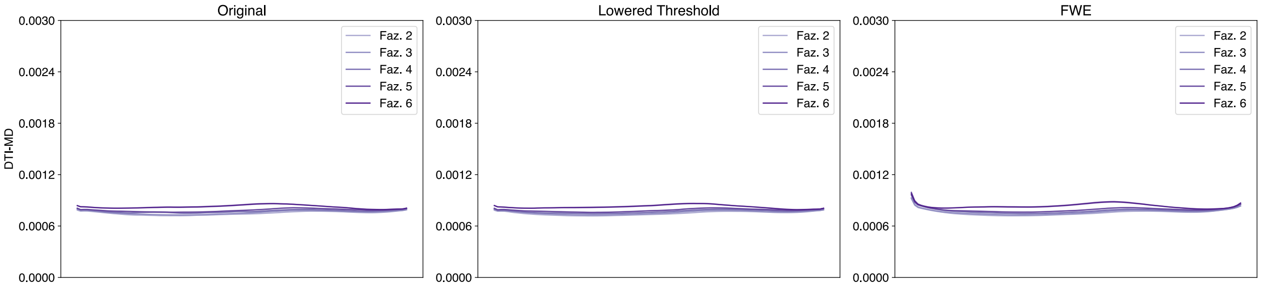


Figure 147: Single-Shell Left Arcuate DTI-MD profiles.

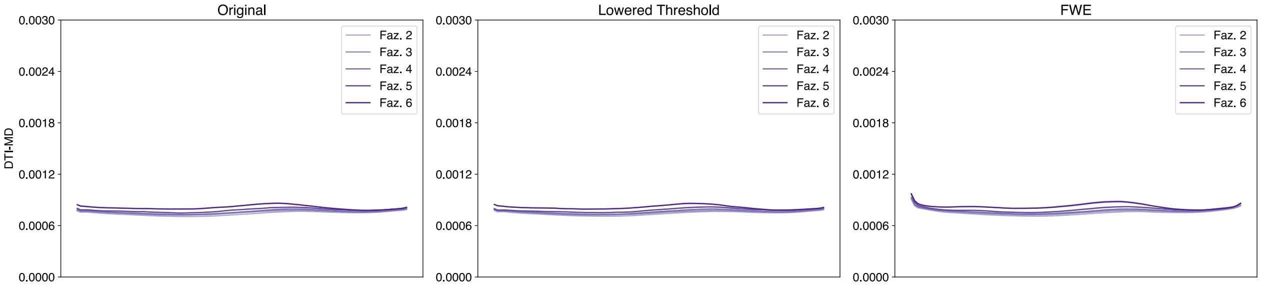


Figure 148: Single-Shell Right Arcuate DTI-MD profiles.

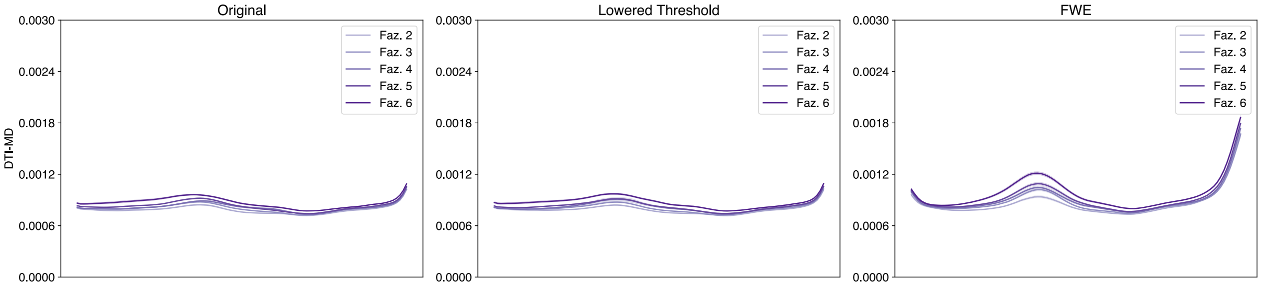


Figure 149: Single-Shell Left Anterior Thalamic DTI-MD profiles.

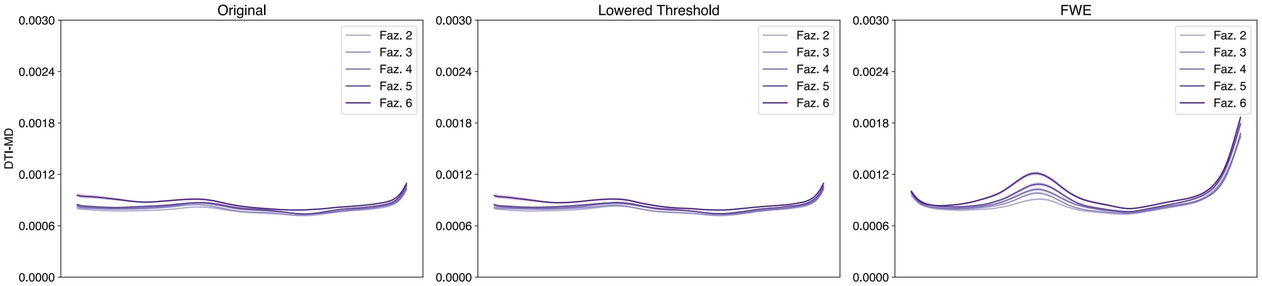


Figure 150: Single-Shell Right Anterior Thalamic DTI-MD profiles.

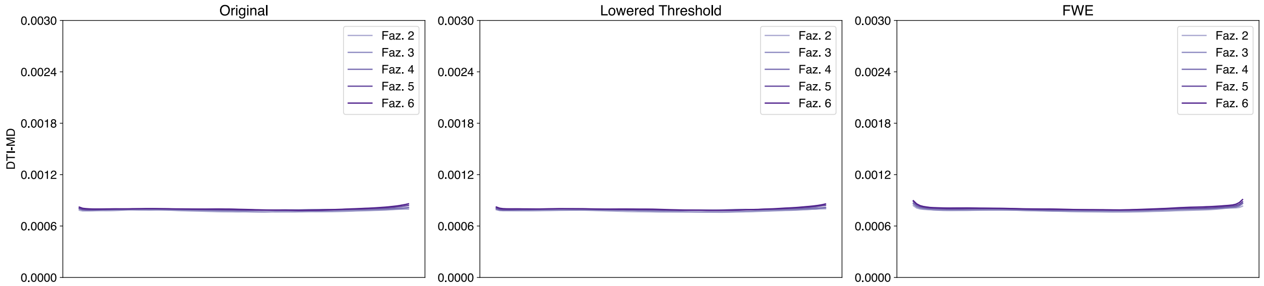


Figure 151: Single-Shell Left Cingulum Cingulate DTI-MD profiles.

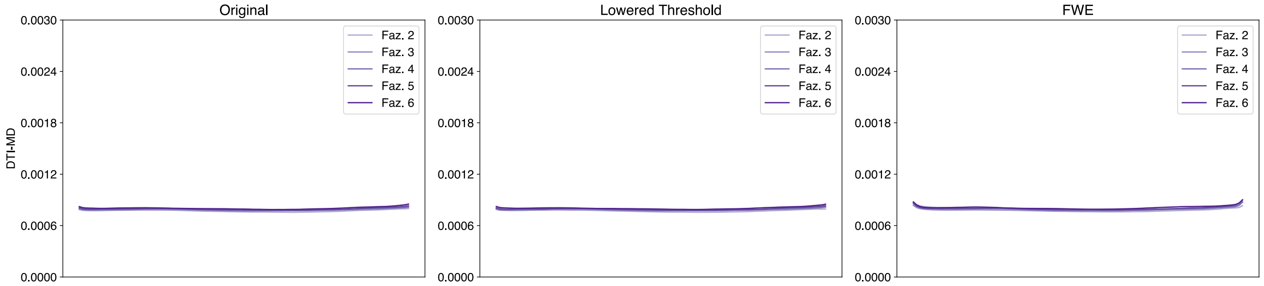


Figure 152: Single-Shell Right Cingulum Cingulate DTI-MD profiles.

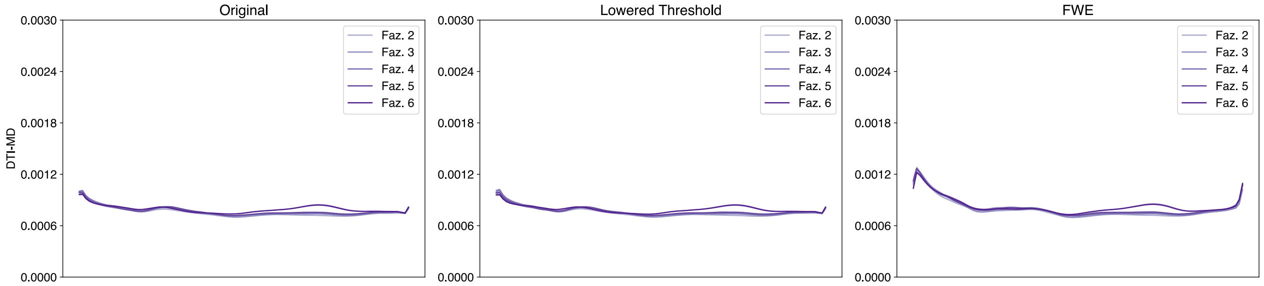


Figure 153: Single-Shell Left Corticospinal DTI-MD profiles.

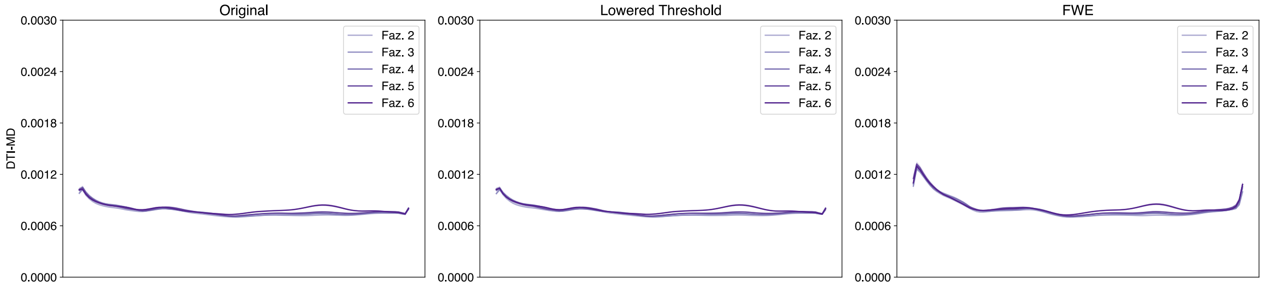


Figure 154: Single-Shell Right Corticospinal DTI-MD profiles.

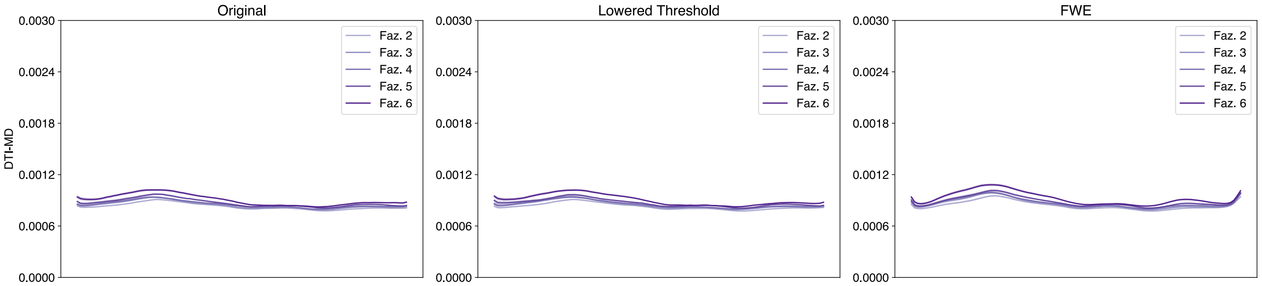


Figure 155: Single-Shell Left Inferior Fronto-Occipital DTI-MD profiles.

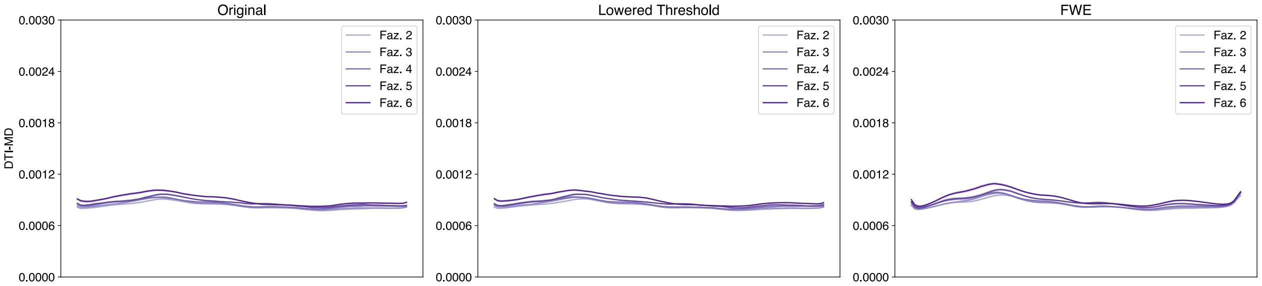


Figure 156: Single-Shell Right Inferior Fronto-Occipital DTI-MD profiles.

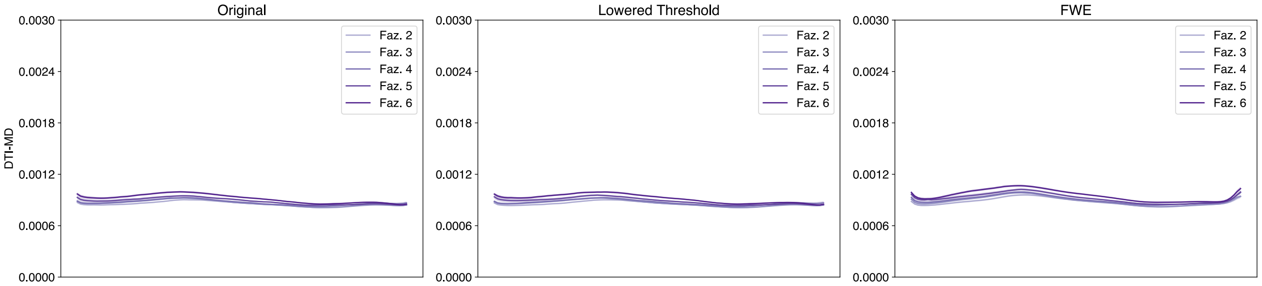


Figure 157: Single-Shell Left Inferior Longitudinal DTI-MD profiles.

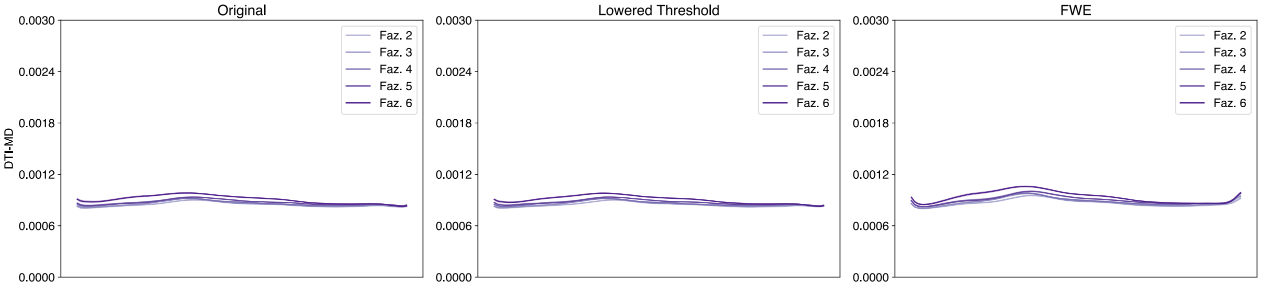


Figure 158: Single-Shell Right Inferior Longitudinal DTI-MD profiles.

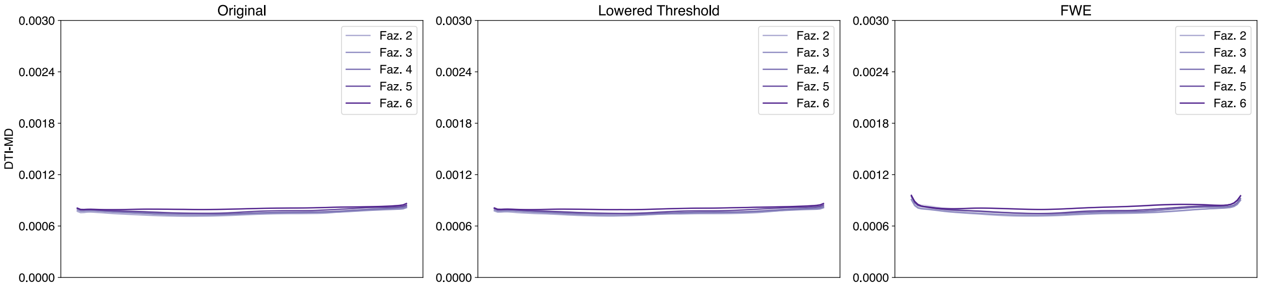


Figure 159: Single-Shell Left Superior Longitudinal DTI-MD profiles.

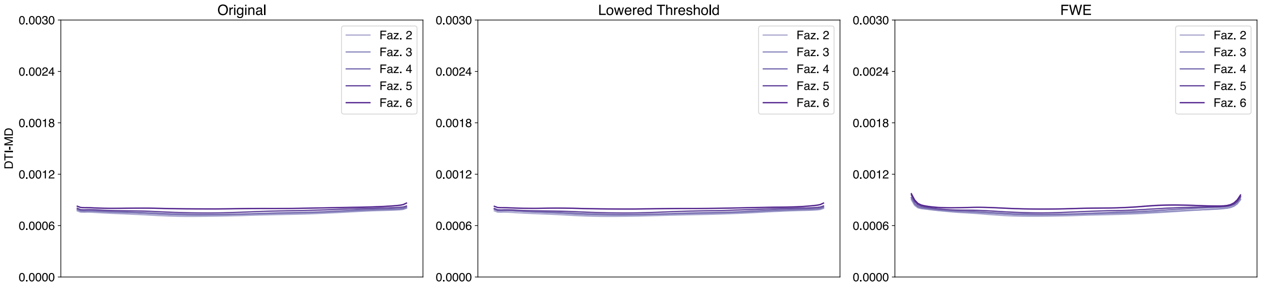


Figure 160: Single-Shell Right Superior Longitudinal DTI-MD profiles.

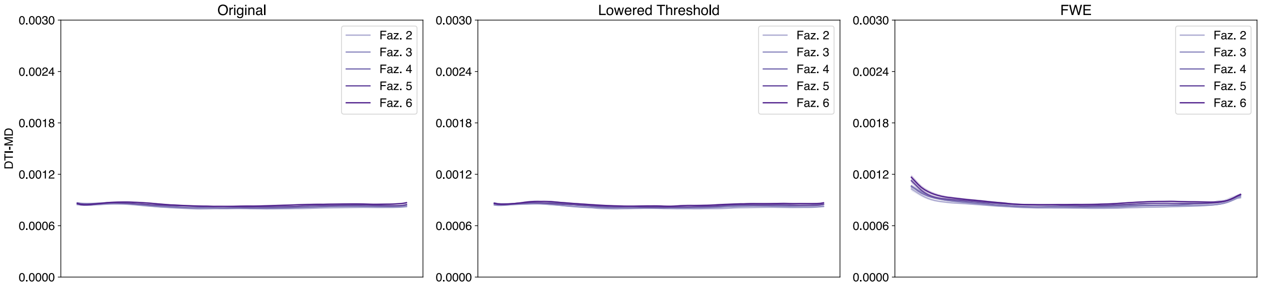


Figure 161: Single-Shell Left Uncinate DTI-MD profiles.

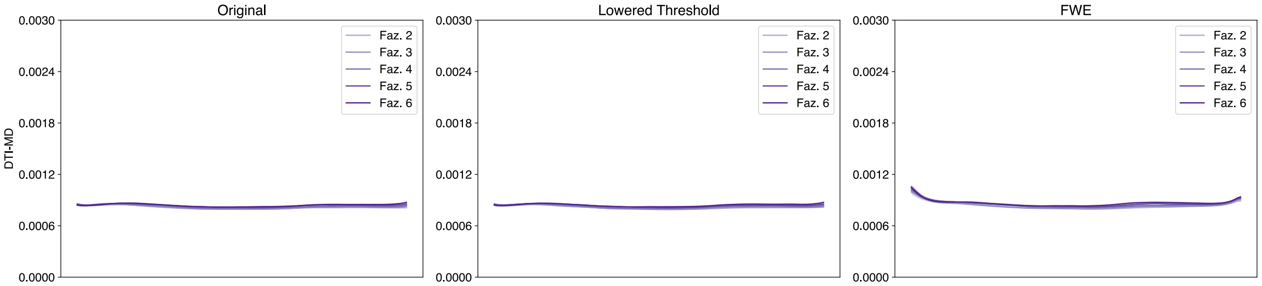


Figure 162: Single-Shell Right Uncinate DTI-MD profiles.

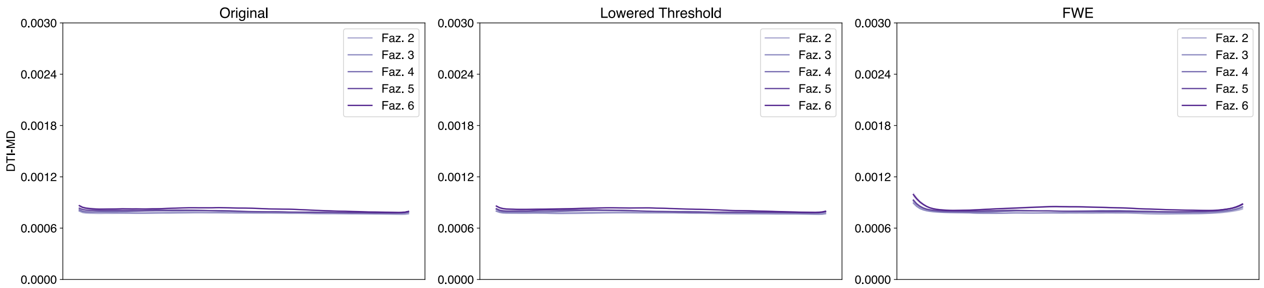


Figure 163: Single-Shell Left Vertical Occipital DTI-MD profiles.

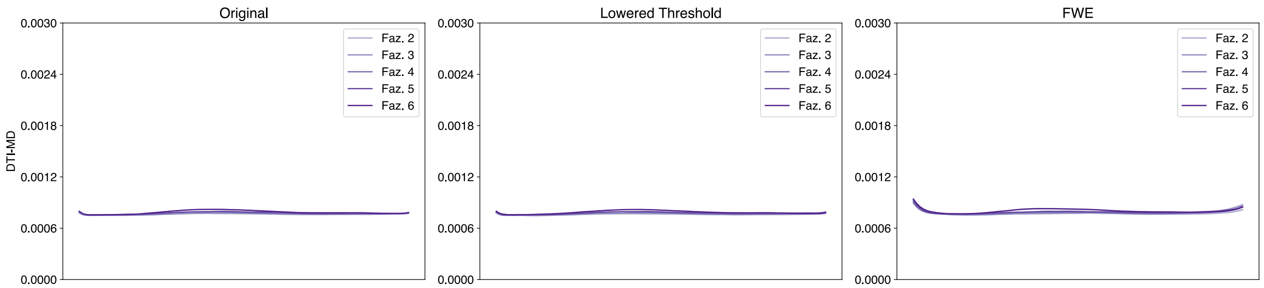


Figure 164: Single-Shell Right Vertical Occipital DTI-MD profiles.

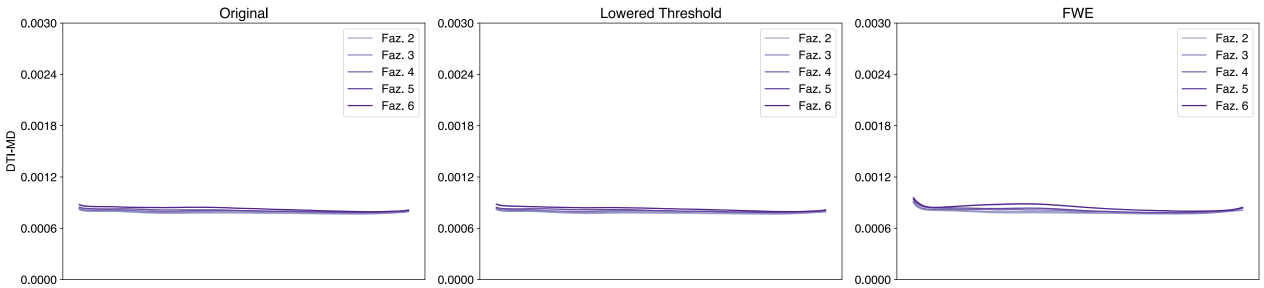


Figure 165: Single-Shell Left Posterior Arcuate DTI-MD profiles.

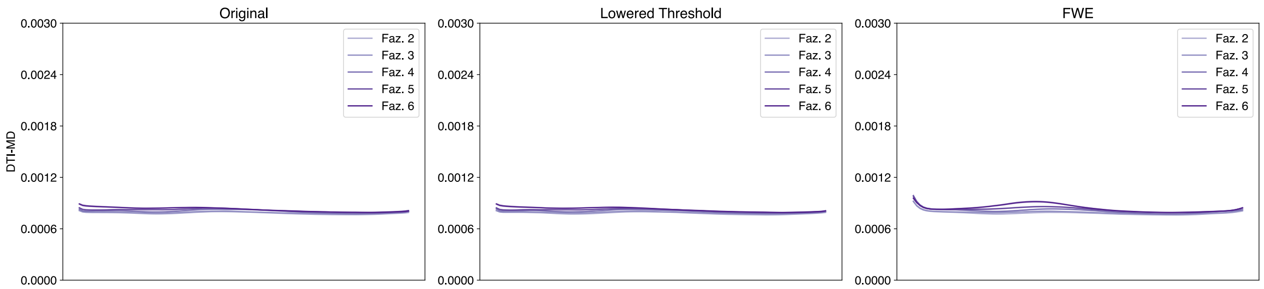


Figure 166: Single-Shell Right Posterior Arcuate DTI-MD profiles.

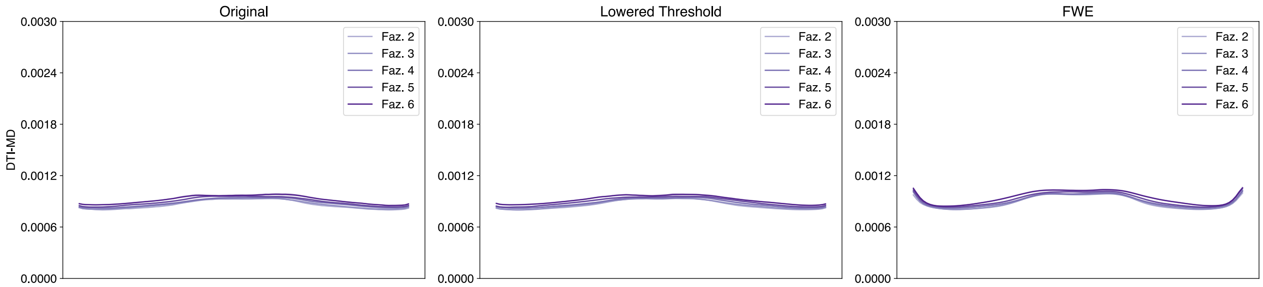


Figure 167: Single-Shell Anterior Frontal Callosum DTI-MD profiles.

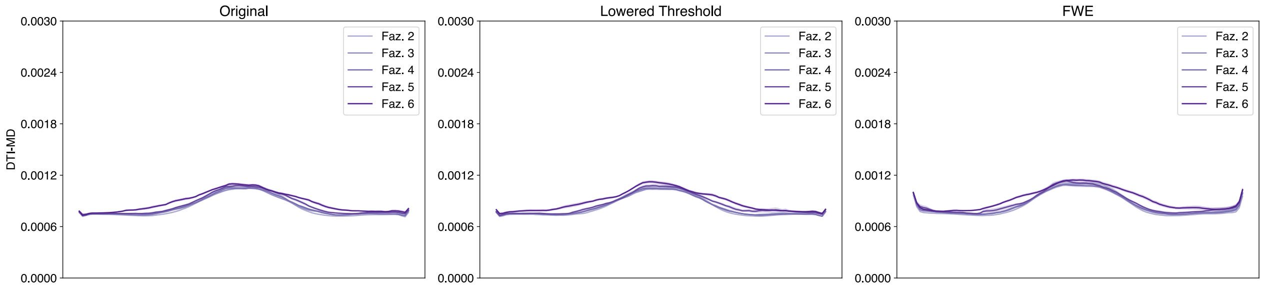


Figure 168: Single-Shell Motor Corpus Callosum DTI-MD profiles.

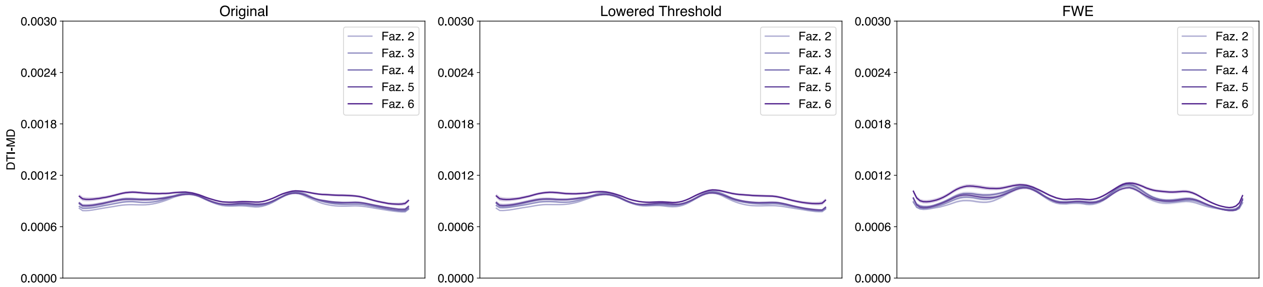


Figure 169: Single-Shell Occipital Corpus Callosum DTI-MD profiles.

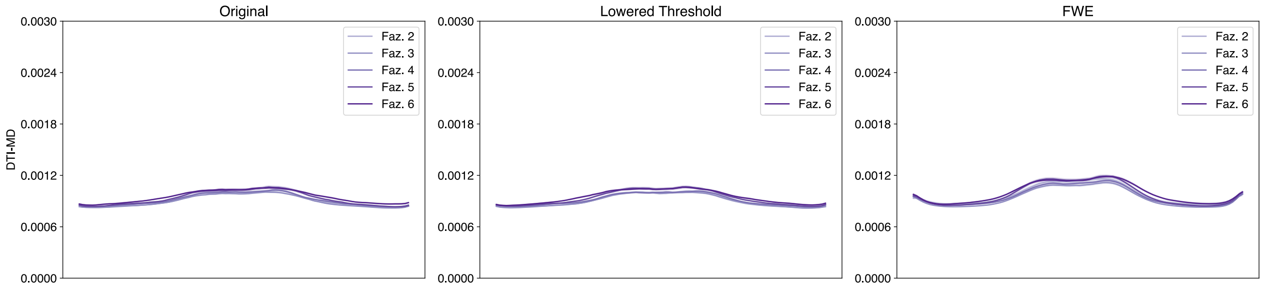


Figure 170: Single-Shell Orbital Corpus Callosum DTI-MD profiles.

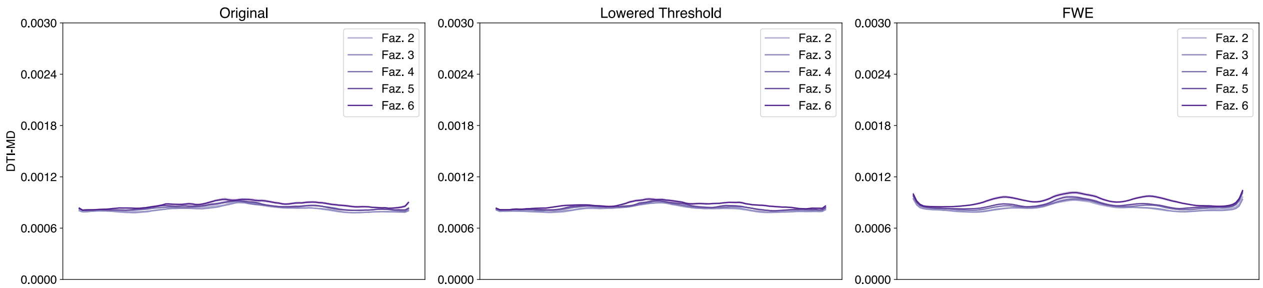


Figure 171: Single-Shell Posterior Parietal Corpus Callosum DTI-MD profiles.

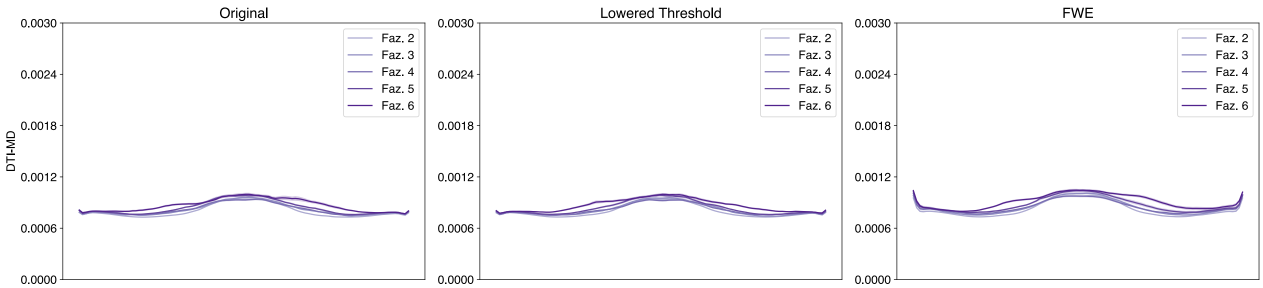


Figure 172: Single-Shell Superior Frontal Callosum DTI-MD profiles.

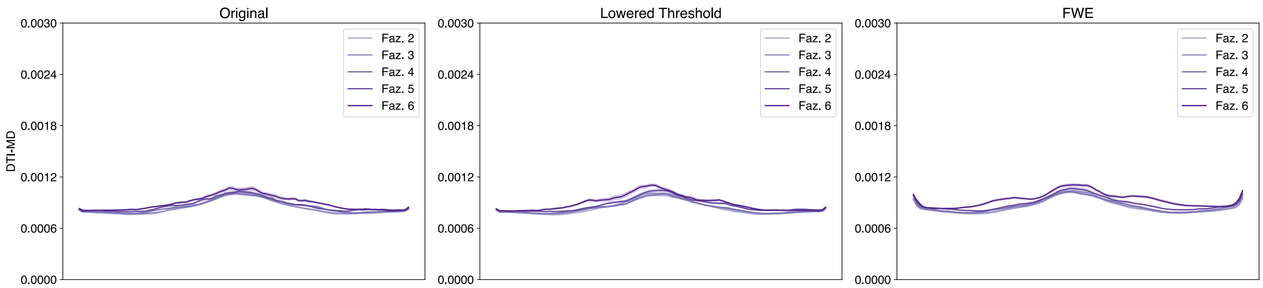


Figure 173: Single-Shell Superior Parietal Corpus Callosum DTI-MD profiles.

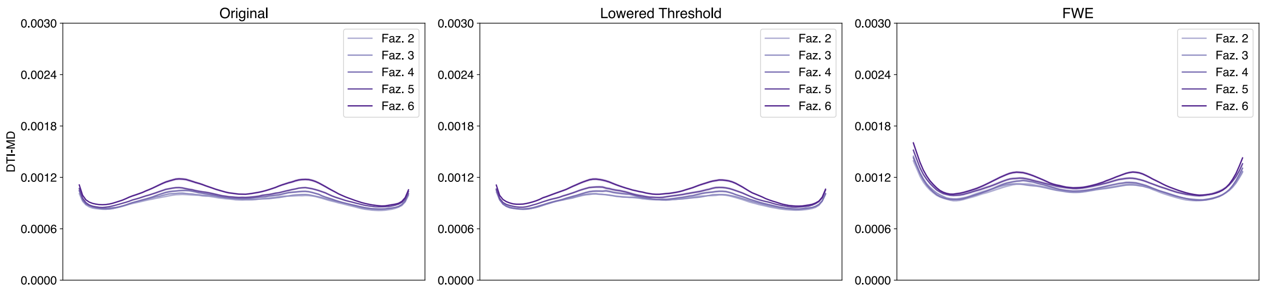


Figure 174: Single-Shell Temporal Corpus Callosum DTI-MD profiles.

### References

Sun, X., & Xu, W. (2014). Fast implementation of DeLong's algorithm for comparing the areas under correlated receiver operating characteristic curves. *IEEE Signal Processing Letters*, *21*(11), 1389–1393. https://doi.org/10.1109/LSP.2014.2337313
